# Guide RNA categorization enables target site choice in Tn7-CRISPR-Cas transposons

**DOI:** 10.1101/2020.07.02.184150

**Authors:** Michael T. Petassi, Shan-Chi Hsieh, Joseph E. Peters

## Abstract

CRISPR-Cas defense systems have been coopted multiple times in nature for guide RNA-directed transposition by Tn7-like elements. Prototypic Tn7 uses dedicated proteins for two targeting pathways, one targeting a neutral and conserved attachment site in the chromosome and a second directing transposition into mobile plasmids facilitating cell-to-cell transfer. We show that Tn7-CRISPR-Cas elements evolved a system of guide RNA categorization to accomplish the same two-pathway lifestyle. Selective regulation of specialized guide RNAs allows long-term memory for access to chromosomal sites upon entry into a new host, while conventional CRISPR features maintain the ability to continually acquire guide RNAs to new plasmid and phage targets. Transposon-encoded guide RNAs are also privatized to be recognized only by the transposon-adapted system working with selective regulation to guard against toxic self-targeting by endogenous CRISPR-Cas defense systems. This information reveals new avenues to engineer guide RNAs for enhanced CRISPR-Cas functionality for genome modification.

## Introduction

CRISPR-Cas systems are widespread in bacteria and archaea providing an efficient defense system from predation from bacteriophages and the burden imposed by other mobile elements (Makarova et al., 2020). CRISPR-Cas systems have also been repurposed in nature for other functions that benefit from programable guide RNA-based sequence recognition (Faure et al., 2019a). One compelling collection of systems involves the occurrence on multiple occasions where a specialized class of transposons called Tn7-like elements have coopted different CRISPR-Cas systems for guide RNA-targeted transposition (Faure et al., 2019b; Peters et al., 2017b). Beyond providing an intriguing example of the propensity of evolution to mix and match useful functions, these systems have also been repurposed in the laboratory as promising tools for genome modification (Klompe et al., 2019a; Strecker et al., 2019). Multiple questions remain with the basic functioning of Tn7-CRISPR-Cas elements, especially with how they interact with other canonical CRISPR-Cas systems. We have discovered mechanisms used by Tn7-CRISPR-Cas elements to categorize guide RNAs, tailoring them for specific functions. This curation of guide RNAs allows regulated pathway choice to recognize highly conserved chromosomal sites while maintaining the ability to continually acquire guide RNAs to new targets.

Tn7-like elements coopted CRISPR-Cas systems on at least four occasions for guide RNA-directed transposition (Peters, 2019). The largest group of Tn7-CRISPR-Cas elements adapted a specific subtype of CRISPR-Cas systems within the type I-F group for guide RNA-directed transposition. Canonical type I-F CRISPR-Cas systems, called type I-F1 systems, use four proteins (Cas8, Cas5, Cas7, and Cas6) as an effector complex called cascade to mature precrRNAs transcribed from a CRISPR array into functional guide RNA complexes. The repeats in the CRISPR array encode the guide RNA handles that are bound by Cas proteins, while target specificity is encoded in the spacers. Cas1 and Cas2 are proteins found across CRISPR-Cas systems for acquiring new spacers (also called adaptation) which are inserted into one end of the array adjacent to a special leader region. Type I CRISPR-Cas systems degrade targets recognized by the system using a helicase-nuclease protein Cas3. The Tn7-CRISPR-Cas elements that derived from the canonical I-F1 systems are called I-F3 systems (Makarova et al., 2020). In the I-F3 Tn7-CRISPR-Cas system the Cas8 and Cas5 proteins are naturally fused. These elements lack the Cas1 Cas2/3 spacer acquisition system found in canonical I-F1 systems, something found true in all four families of independently evolved Tn7-CRISPR-Cas elements. Tn7-CRISPR-Cas elements collect new targeting information *in trans* using functions borrowed from canonical CRISPR-Cas systems. (Peters et al., 2017b). This dependency on canonical CRISPR-Cas systems for acquiring new guide RNAs would also render them vulnerable to the targeted degradation capacity of these same systems.

Prototypic Tn7 and Tn7-like elements are known for the control they have over target site selection. Prototypic Tn7 uses five element-encoded proteins for two pathways that direct transposition into different classes of transposition targets (Waddell and Craig, 1988), a specific neutral chromosomal attachment (*att*) site and mobile genetic elements capable of cell-to-cell transfer. A core machinery used for all transposition involves a heteromeric transposase, TnsA+TnsB, for the breaking and joining functions that underlie transposition that is controlled by a AAA+ regulator protein, TnsC (Bainton et al., 1991; Bainton et al., 1993; Stellwagen and Craig, 1998). TnsABC must function with one of two target site selection proteins, TnsD/TniQ or TnsE, to recognize the different types of transposition targets using distinct mechanisms. Tn7 transposition mediated by the sequence specific DNA binding protein TnsD/TniQ allows high frequency transposition into its *att* site downstream of the *glmS* gene. Because TnsD/TniQ recognizes the coding region of the essential and highly conserved *glmS* gene, the Tn7 TnsD/TniQ *att*-site pathway virtually ensures a place for the transposon to integrate in a new bacterial host (Mitra et al., 2010). Tn7 or diverged families of Tn7-like elements have been identified in 10-20% of sequenced bacteria with homologs of the TnsA, TnsB, TnsC and TnsD/TniQ proteins (Peters et al., 2017b). Tn7-like elements are presumed to have evolved new TnsD/TniQ DNA binding specificities to recognize other coding-regions observed in the expanded collection of *att* sites in different families of Tn7-like elements (Peters, 2019). Prototypic Tn7 will also preferentially recognize mobile plasmids and bacteriophages capable of transfer between bacteria using a second transposition pathway carried out by TnsABC + TnsE (Finn et al., 2007; Wolkow et al., 1996). TnsE recognizes specific features found during DNA replication associated with plasmid mobilization to facilitate transfer of the element between bacteria (Parks et al., 2009; Peters and Craig, 2001; Shi et al., 2015). Unlike TnsD/TniQ, TnsE is not conserved across the other major families of Tn7-like elements and presumably other mechanisms are used to facilitate transfer of the element to new bacterial hosts.

We present a comprehensive bioinformatic analysis of I-F3 Tn7-CRISPR-Cas elements that reveals mechanisms that allowed the evolution of guide RNA-directed transposition involving categorization of guide RNAs. This updated analysis indicates that all I-F3 Tn7-CRISPR-Cas insertion events are explained by guide RNAs encoded in CRISPR arrays within the element. A form of curation allows the I-F3 elements to maintain different classes of guide RNAs to mirror the two-pathway lifestyle found with prototypic Tn7, but with a guide-RNA-only system. Guide RNA-directed transposition into the chromosome occurs via CRISPR arrays that are under the control of a specialized transcriptional regulation system that directs pathway choice or using an atypical CRISPR repeat structure that allows the guide RNA to be private to the Tn7-CRISPR-Cas transposon. The system of guide RNA categorization found in I-F3 Tn7-CRISPR-Cas elements helps explain how they interact with related type I-F CRISPR-Cas systems, such as the ability to tolerate self-targeting guide RNAs that would otherwise allow canonical CRISRP-Cas systems to degrade the host chromosome.

## Results

### I-F3 Tn7-CRISPR-Cas element targeting is explained by spacers in atypical CRISPR array configurations

We conducted an updated bioinformatics analysis of the major family of Tn7-CRISPR-Cas elements which use the I-F3 variant of CRISPR-Cas systems (Experimental procedures) (Peters, 2019; Peters et al., 2017b). An analysis of over 53,000 genomes from gamma proteobacteria identified 801 Tn7-like elements that encode the type I-F3 CRISPR-Cas system found primarily across the orders *Vibrionales* and *Aeromonadales* in two branches (Figure 1). As noted previously one branch, which we refer to as I-F3a (Figure 1), primarily uses attachment sites adjacent to the *yciA* and *guaC* (IMPDH) genes. A second branch, which we refer to as I-F3b, is primarily found in an attachment site downstream of the *ffs* gene encoding the signal recognition particle (Figure 1). Analysis of the elements in the I-F3b branch revealed a subbranch where elements resided in a different attachment site downstream of the *rsmJ* gene (Figure 1). As part of this analysis we reexamined CRISPR arrays and made a striking finding that altered our understanding of how transposition is targeted across all of the I-F3 elements. Previously it was assumed that Tn7-CRISPR-Cas elements evolved to recognize new *att* sites by adapting the sequence-specific DNA binding ability of the TnsD/TniQ protein. However, we can now show that the insertion position of all elements can be explained by guide RNA-directed transposition; for essentially all of the I-F3 elements we can identify a spacer within element-encoded CRISPR arrays that matches a region ∼48 bp from the right end of the element (Figures 1, 2a, and 2b)(Supplementary Table S1). In each of these cases the spacer in the array matches the same protospacer in the *yciA, guaC, ffs*, or *rsmJ* genes (Figure 2b). The specific position of the protospacer that is recognized also appears important. In addition to being at one of the ends of the gene to direct transposition just outside the reading frame, the spacers matching the *yciA, guaC,* and *rsmJ* genes are all found in the same reading frame register that aligns the variable wobble position of the codons with every sixth position in the guide RNA, a position known to flip out and not required to match the protospacer (Fineran et al., 2014; Jackson et al., 2014; Mulepati et al., 2014; Zhao et al., 2014). While it has been previously proposed that guide RNA-directed transposition allowed Tn7-CRISPR-Cas elements to primarily target mobile plasmids to allow movement between bacteria (Peters, 2019; Peters et al., 2017b), we now find that transposition directed into the major chromosomal *att* sites used by the element is also guide RNA-directed.

**Figure 1.**
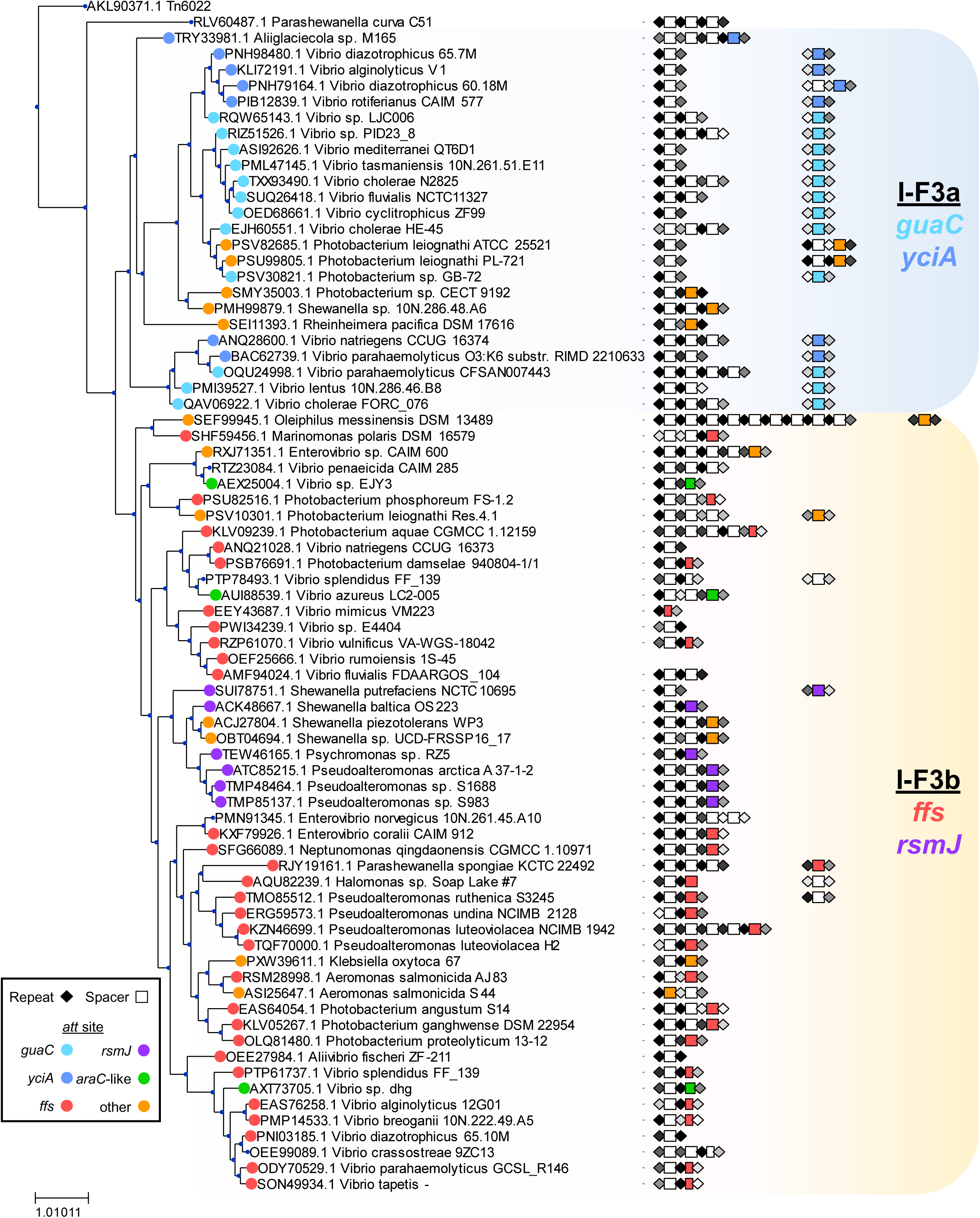
Tn7-like elements with I-F3 CRISPR-Cas systems found in gamma proteobacteria. A TnsA protein similarity tree representing 801 elements indicated by host strain. Elements that were >90 percent identical are indicated with a single representative. A similarity score was calculated for repeats (Experimental procedures) and indicated as black as the highest score and lighter shades of grey as the score decreases. Spacers matching the protospacer found adjacent to the point of insertion are indicated by color. Width of rectangle indicating the spacers on the figure is based on predicted spacer length (Supplementary Table S1). See text for details.

**Figure 2.**
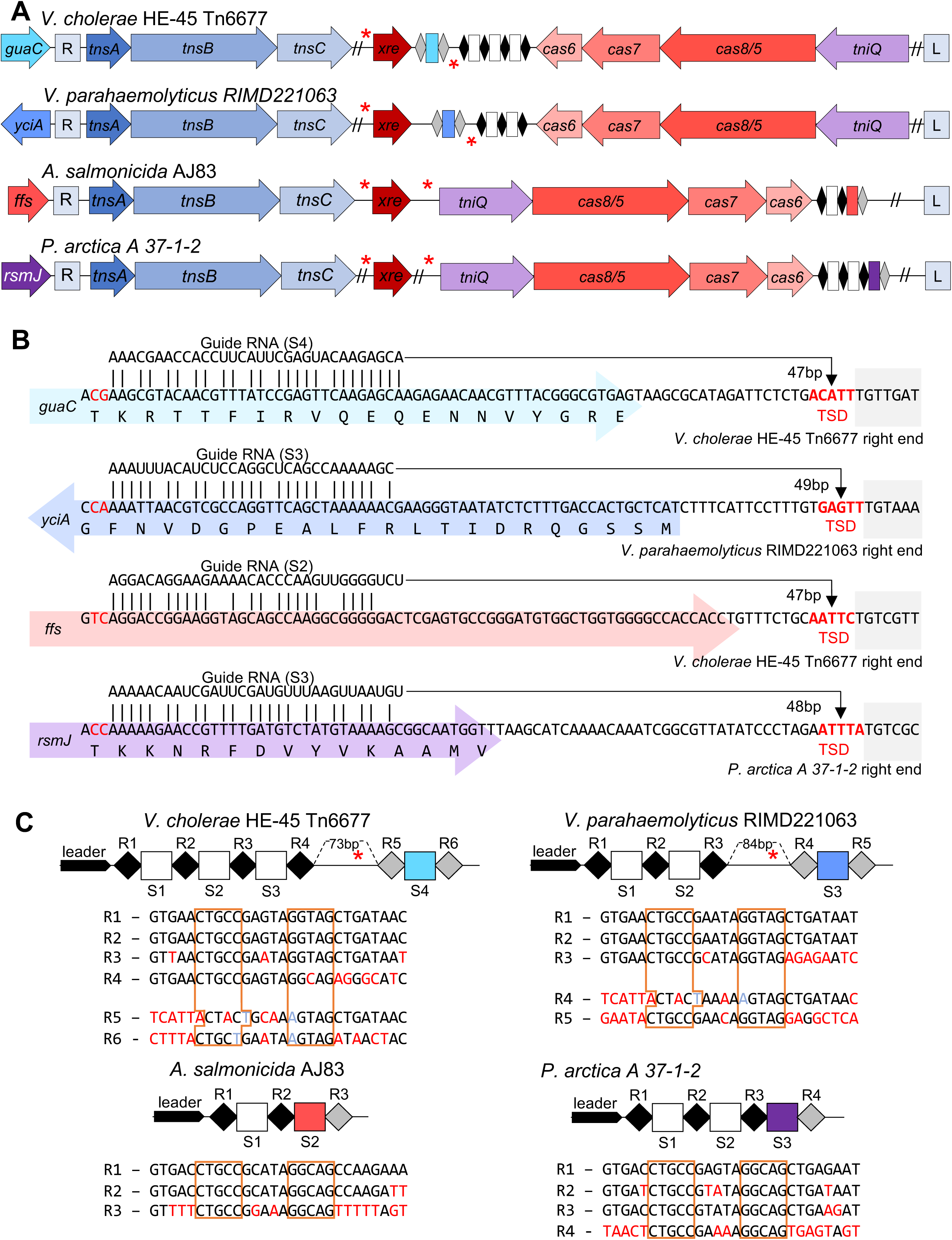
Selected representatives from four *att*-site families of Tn7-like elements with I-F3 CRISPR-Cas systems. Representatives for three major families (*att* sites; *yciA, guaC,* and *ffs*) and one minor family (*rsmJ*) are indicated by host. (a) Transposition genes (*tnsA, tnsB, tnsC,* and *tniQ/tnsD*), Cas genes *cas6, cas7,* and *cas8/5,* and regulator *xre* are indicated. The CRISPR arrays are indicated as in Figure 1. The left (L) and Right (R) ends of the elements are indicated and putative Xre binding sites (red asterisks). (b) Matches between the guide RNA and protospacer are shown in detail with the gene (colored block arrow), right end of the element (grey box), with host indicated. Distance from the protospacer to the target site duplication (TSD, bold red) is shown. (c) CRISPR array is indicated with the leader region and spacer (S#) and repeats (R#) indicated showing the sequence of the repeats. Sequence differences from the first repeat (red), noting changes maintaining the stem (light blue) and the inverted repeats that makes the stem (boxed orange) are indicated. The size of the gap in the array is indicated noting the putative Xre regulatory site.

Of additional interest, the spacers that recognize each one of the four major *att* sites are located in a specific position in the element-encoded CRISPR array(s). There are obvious trends with this position and the configuration of the CRISPR arrays that differed in the two major branches of I-F3 elements. In the branch of I-F3a elements, the spacer that matched the *yciA* and *guaC att* sites was located after a 70-90 bp gap in the array found immediately downstream of the *tniQ, cas8/5, cas7, cas6* operon (Figures 2a and 2c)(see below). In these cases where the CRISPR array was not contiguous it was not clear if the array was transcribed as a single precrRNA or if all of the spacers were capable of being matured into functional guide RNA complexes (addressed below). In the I-F3b branch of elements that recognizes *att* sites associated with the *ffs* and *rsmJ* genes the *att*-site specific spacer tended to be found in the single CRISPR array located downstream of the *tniQ-cas* operon, but always as the last spacer in the array (Figures 1 and 2a).

Around five percent of Tn7-CRISPR-Cas insertions identified in bacterial genomes are not located in one of the four major *att* sites (Supplemental Table S1). However, even with the insertions outside the major *att* sites we could still identify a spacer in the array that was specific to a protospacer ∼48bp from the right end of the element (Figure 1). In all cases, the spacer that directed transposition via a guide RNA into the chromosome was still the last in the array suggesting this placement was significant for the pathway choice with transposition.

Only one transposition event was identified in a plasmid rather than in the chromosome in our analysis. The Tn7-CRISPR-Cas element in *Aeromonas salmonicida* S44 was on a large plasmid in the strain (pS44-1) that was predicted to be a mobile plasmid based on the presence of genes with known roles in conjugal DNA transfer (*tra* genes). Transposition into the site on the plasmid could still be explained by a guide RNA-encoded in the array, however, in this case the spacer was at the leader-proximal position in the array (Supplemental Figure S1a-c). Interestingly, a near-identical Tn7-CRISPR-Cas element found in the *ffs att* site in *Aeromonas hydrophila* AFG_SD03 had a spacer that recognized the same plasmid-encoded gene, but at a different position (Supplemental Figure S1b-c), suggesting a possible plasmid vector important for the dispersal of these elements from *Aeromonas*.

In addition to their distinctive position at the end of or distal to the CRISPR array, the *att* spacers were flanked by repeats with distinctive and novel sequences. New spacers are added to a CRISPR array at the leader-proximal end of the array in a process that duplicates the leader-proximal repeat (Xiao et al., 2017). Therefore, although repeats can diverge over time, the first and second repeats start out identical in CRISPR arrays. In I-F3 Tn7-CRISPR-Cas elements the terminal spacer that was used for guide RNA-directed transposition into the chromosome was invariably flanked by repeats that were highly diverged from the leader-proximal repeat (Figure 2c and Supplemental Table S1). We call the highly diverged repeats, “atypical” repeats, and a guide RNA formed from these sequences atypical guide RNAs. Based on the high divergence from the leader-proximal repeat, it was not clear if the diverged repeats were still functional for guide RNA-directed transposition or if the diversification had rendered them nonfunctional.

### Highly diverged atypical repeat-spacer units can form functional guide RNA complexes

To help understand the unique nature of the CRISPR array structure found in I-F3 Tn7-CRISPR-Cas elements we established guide RNA-directed transposition in a heterologous and genetically tractable system, *E. coli*. Previous studies trying to establish Tn7-CRISPR-Cas transposition in a heterologous host relied heavily on indirect PCR-based techniques to assess transposition, techniques that are vulnerable to artifacts (Rice et al., 2020; Strecker et al., 2020). To get a complete picture of Tn7-CRISPR-Cas transposition, we used an assay that monitored full transposition events. In this assay, candidate transposition targets to be tested were crossed by homologous recombination onto a conjugal F plasmid. Transposition events were monitored by mating the conjugal plasmid into a tester strain after inducing expression of the components of the guide RNA-directed transposition system (Supplemental Figure S2a).

We used informatics to identify Tn7-CRISPR-Cas elements that show clear evidence of robust crRNA-guided transposition, reasoning that such systems would likely show high activity in the laboratory setting. Elements identified in a plasmid in *Aeromonas salmonicida* S44 and in the *ffs* attachment site in *Aeromonas hydrophila* AFG_SD03 were of particular interest because they were near-identical, but found in different species at distinct insertion points suggesting they were functional recently (Supplemental Figure S1a). For the transposition (Tns) and Cas proteins we utilized a coding sequence configuration we predicted to be highly active for transposition by looking for consensus across multiple elements found in *Aeromonas* (Experimental procedrues). To monitor transposition using a modular system, we constructed the *tnsABC*, *tniQ-cas8/5,7,6*, and the CRISPR array in separate expression vectors. A mini Tn7-CRISPR-Cas element was constructed with *cis*-acting transposon end sequences predicted by putative TnsB-binding sites (Peters, 2015) flanking an antibiotic resistance determinant that was situated in the chromosome, the donor site for transposition in our assays.

Initially we analyzed candidate guide RNAs produced from the wild-type configuration of the CRISPR array found in *A. salmonicida* S44. In this configuration the leader-proximal spacer was a perfect match to a mobile plasmid-encoded gene from the native host and the second/terminal spacer had a degenerate match to the *ffs* protospacer with 10 mismatches (Figure 3a, Supplemental Figure S1c). Some of the mismatches between the spacer and the protospacer in the target were at every sixth position and therefore would not impact recognition of the *ffs* guide RNA target (Supplemental Figure S1c). Monitoring transposition following expression of the native array configuration confirmed that functional guide RNAs were produced both from the spacer with the canonical repeat structure at the leader-proximal position and the terminal spacer flanked by highly diverged atypical repeats (Figure 3b). Interestingly, guide RNA-mediated transposition occurred at a higher frequency with the *ffs*-specific spacer even though it contained mismatches and was flanked by atypical repeats (Figure 3b).

**Figure 3.**
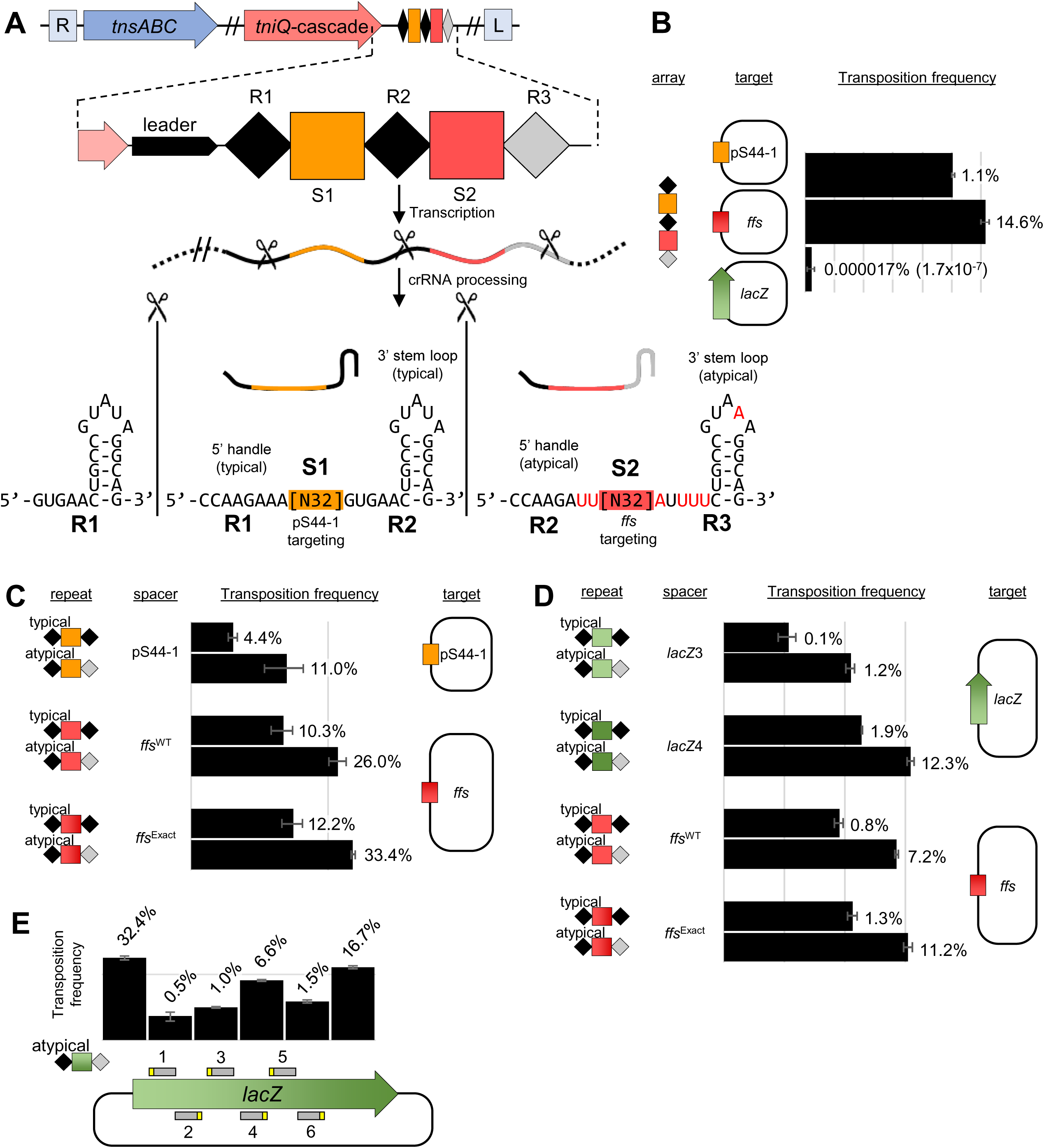
I-F3b element derived from *A. salmonicida* S44 allows RNA-guided transposition with typical and atypical repeats. Various transposition targets were tested using the *A. salmonicidia* S44 native array or as individual repeat-spacer-repeat units. (a) Simplified representation of transposition/CRISPR-associated genes, CRISPR array (marked as in Figure 1) and the resulting typical and atypical guide RNAs with the 5’ and 3’ handles indicated. Position of Cas6 processing is indicated (scissors). (b) Frequency of transposition found with the native *A. salmonicidia* S44 array with targets constructed into an F plasmid; *A. salmonicida* S44 plasmid pS44-1 (pS44-1), chromosomal *ffs att* site (*ffs*) or a negative control, *lacZ* gene. (*lacZ*) (c-e) Transposition frequency found with a single repeat-spacer-repeat unit in various combinations of spacers with typical or atypical repeats from *A. salmonicida* with the indicated targets constructed into an F plasmid. Transposition assay was as described in the Experimental procedures and the average of three experiments is indicated +/- one standard deviation (n=3).

To test the individual contributions of the spacer, protospacer, and repeat sequences, we designed CRISPR array constructs with leader-proximal (typical) or terminal (atypical) flanking repeats as a single guide RNA expression construct and tested various native and synthetic spacer sequences individually. Not only were the guide RNAs with the atypical repeats functional, but also consistently allowed a higher frequency of transposition when compared to the typical repeat when tested with three different spacers (Figure 3c). Additionally the *ffs*-specific spacer showed a higher frequency of transposition than the spacer directed at the plasmid target even though the plasmid spacer had a perfect match to its target and the *ffs* spacer had 10 mismatches to its target (several that were not at the sixth positions that are predicted to be flipped out) (Figure 3c). Altering the native *ffs*-specific spacer so that it was a perfect match to the *ffs* protospacer consistently allowed a modestly higher frequency of transposition (Figure 3c-d).

Guide RNA complexes were also designed using spacers matching different positions in *lacZ* (Figures 3d-e). We found that transposition frequency varied as much as 10-fold with different spacers, even though the sequences recognized all had the same candidate PAM sequence, a result that was not explained by the DNA strand that was targeted in the highly expressed *lacZ* gene (Figures 3d-e). However, regardless of the spacer tested, a modestly higher transposition frequency was consistently found with guide RNAs with the atypical repeats when compared with typical repeats in the Tn7-CRISPR-Cas system from *A. salmonicida* S44 (Figures 3c-d). These experiments confirmed that a functional guide RNA complex could be produced from atypical repeats and hinted that the functionality of these complexes may show important differences from typical repeats. We found that multiple different positions could also be targeted in the *E. coli* chromosome using guide RNA-directed transposition supporting a view that this was not a plasmid-specific process (Supplemental Figure S1d-e).

### Atypical repeats form functionally distinct guide RNA complexes with the Tn7-CRISPR-Cas system from *A. salmonicida* S44

Our experiments suggested that the guide RNA produced from atypical repeats was functional and appeared to allow enhanced transposition activity with the Tn7-CRISPR-Cas system derived from *A. salmonicida* S44. To get a better understanding of the relevance of differences in the repeat sequences, we compared the sequences of the leader-proximal typical repeats with the atypical repeats flanking the terminal spacers to look for common trends across the two branches (Figure 4a). In both branches there were common trends in the final repeat encoding the 3’ handle of the guide RNA with a tendency to lose the typical GTG (positions 1-3), a loss of conservation of the region cleaved from the final guide RNA (positions 21-28), and a general enrichment for adenines in the loop (Figure 4a). Functional differences from changes with the typical and atypical repeats were examined with the *A. salmonicida* S44 element by making changes to the repeat regions encoding the 5’ and 3’ handles of guide RNAs (Figure 4b). While not subject to an extensive analysis, changing the GUG region to an AUU in the 3’ handle of a typical repeat resulted in a significant change in the frequency of guide RNA-directed transposition (Figure 4b). Changing the AUU to a GUG in an atypical array did not show the same significant effect (Figure 4b).

**Figure 4.**
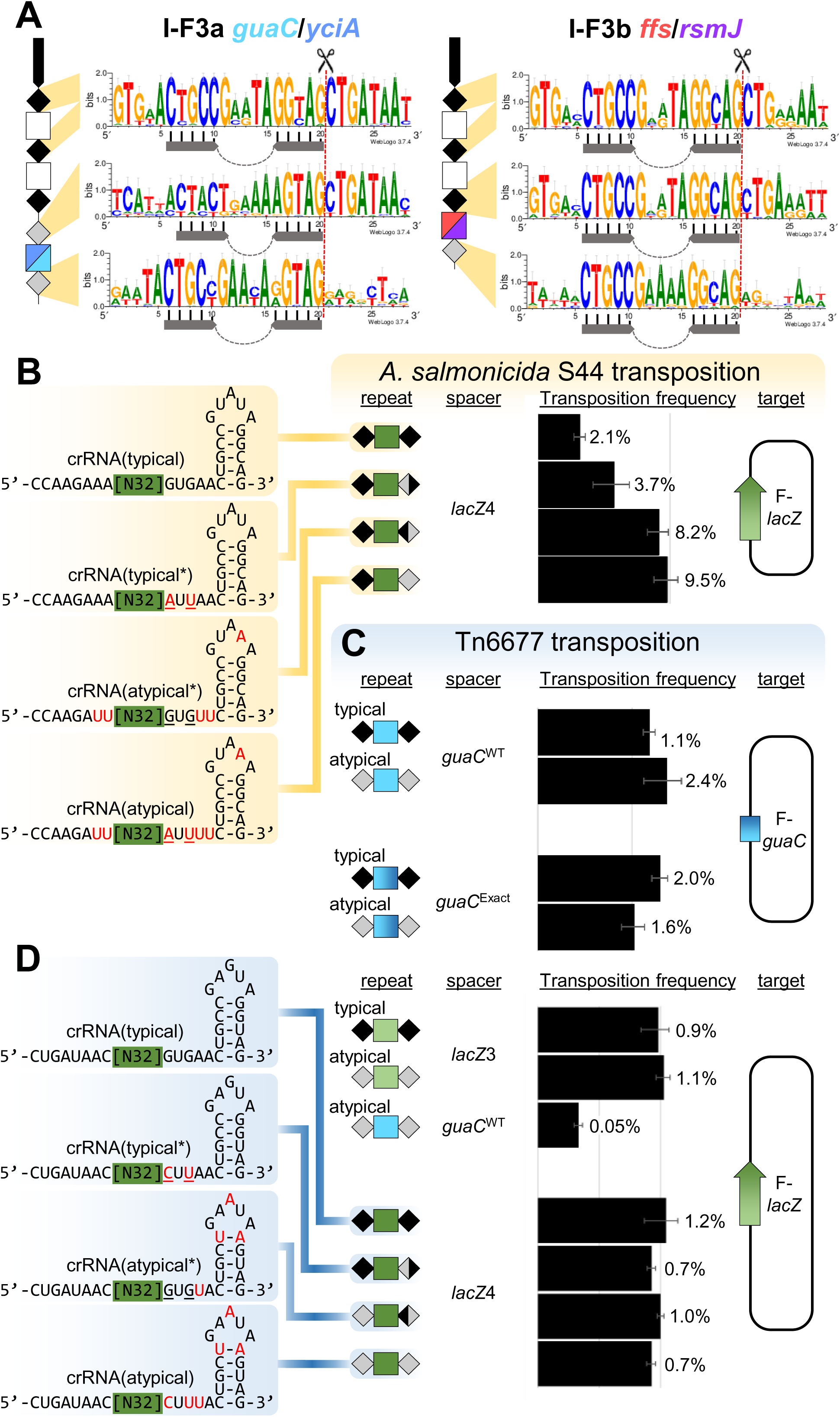
Analysis of atypical repeat sequences. (a) Consensus sequence of the typical and atypical as a function of position in the array. Symbols are as in previous Figures with the stem-loop indicated (top, n=85 for I-F3a, n=74 for I-F3b), (middle, n=51 for I-F3a, n=41 for I-F3b) or (bottom, n=51 for I-F3a, n=41 for I-F3b). (b) Frequency of transposition found with changes in the *A. salmonicida* atypical guide RNAs. (c-d) Frequency of transposition with various typical and atypical repeat combinations from Tn6677 with indicated spacers and their associated targets. Typical guide RNA were tested, comparing with naturally-occurring changes from atypical repeats (red) or engineered mutations (underlined) with the indicated targets constructed into an F plasmid. Transposition assay was as described in the Methods and the average of three experiments is indicated +/- one standard deviation (n=3).

Previous work with a different I-F3 Tn7-CRISPR-Cas systems found with the Tn6677 element from *Vibrio cholerae* HE-45 indicated that guide RNA complexes could be directed to programed target sites in *E. coli* using guide RNAs (Klompe et al., 2019a). The Tn6677 element is in the I-F3a branch of elements and provided a good point of comparison for understanding differences between the two branches of I-F3 Tn7-CRISPR-Cas elements (Figure 1). Tn6677 naturally resides in the *att* site downstream of *guaC* and consistent with the trends we identified above, this element carries the *att* site targeting spacer in a noncontiguous array with an atypical repeat structure (Figure 2). The Tns, Cas, and CRISPR array modules from Tn6677 were constructed under lactose and arabinose expression systems and tested in the transposition assay used for the *A. salmonicida* S44 element above (Supplemental Figure S2a). Like the result found with the *A. salmonicida* S44 transposon, we found that transposition into the native *guaC* attachment site used by Tn6677 required the *guaC*-specific guide RNA encoded in the array and that the atypical repeats in Tn6677 were also functional (Figure 4c). We tested if there was any functional relevance with changes with the typical verses atypical array structure with this element from the I-F3a branch of Tn7-CRISPR-Cas transposons. However, with Tn6677 a similar frequency of transposition was found with the typical and atypical arrays (Figure 4c) or with modest changes in the typical and atypical repeats (Figure 4d). Naturally occurring Tn7- like and Tn7-CRISPR-Cas elements control the left to right orientation with which they insert. Consistent with previous work, Tn6677 seemed to be somewhat relaxed for orientation control (Supplemental Figure S2b). The Tn7-CRISPR-Cas element from *A. salmonicida* S44 showed the expected bias for one orientation found with canonical Tn7 and found naturally when 24 independent insertions were analyzed (Supplemental Figure S2b)(Peters, 2015). For the *A. salmonicida* element, we were able to confirm the target site duplication expected when full transposition occurs and the slight wobble found in previous work with the other I-F3 system (Supplemental Figure S2c)(Klompe et al., 2019a).

### Atypical repeats form private guide RNA complexes with the Tn7-CRISPR-Cas system from *A. salmonicida* S44

A question not previously addressed with Tn7-CRIPSR/Cas systems involves possible cross-talk between CRISPR arrays with other type I-F CRISPR-Cas systems. If the CRISPR array from a Tn7-CRISPR-Cas element with the guide RNAs specific to the chromosome could be used by a standard I-F1 system, the chromosome *att* site would be a target for degradation. This could limit the spread of I-F3 Tn7-CRISPR-Cas elements if it entered a new host that encoded a standard I-F1 CRISPR-Cas system. We investigated if the typical and atypical CRISPR arrays from I-F3 CRISPR-Cas systems could be accessed by a I-F1 system using a transformation-based assay previously established by others (Chowdhury et al., 2017a). In our work testing the *P. aerigunosa* system we co-expressed the Cas proteins and a single spacer CRISPR array with arabinose and T7 expression systems (Vorontsova et al., 2015). Repeats from the type I-F1 system from *P. aerigunosa*, the type I-F3b system from *A. salmonicida*, or the type I-F3a *V. cholerae* Tn6677 were examined using a transformation efficiency assay examining plasmids with and without a protospacer. In control experiments we observed robust interference using the I-F1 CRISPR-Cas system from *P. aeruginosa* PA40 (Figure 5). Transformation was decreased over two orders of magnitude with the plasmid encoding a protospacer (spacer match) compared to a plasmid that lacked the protospacer. Similarly, the typical repeat of the I-F3b system from *A. salmonicida* S44 also allowed robust interference with the plasmid transformation assay. This was not unexpected because the repeats from the canonical I-F1 and I-F3 Tn7-CRISPR-Cas systems are similar (Supplemental Figure S3) and it is likely that the I-F3 Tn7-CRISPR-Cas systems rely on standard I-F1 systems for spacer acquisition (see Discussion). A different result was found when the spacer was situated with atypical repeats from the I-F3b system found with the *A. salmonicida* Tn7-CRISPR-Cas system; guide RNA complexes formed with atypical repeats were not functional for interference in the plasmid transformation assay (Figure 5). The compromised use with the atypical repeats for interference was in contrast to the enhanced use we found for guide RNA-directed transposition with the I-F3b system from *A. salmonicida* (Figures 3 and 4). This result is of particular interest because it provides a mechanism that would allow chromosomal-targeting spacers to be tolerated in hosts with standard I-F1 CRISPR-Cas systems by allowing them to remain private to the I-F3b system. This privatization was absent in the Tn6677 I-F3a system from *V. cholerae*. With the I-F3a *Vc* system, robust interference was found with either the typical repeats or the highly diverged atypical array from this element. The results below suggest that I-F3a Tn7-CRISPR-Cas elements instead use a separate transcription network to tolerate self-targeting spacers (see below).

**Figure 5.**
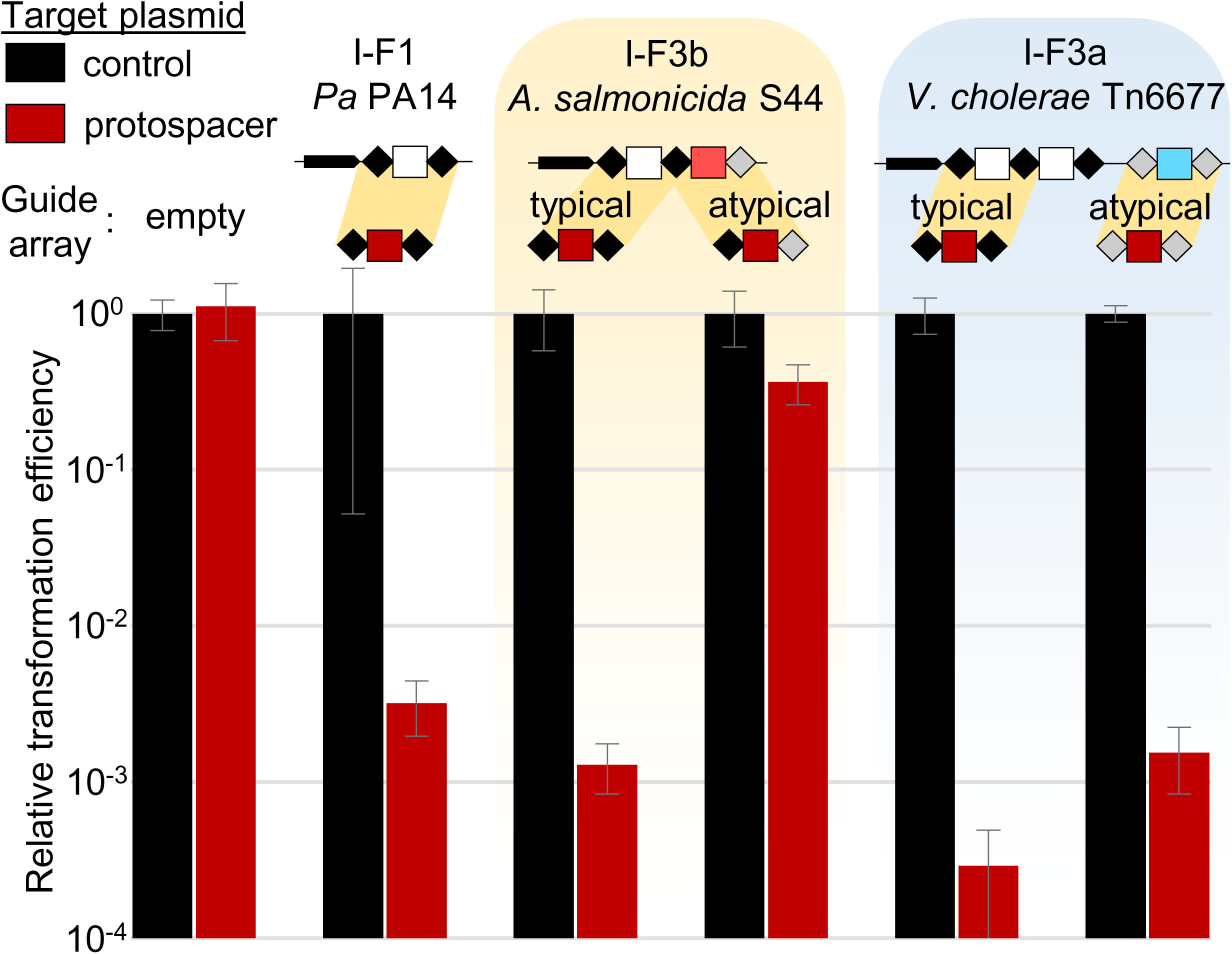
*P. aeruginosa* type I-F1 Cascade can utilize heterologous I-F3 CRISPR arrays in a plasmid interference assay, but I-F3b atypical guide RNAs are privatized. Expression of *P. aeruginosa* protein components with various arrays reduces transformation efficiency for protospacer containing plasmid (*ffs*), but not control (empty). Single unit arrays from *P. aeruginosa* PA14, *A. salmonicida* S44, and *V. cholerae* HE-45 Tn6677 were tested in various typical and atypical repeat configurations as indicated. Repeat configuration sequences are presented in Supplemental Figure S3b. All data indicate mean +- standard deviation of three biological replicates.

### I-F3 elements utilize XRE-family transcriptional regulators to regulate CRISPR-Cas components

To better understand I-F3 Tn7-CRISPR-Cas element dissemination, we searched for genes conserved among diverse members of this group. One of the other genes found conserved across I-F3 Tn7-CRISPR-Cas elements were predicted XRE-family transcriptional regulators. The *xre* gene resides at a conserved position between the *tnsABC* and *tniQ-cas8/5,7,6* operons in nearly all I-F3 elements (Figure 2a). While each of the two branches of I-F3 elements have *xre* genes, the predicted regulatory gene in each branch segregated with phylogenetically distinct families of controller (C) proteins associated with restriction-modification systems. I-F3a elements have a 68 amino acid Xre protein related to C.*Ahd*I and I-F3b elements have a ∼100 amino acid Xre protein related to C.*Csp*231I (Supplemental Figure S4a). Candidate regulatory features could also be identified with the *tniQ-cas* and CRISPR arrays based on homology with the previously established systems (Supplemental Figure S4b, see below)(Streeter et al., 2004).

We analyzed putative promoter regions in I-F3a elements and discovered candidate sites for Xre-mediated regulation upstream of *xre* as well as directly upstream of the *att*-targeting spacer in Tn6677 and other members of this branch of elements (Figure 6a). To confirm physical interaction of element bound Xre regulators with these potential binding motifs, we purified Xre from two elements in the I-F3a branch, *V. cholerae* HE-45 Tn6677 (Vc) and *V. parahaemolyticus* RIMD221063 (Vp). Both Xre proteins showed sequence-specific interaction with the candidate promoter sequences upstream of *xre* and upstream of the *att*-targeting spacer as demonstrated by electromobility shift assays (Figure 6c). A functional role for this interaction was shown by a LacZ reporter assay testing expression from promoters pXre or pArray2 for both I-F3a elements tested in the absence of Xre or with varying expression levels using a titratable pBAD promoter from a plasmid. Xre was found to autoregulate its own pXre promoter, which allowed minimal transcription without Xre, was activated by low amounts of Xre, and repressed as the expression of Xre increased (Figure 6e). Meanwhile the pArray2 promoter identified for the *att*-targeting spacer was highly expressed when Xre was not present and increasingly repressed with increasing amounts of Xre induction (Figure 6e). As explained in greater detail in the Discussion, the regulation we observed would allow a burst of the atypical guide RNA that is specific to the *guaC* or *yciA att* sites with I-F3b elements upon entry into a new host via zygotic induction with a dual regulatory role as an inducer at low Xre concentration and a strong repressor at higher concentrations.

**Figure 6.**
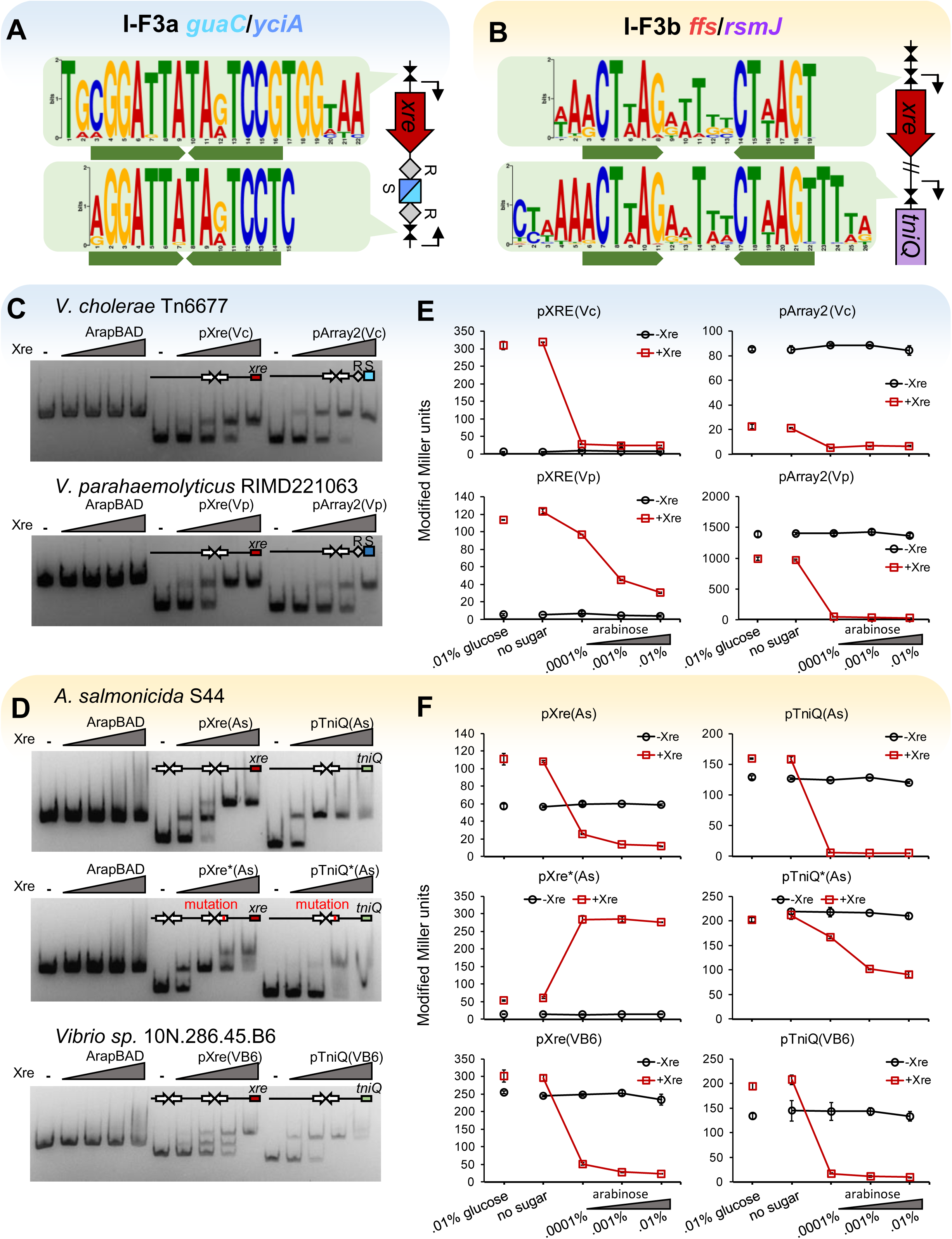
Xre proteins regulate components of RNA-guided transposition. (a-b) Consensus sequence for putative Xre binding motifs in I-F3a and I-Fb elements. (c-d) Xre-binding resolved by EMSA. DNA fragments with the transcription control regions were incubated with increasing amounts of Xre protein from the respective element before electrophoresis (100 nM DNA; protein:DNA ratios = 0,2,5,10,20:1). (e-f) Promoter function resolved by LacZ expression monitored by Miller units at various arabinose controlled Xre expression levels. Transcription control regions are indicated from *V. cholerae* HE-45 Tn6677 (Vc), *V. parahaemolyticus* RIMD221063 (Vp). *A. salmonicida* S44 (As)*, and Vibrio sp.* 10N.286.45.B6 (VB6).

I-F3b elements were also surveyed for inverted repeat motifs to investigate the functional role of the conserved C*.Csp*231I-like XRE regulator. Like I-F3a elements, conserved motifs were found in the promoter region of *xre* that were nearly identical to those used by C.*Csp*231I (Figure 6b, Supplemental Figure S4b)(McGeehan et al., 2011). Unlike the I-F3a elements, the conserved motif could not be identified upstream of the CRISPR arrays with the I-F3b elements, and instead we found a single copy of this motif upstream of the *tniQ-cas8/5,7,6* operon (Figures 2a and 6b). To confirm a role for these motifs in regulation, we selected two I-F3b elements from *A. salmonicida* S44 (As) and *Vibrio sp.* 10N.286.45.B6 (VB6), and confirmed binding to these sites with EMSA and purified Xre regulator. Binding to the two predicted motifs in the upstream region of Xre could be visualized as two separately migrating species (Figure 6d). Mutating the *xre*-proximal regulatory motif weakened interaction as demonstrated by the higher concentration of protein required to achieve a full mobility shift (Figure 6d). We additionally visualized interaction with the motif upstream of *tniQ-cas8/5,7,6* and confirmed the sequence-specific nature of binding by utilizing a mutated motif which weakened interaction. LacZ reporter assays were again used to confirm a functional role in regulation. Xre regulator was shown to act as a repressor of its own pXre promoter (Figure 6f). Interestingly, mutation of the proximal binding site which impaired binding *in vitro* resulted in Xre regulator instead acting as an activator, suggesting interaction with the distal site activates transcription while interaction with the proximal site represses it (Figure 6f). Similar to the result with the I-F3a elements, we found that the XRE regulator was able to repress *tniQ-cas8/5,7,6* expression and this repression was impaired by mutation of the conserved binding motif (Figure 6f). As elaborated on in the discussion, Xre regulation in the I-F3b systems would allow high expression of the TniQ-Cascade guide RNA system when the element entered a new host, providing a mechanism for encouraging transposition into a conserved host chromosome *att* site.

## Discussion

Our findings have revealed that the major family of Tn7-CRISPR-Cas systems have evolved to be dedicated for guide RNA-directed transposition. We find that all transposition can be accounted for by spacers in the element-encoded CRISPR array (Figure 1) and that transposition with members of both major branches of these elements require guide RNAs for *att*-targeted transposition (Figures 3 and 4). We also show that the spacers used to target chromosomal sites show certain characteristics; they are flanked by repeats that are highly diverged from those normally used for the process of spacer acquisition and reside at the last position in a CRISPR array (Figures 1 and 2). The repeat divergence appears to be specifically beneficial for a type I-F3b element from *A. salmonicida* S44, these atypical guide RNA complexes are unusable for interference by a canonical I-F1 system (Figure 5), while allowing for a higher level of transposition than found with the typical guide RNAs (Figure 3). We show that each of the major branches of I-F3 Tn7-CRISPR-Cas systems have separately coopted XRE family regulators (Supplementary Figure S4). XRE family regulators are common in mobile elements allowing a coordinated response upon entry into a new host. In the adaption of this system to I-F3 Tn7-CRISPR-Cas elements the regulatory features will result in a burst of expression specifically favoring guide RNAs directing transposition into chromosomal *att* sites (Figure 6). Regulation of the functional diverged guide RNAs in I-F3 elements allows a private transcription network that is likely important for I-F3 elements to tolerate self-targeting spacers.

### Atypical arrays can form private guide RNA complexes in the I-F3b system from *A. salmonicida* S44

A significant consequence of I-F3 Tn7-CRISPR-Cas elements maintaining a guide RNA-only directed lifestyle is the hazard of canonical I-F1 CRISPR-Cas systems using the self-targeting guide RNAs that are encoded in the element. Self-targeting spacers that are accessed by other canonical CRISPR-Cas systems would be a liability whenever the element entered a host that already had this family of CRISPR-Cas systems. To this point, we find that a I-F1 system can use typical guide RNAs encoded in I-F3 CRISPR arrays in interference (Figure 5). Standard I-F1 systems are found in *Vibrionales* and *Aeromonadales* and with our analysis we found multiple specific examples where I-F3 Tn7-CRISPR-Cas elements reside in strains that have standard I-F1 systems (Supplemental Table S1). Self-targeting spacers have been identified in the work of others and have even been used as a tool to help identify anti-CRISPR systems (Borges et al., 2017). However, established members of anti-CRISPR systems are not widely conserved in elements with Tn7-CRISPR-Cas systems in our analysis (Wiegand et al., 2020).

In the case of the I-F3b system from *A. salmonicida* S44, the atypical repeat appears to be a specific adaptation that allows a higher frequency of guide RNA targeted transposition (Figures 3 and 4) and privatization from a canonical I-F1 interference system (Figure 5). It is unresolved from our work how spacers in the I-F3 CRISPR arrays can be private from the I-F1 system, but could result from poor processing by the canonical system. A better understanding of what allows atypical guide RNAs to function differently in the I-F3b element derived from *A. salmonicida* may offer avenues for optimizing guide RNA-directed transposition in future work by modifying the repeats. Interestingly, the type I-F3a Tn7-CRISPR-Cas system from Tn6677 did not show enhanced transposition with the atypical array found in this system; the frequency of transposition was the same with the typical and atypical repeat, even though it was highly diverged (Figures 4c and 4d). The highly diverged repeats from Tn6677 were also readily used by a canonical I-F1 interference system (Figure 5), suggesting that other mechanisms are required for tolerating self-targeting arrays in this element (see below). Highly diverged repeats around a certain spacer in arrays would provide a benefit by helping to protect against any method of spacer loss that depends on homology.

We suspect that there are multiple distinct ways that I-F3 Cas proteins have adapted to recognize specialized atypical repeats based on our bioinformatics. Interestingly, we notice that in one subbranch within the I-F3b elements the final spacer is reduced by 10 to 12 base pairs in length (Figure 1 and Supplemental Figure S5). These smaller spacers produce functional guide RNAs as predicted by the commensurate natural repositioning of the insertion closer to the protospacer (Supplemental Figure S5). Previous work in closely related CRISPR-Cas systems suggests that guide RNAs of this length will not be functional for targeting transposition nor robust interference (Klompe et al., 2019a; Kuznedelov et al., 2016). However, the ability of the I-F3b systems to accommodate shorter guide RNAs could provide another mechanism of privatization from other I-F CRISPR-Cas systems. Intriguingly, naturally occurring minimal type I-F2 CRISPR-Cas systems tested in the laboratory are not functional for interference with similarly truncated guide RNAs, but can still form complexes capable of forming R-loops to matching protospacers (Gleditzsch et al., 2016).

### XRE family proteins were coopted in different ways in the two branches of I-F3 CRISPR-Cas elements

XRE family regulators are common on mobile elements allowing a coordinated regulatory program to facilitate adaptation upon entry into a new host. These systems take advantage of “zygotic” induction where the gene program is reset when they enter a new host that lack the Xre protein. This type of regulatory scheme was investigated in restriction-modification systems which utilize XRE family regulator C proteins to control temporal regulation of methylation and restriction (Rodic et al., 2017). When the restriction-modification system enters a naïve host on a mobile element, methylation of host DNA must precede endonuclease expression to prevent self-degradation. Methylation activity must remain at a modest level to methylate newly replicated host DNA without protecting subsequent invading DNA elements while the endonuclease must be expressed to defend against other mobile elements. Based on previous work with related C.*Ahd*I (Bogdanova et al., 2008; McGeehan et al., 2006) and C.*Csp*231I (Rezulak et al., 2016; Shevtsov et al., 2015) proteins and our *in vivo* an *in vitro* analysis of the promoters and Xre regulators found in I-F3 systems, we can make predictions to suggest why these are core conserved features in Tn7-CRISPR-Cas elements.

Both branches of I-F3 elements likely benefit from high expression of Xre-controlled promoters when they enter a new host that lacks the repressor protein. Using the I-F3a elements tested in our experiments as an example, the *att*-targeting spacer, *guaC* or *yicA*, becomes derepressed upon entering a new host. Its high expression level would program the Tn7-CRISPR-Cas element to transpose into the neutral chromosomal *att* sites. As the Xre repressor increases in concentration the promoter expressing the *att*-targeting spacer is predicted to be shut off based on the tight repression found in our experiments even under the lowest induction level (Figure 6e). Xre regulation, which includes activation at low concentrations (Figure 6e), is predicted to delay repression of *att*-targeting spacer to allow transposition to occur while also, presumably, ensuring a stable Xre concentration after transposition. The ability to categorize guide RNAs with differential expression from two arrays would allow high expression on entry, but repression upon establishment of the element, thereby helping guard against potential self-targeting with this guide RNA in the presence of endogenous I-F1 systems. Presumably a basal expression level from the primary CRISPR array would allow the production of sufficient guide RNA complexes to recognize mobile elements to allow horizontal transfer of I-F3a elements into new hosts. Careful controlled expression may be especially important in I-F3a elements, because both typical and atypical could be used for interference by a I-F1 system (Figure 5).

In the I-F3b elements, the TniQ-Cascade operon itself is subject to Xre regulation. Upon entry into a new host, high expression of the I-F3b machinery is expected until sufficient repressor protein accumulates (Figure 6). Presumably this bolus of expression would facilitate transposition into the chromosomal *ffs* or *rsmJ att* site. In CRISPR-Cas systems, the total number of effective guide RNA complexes will be a function of the amount of cascade components as related to the number of different spacers and their expression levels (Martynov et al., 2017). Therefore, the I-F3b systems like the ones found with *A. salamonicida* S44 may draw additional benefit by swamping the cell with effector complexes to overwhelm any I-F1 systems potentially found in a new host. While I-F3b elements do not rely on transcriptional regulation to privatize the self-targeting spacer from any endogenous CRISPR-Cas system found in a new host, they have nonetheless coopted a similar regulatory pathway to activate transposition function in a new host.

### The two-pathway lifestyle is maintained in diverse Tn7-like elements using different mechanisms

The prototypic Tn7 element maintains two transposition pathways which are directed by two dedicated target site selection protein (Peters, 2014). We find that the type I-F3 Tn7-CRISPR-Cas elements use a form of spacer categorization involving repeat diversification and transcriptional regulation as mechanisms to establish the same two-pathway lifestyle.

The outlier element found in *Parashewanella curva* C51 provides an interesting alternative hybrid two-pathway strategy from the other elements using the I-F3 CRISPR-Cas system (Figure 1). Unlike the other I-F3 family elements indicated in Figure 1, this element does not have a spacer that could form a guide RNA complex to recognize where it inserted in the chromosome. Interestingly, this element is from a small group of representatives that encode two TniQ proteins. One TniQ is associated with the Cascade operon and the second is similar to a TniQ protein found in other Tn7-like elements lacking CRISPR-Cas components that reside in an attachment site downstream of the *parE* gene encoding the essential and conserved toposiomerse IV, subunit B (Supplemental Figure S6). These elements are predicted to use guide RNA-directed transposition to recognize mobile elements, but presumably continue to rely on the mechanism used by prototypic Tn7 and most other Tn7-like elements (Peters, 2014), in this case with the TniQ protein recognizing the *parE* gene coding region. In two previously identified instances where type I-B CRISPR-Cas systems were captured by other Tn7-like members the elements have a similar configuration with two *tniQ* genes and likely function in a similar way to allow a two pathway life-style (Peters et al., 2017b). It is unclear how Tn7-CRISPR-Cas elements with a type V-K CRISPR-Cas systems control the use of two pathways given these elements only encode a single *tniQ* gene (Faure et al., 2019b).

The guide RNA complexes used by the major *att* pathways recognizing the *guaC, yciA, ffs,* and *rsmJ* have certain characteristics that favor their maintenance and utility across bacterial species. In addition to recognizing a site close to the end of a gene the specific sequence recognized also always occurs in a specific register with the reading frame where the sixth position aligns with the wobble position of codons (Figure 2b). Given that every sixth position in the guide RNA does not make base pair contacts with protospacer DNA, this placement of the guide RNA provides a partial safeguard to differences in protospacers that can be encountered from codon bias in the genes that are recognized for the major attachment sites. While presumably any guide RNA would benefit from the ability to accommodate the wobble position, this is more important for the lifestyle of Tn7-CRISPR-Cas elements that rely on identifying a safe integration site whenever they enter a new host and one that must serve as a form of long-term memory. Similar considerations may be important in other cases where CRISPR-Cas has been coopted for alternative activities involving self-targeting guide RNAs (Faure et al., 2019a). Presumably there is a barrier to initially obtaining and fixing these new spacers because we show that spacers with typical repeats can be accessed by a canonical I-F1 system (Figure 5). This may help explain why we only found one additional example outside of the *guaC, yicA, ffs* and *rsmJ*, sites where the same protospacer was recognized in more than one Tn7-CRISPR-Cas element; a spacer that recognizes a AraC-family protein of unknown function was found four times (Supplementary Figure S5b and Supplemental Table S1).

It will be interesting to learn how the adaptations found in I-F3 Tn7-CRISPR-Cas systems extend to canonical CRISPR-Cas systems and in other cases where CRISPR-Cas systems have been coopted for new functions. The examples shown here join numerous others where it is now clear that CRISPR-Cas activity is regulated to offset potential negative impacts of these potent systems (Hoyland-Kroghsbo et al., 2017; Patterson et al., 2016; Westra et al., 2010). Transposition levels with the I-F3b system developed in our work is robust, occurring in as much as one third of the cell population, suggesting that it should be a good candidate for future application-based work. The new information with atypical guide RNAs also suggests additional avenues for altering guide RNA activity for practical applications in Tn7-CRISPR-Cas systems or potentially CRISPR-Cas systems in general.

## Acknowledgements

This work was supported by National Institutes of Health Grant R01 GM129118. Shan-Chi Hsieh was supported by a training fellowship from the Taiwan government. We thank Nancy Craig and Ailong Ke for comments on the manuscript. Cornell University has filed patent applications related to this work with JEP as inventor. We are grateful for reagents provided by John Cronan, Tobias Doerr, Elizabeth Raleigh, Ailong Ke, Addgene, and the *E. coli* Genetic Stock Center.

## Experimental Procedures

### Bacterial Strain and Plasmids

Strains and plasmids used in this study are listed in Tables 1 and 2. Primers and synthesized gene fragments are listed in Tables 3 and 4. Standard molecular cloning techniques were used to make the vectors described below using vendor instructions.

**Table 1.**
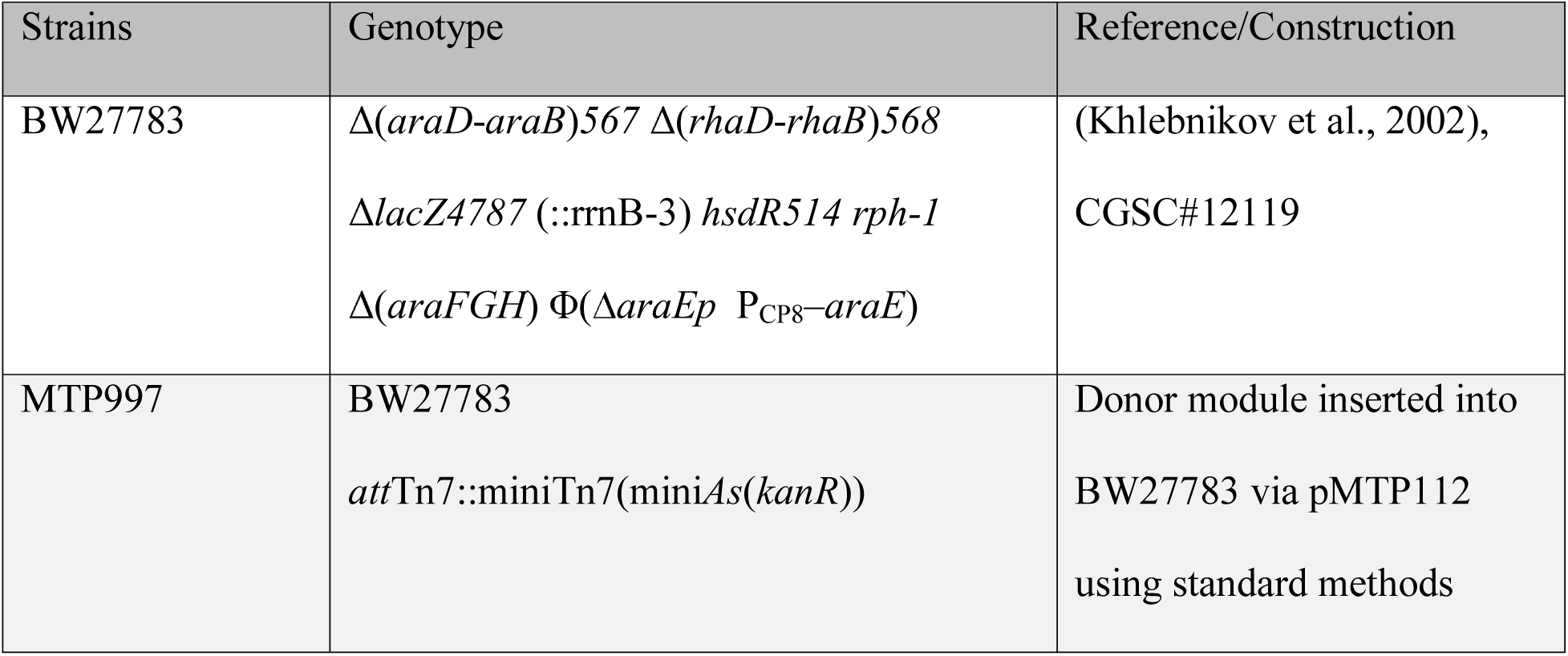
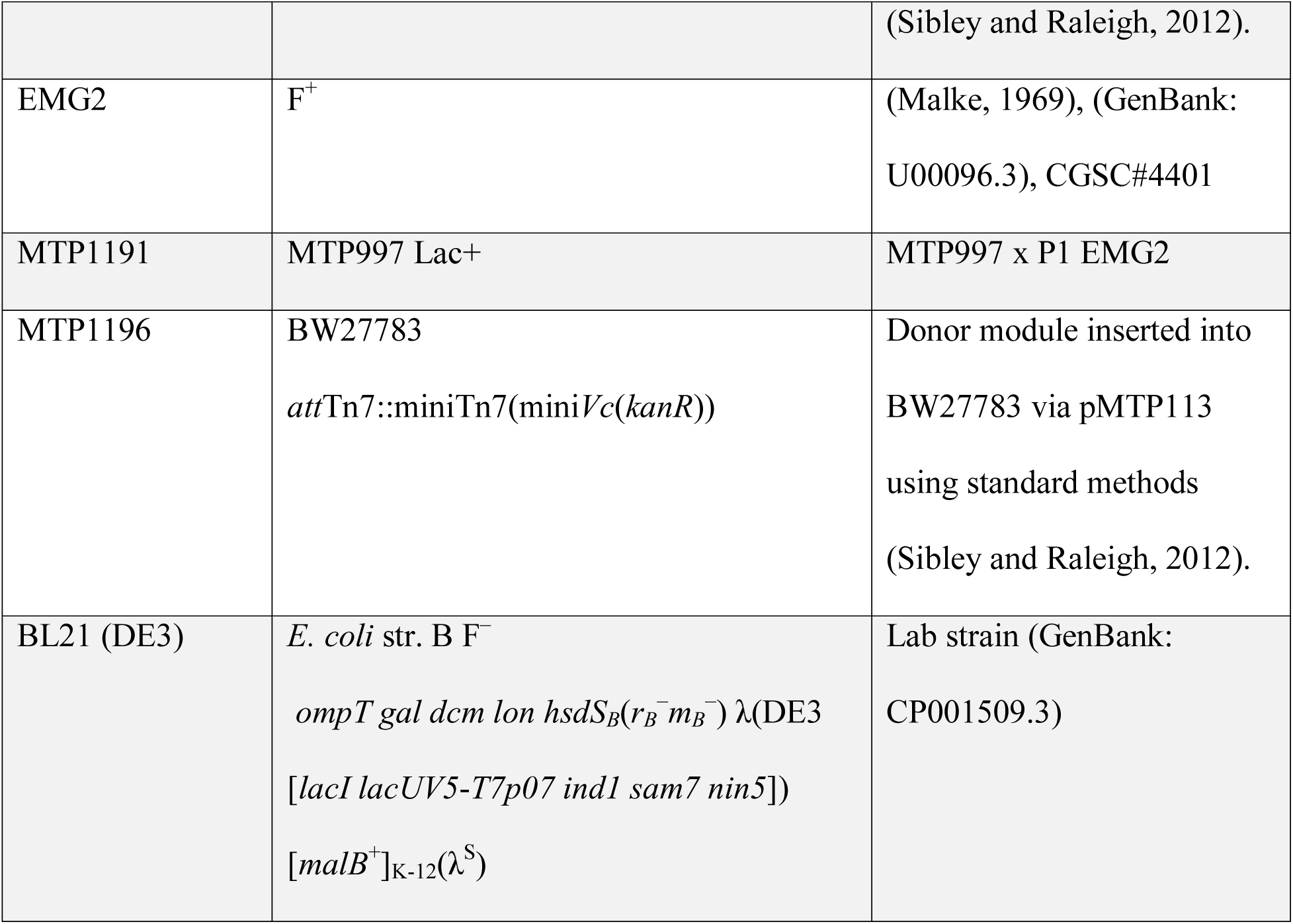

**Table 2.**
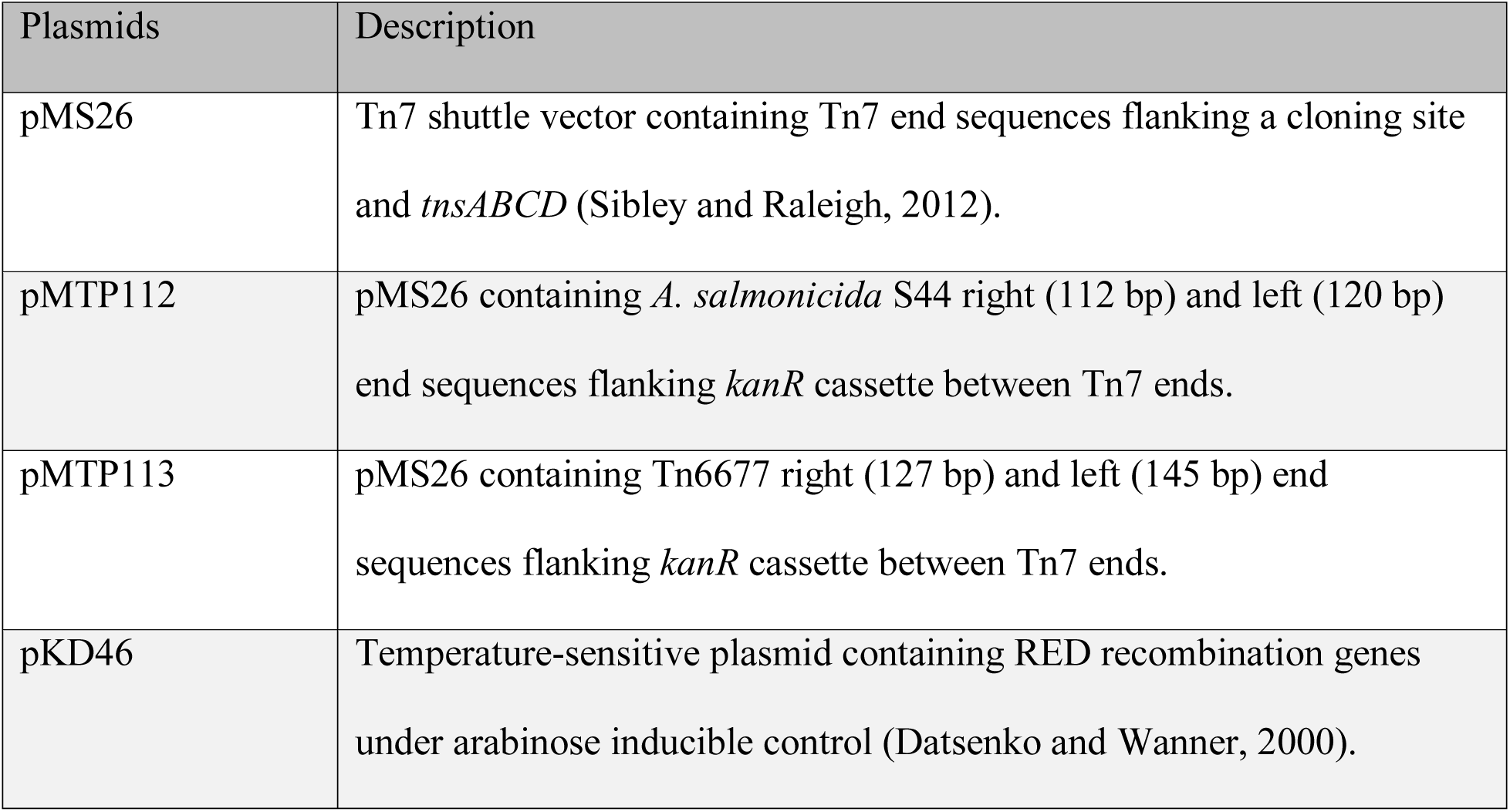
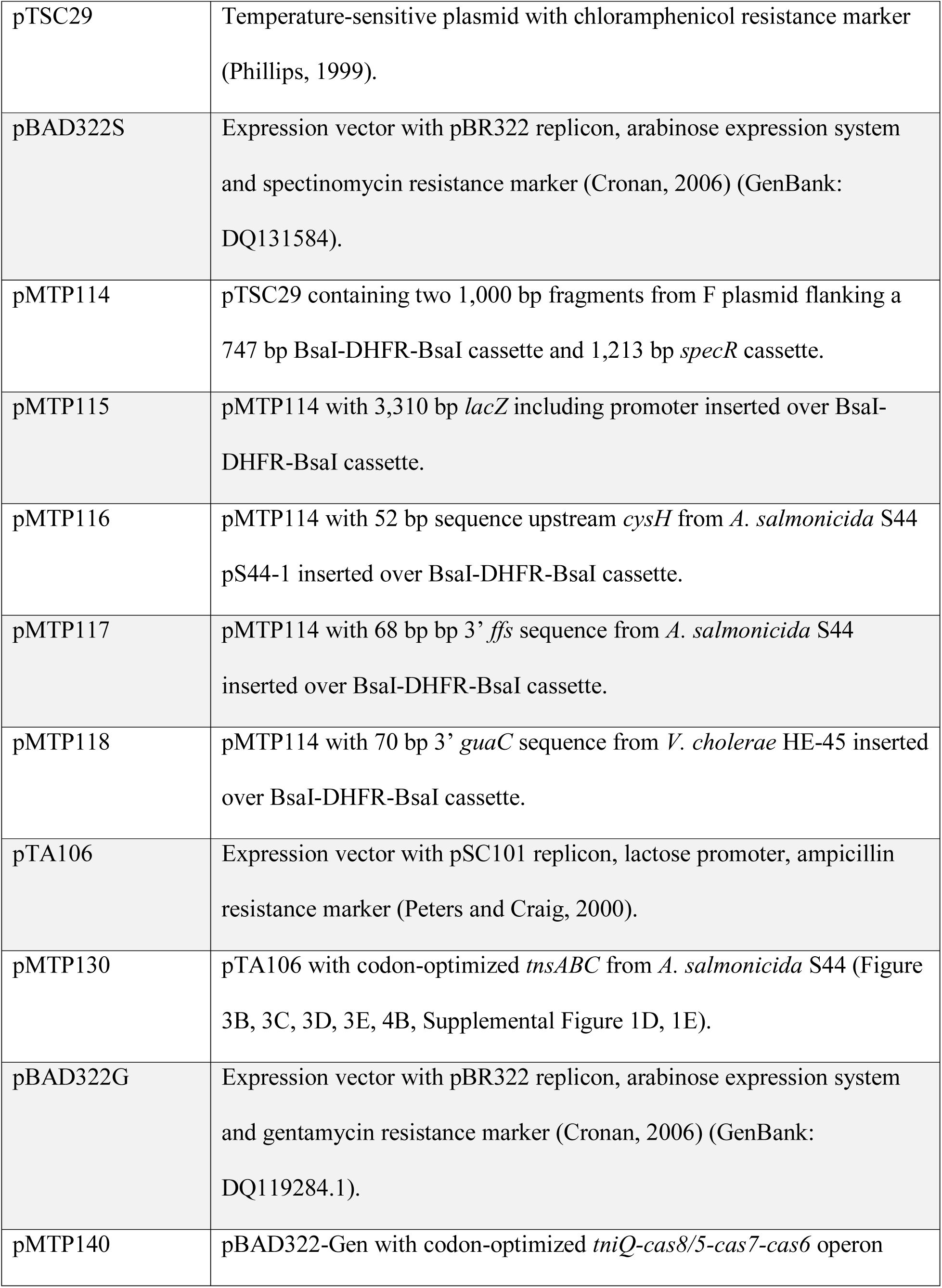
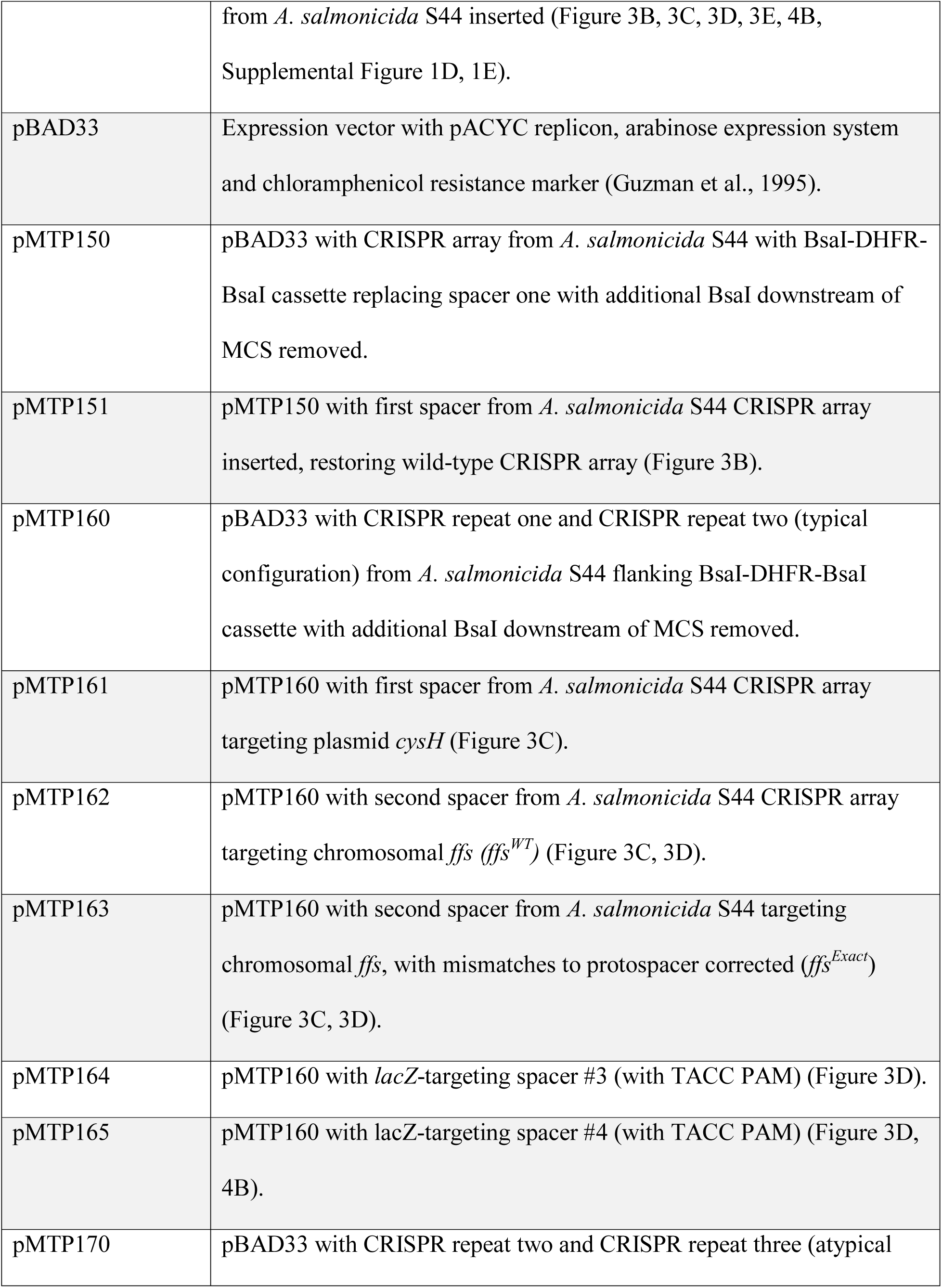
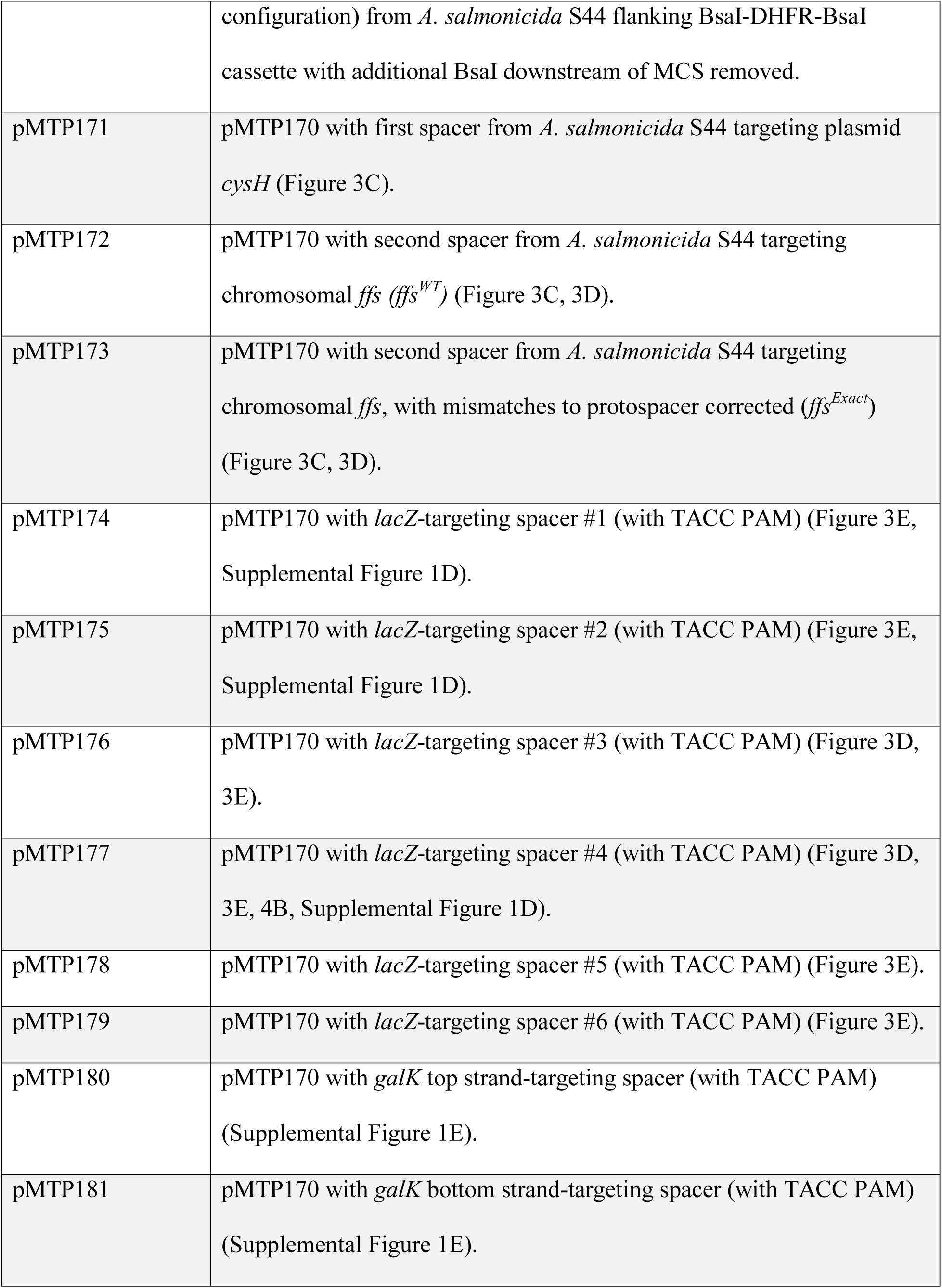
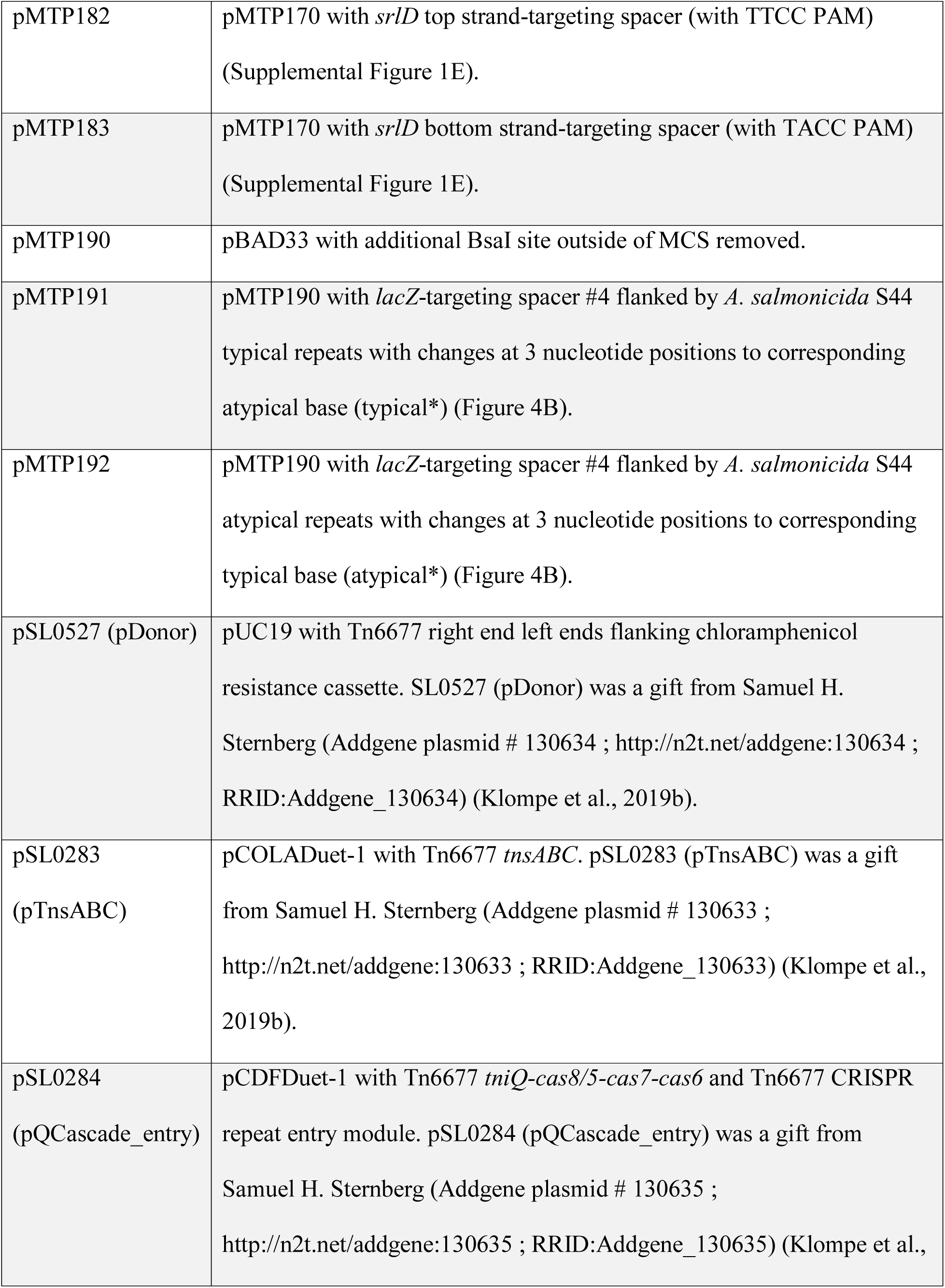
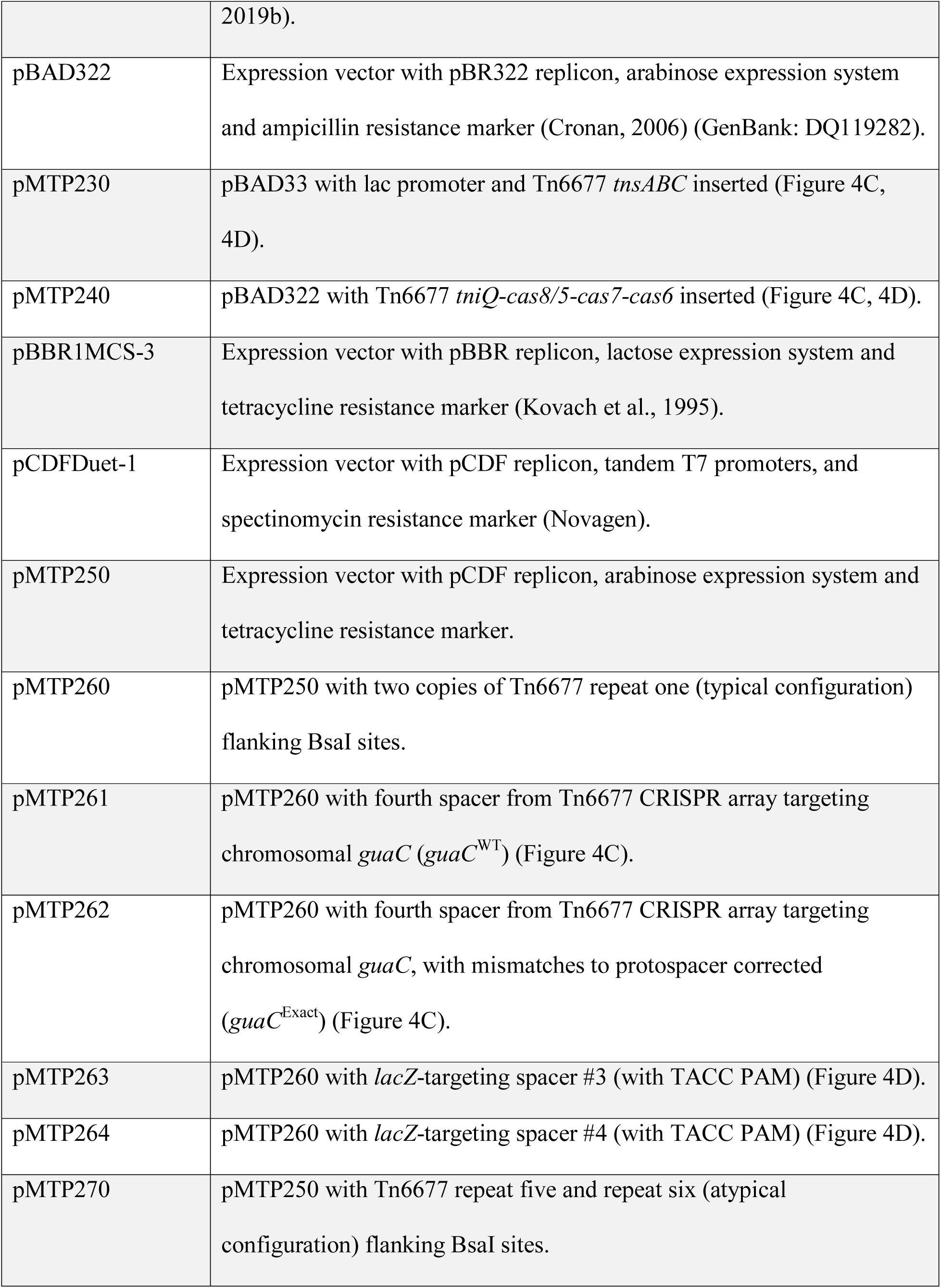
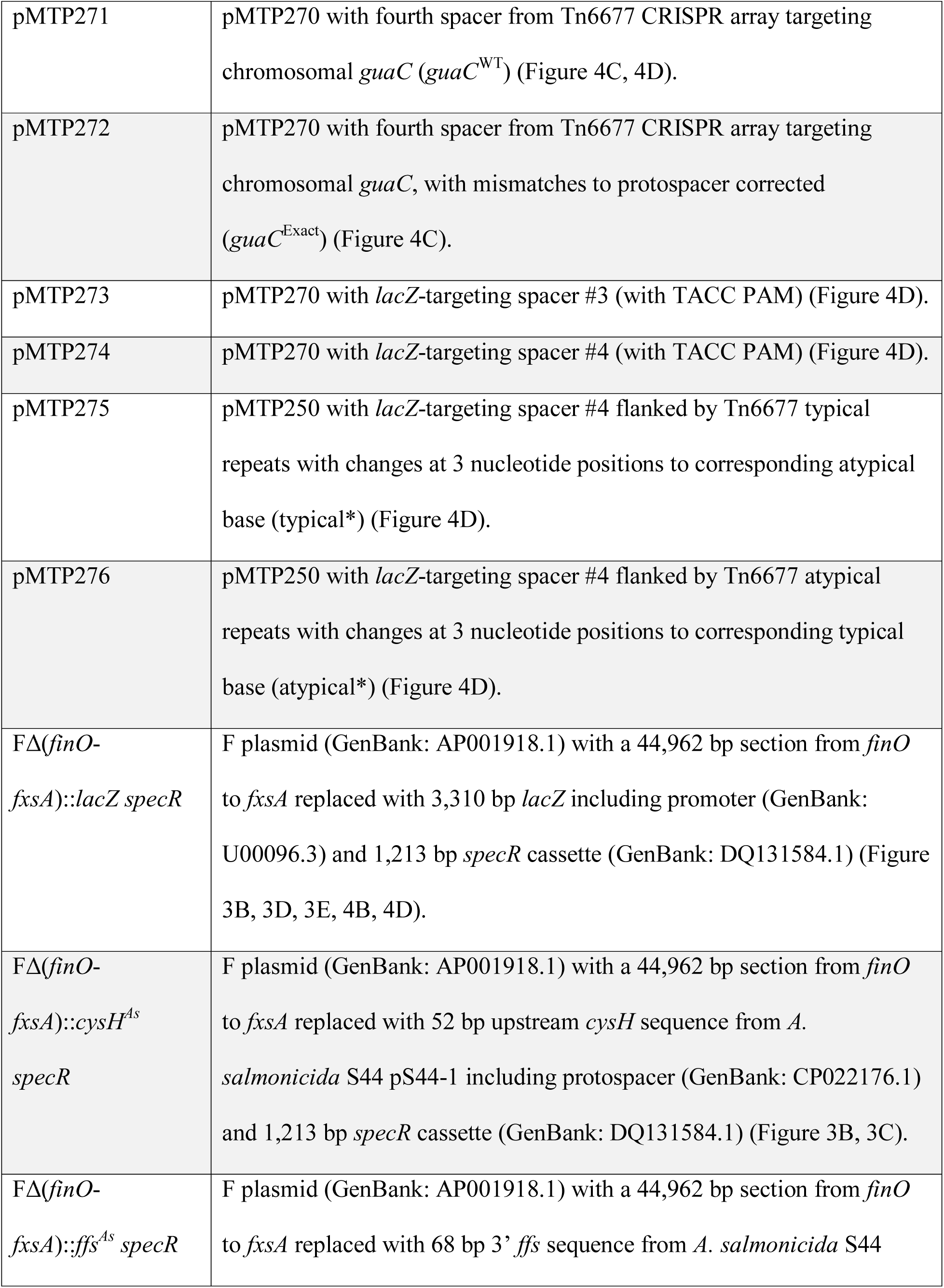
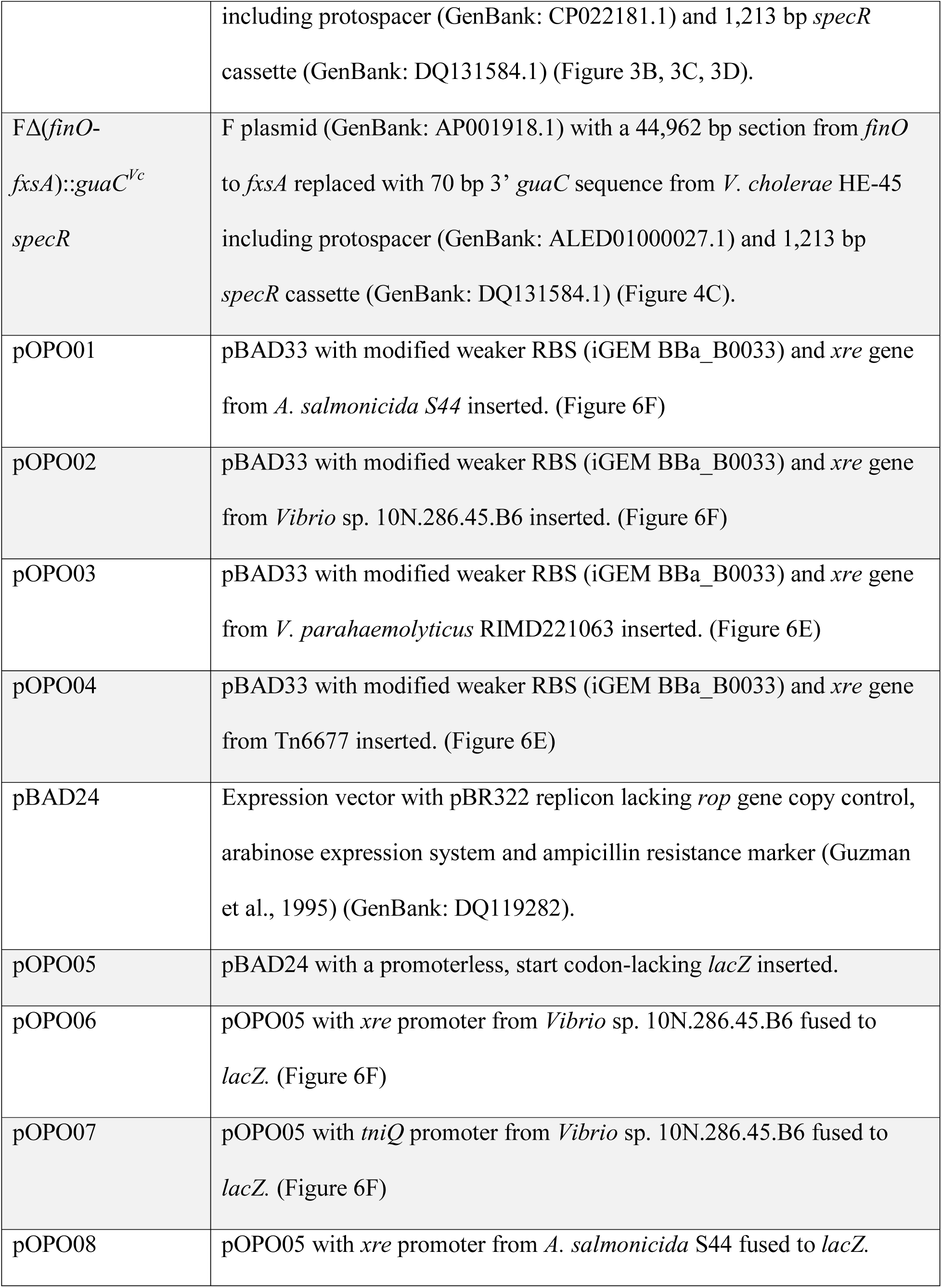
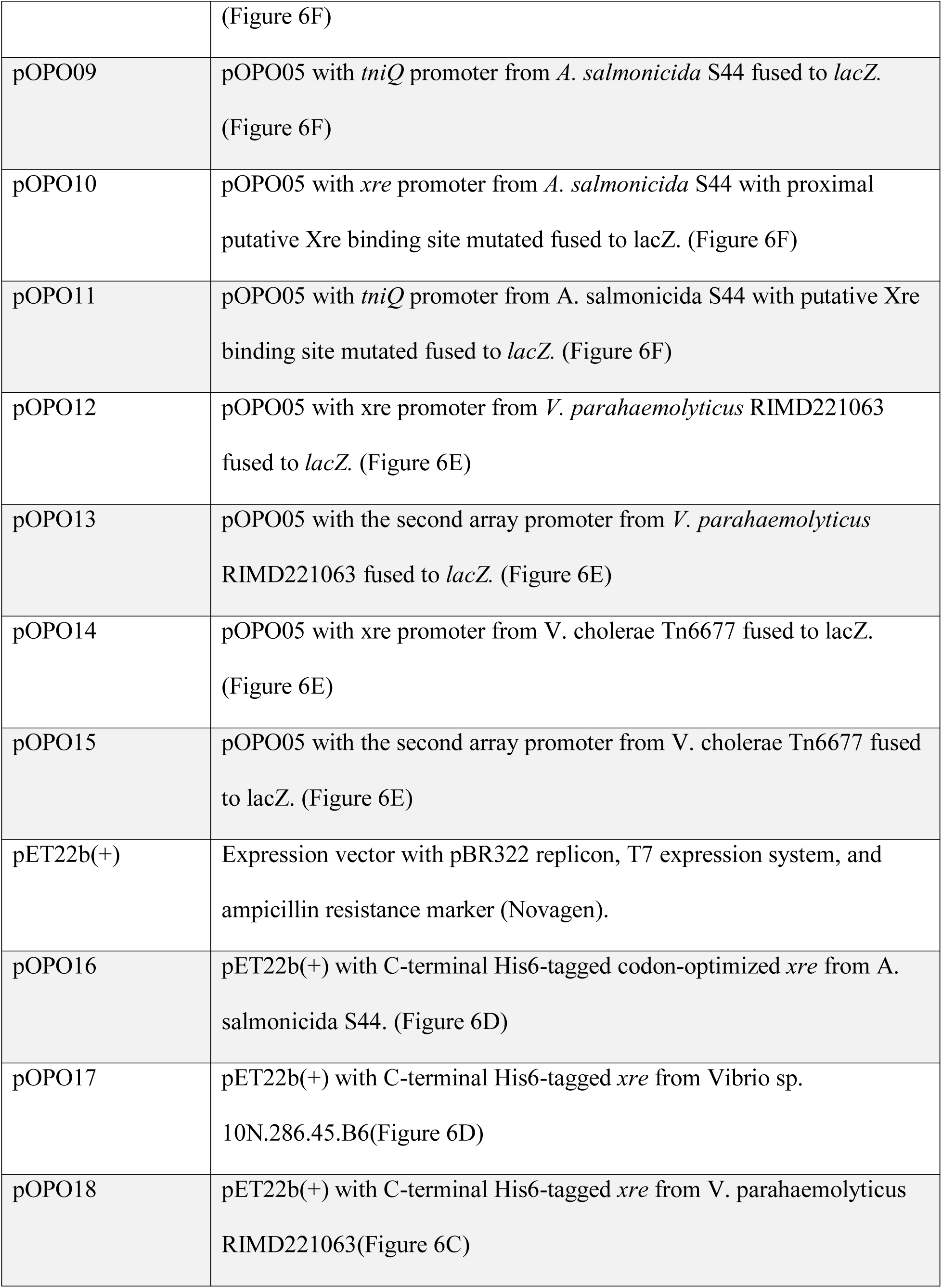
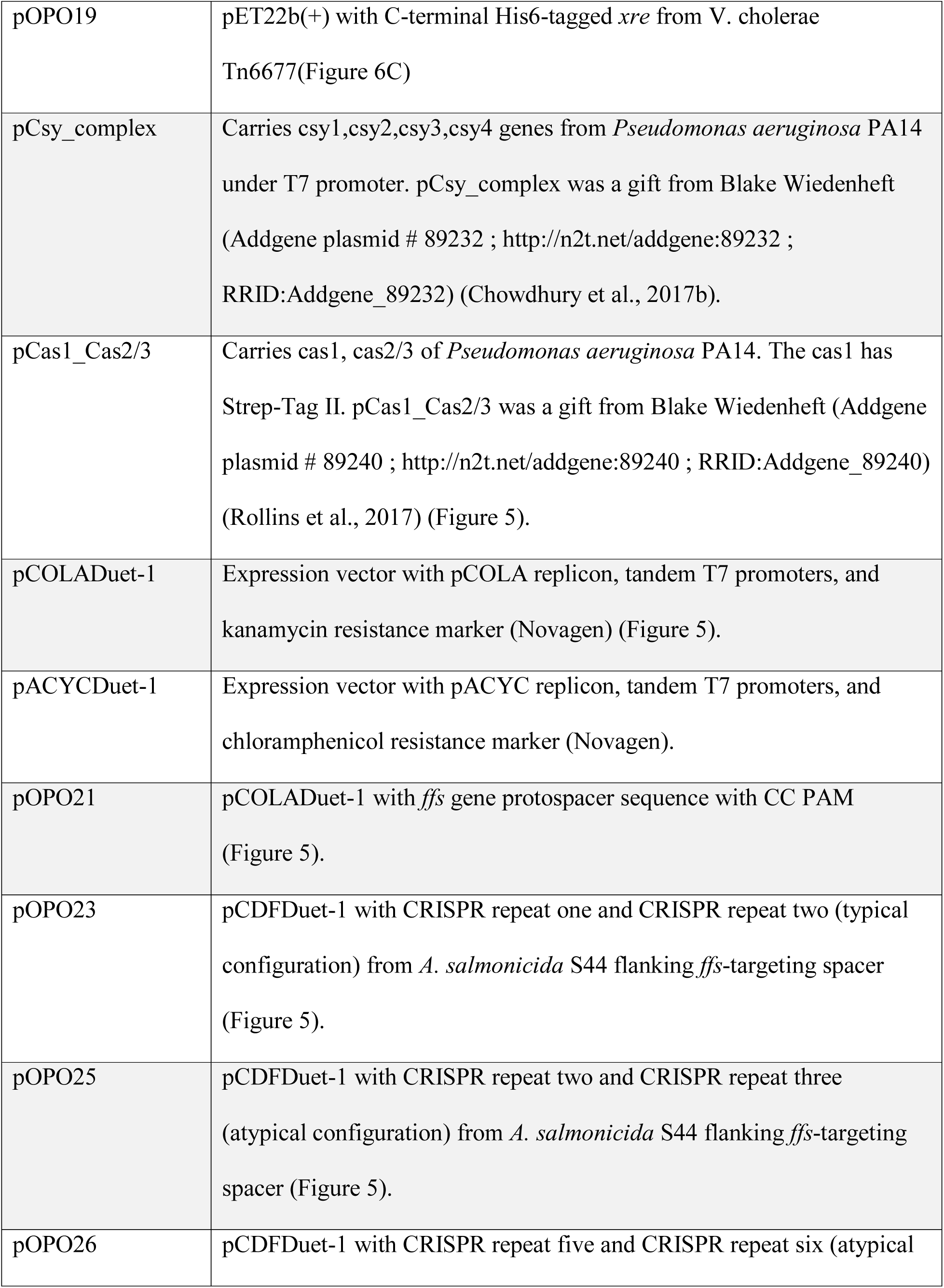
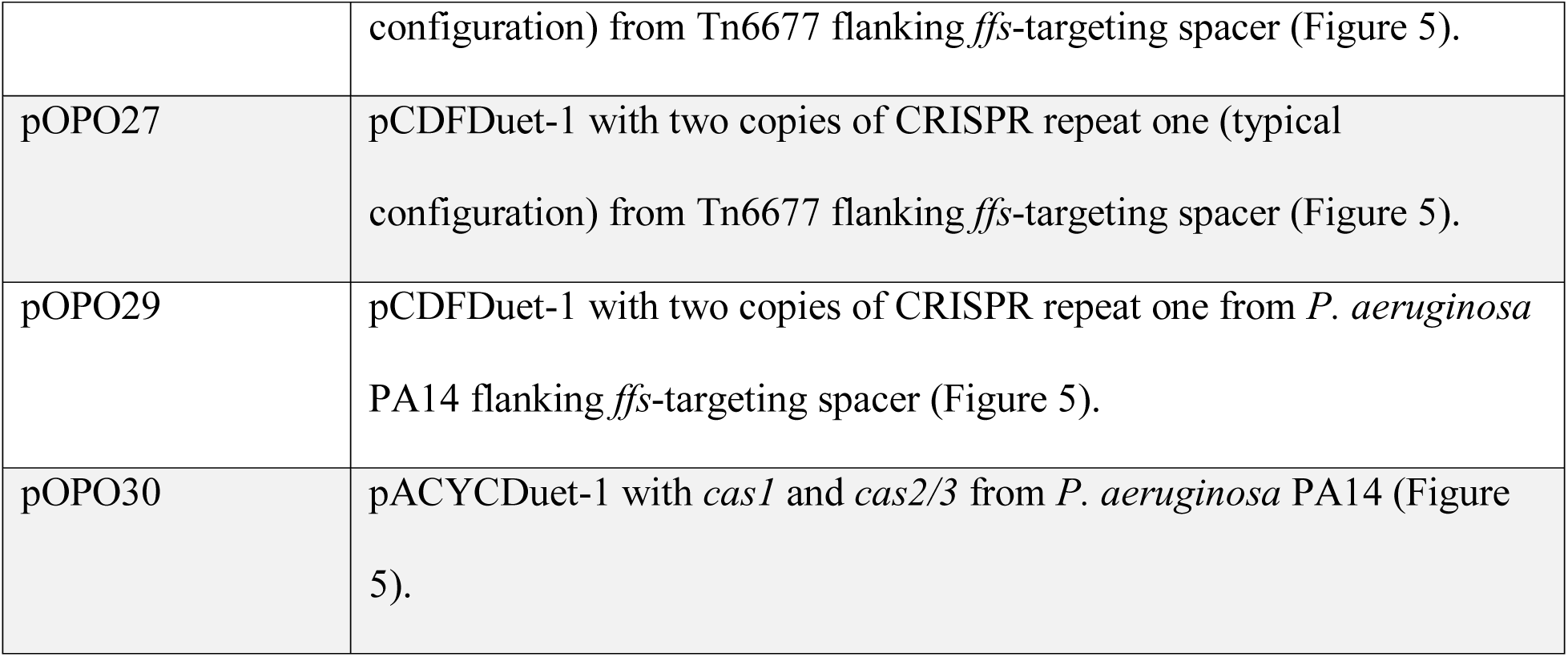

**Table 3.**
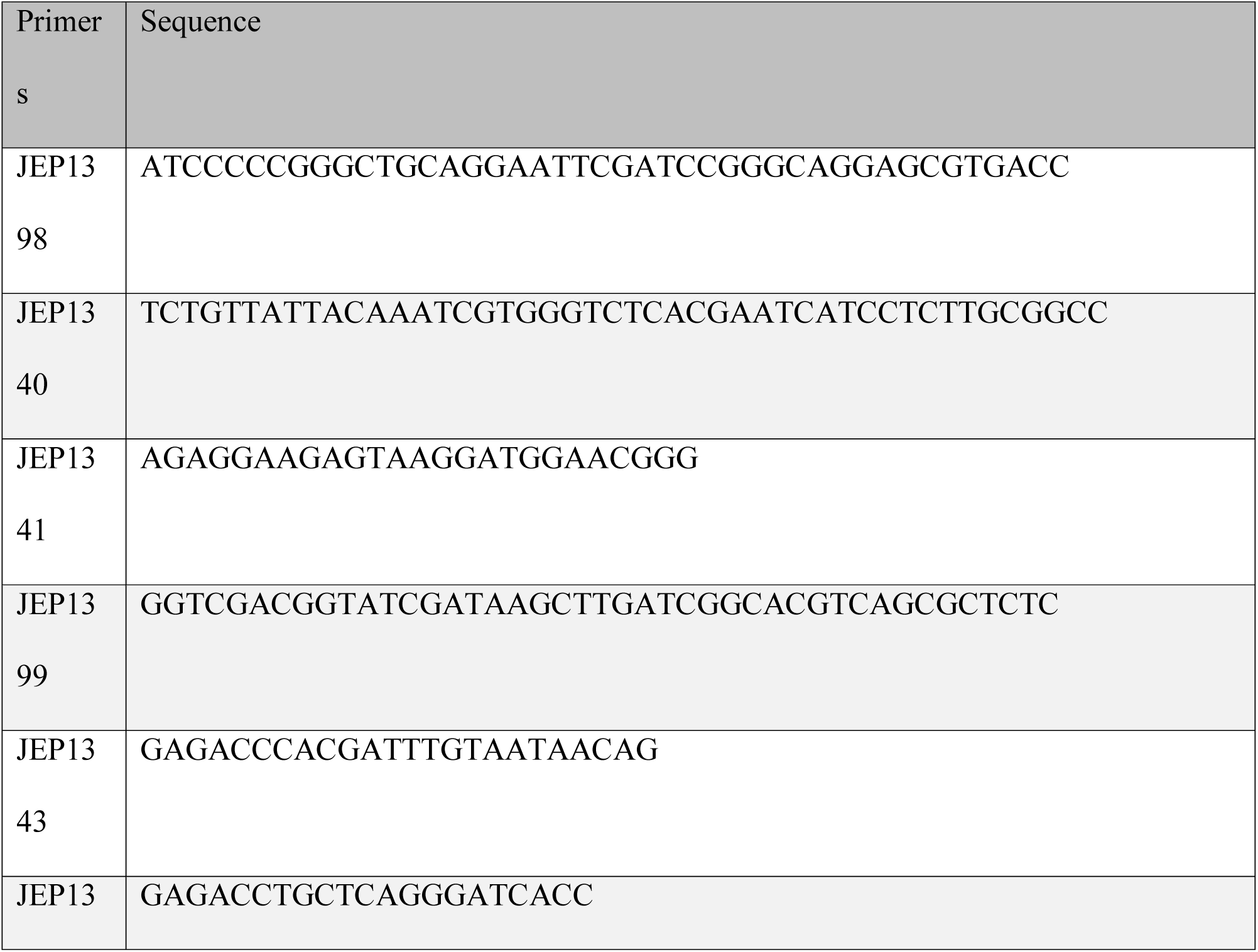
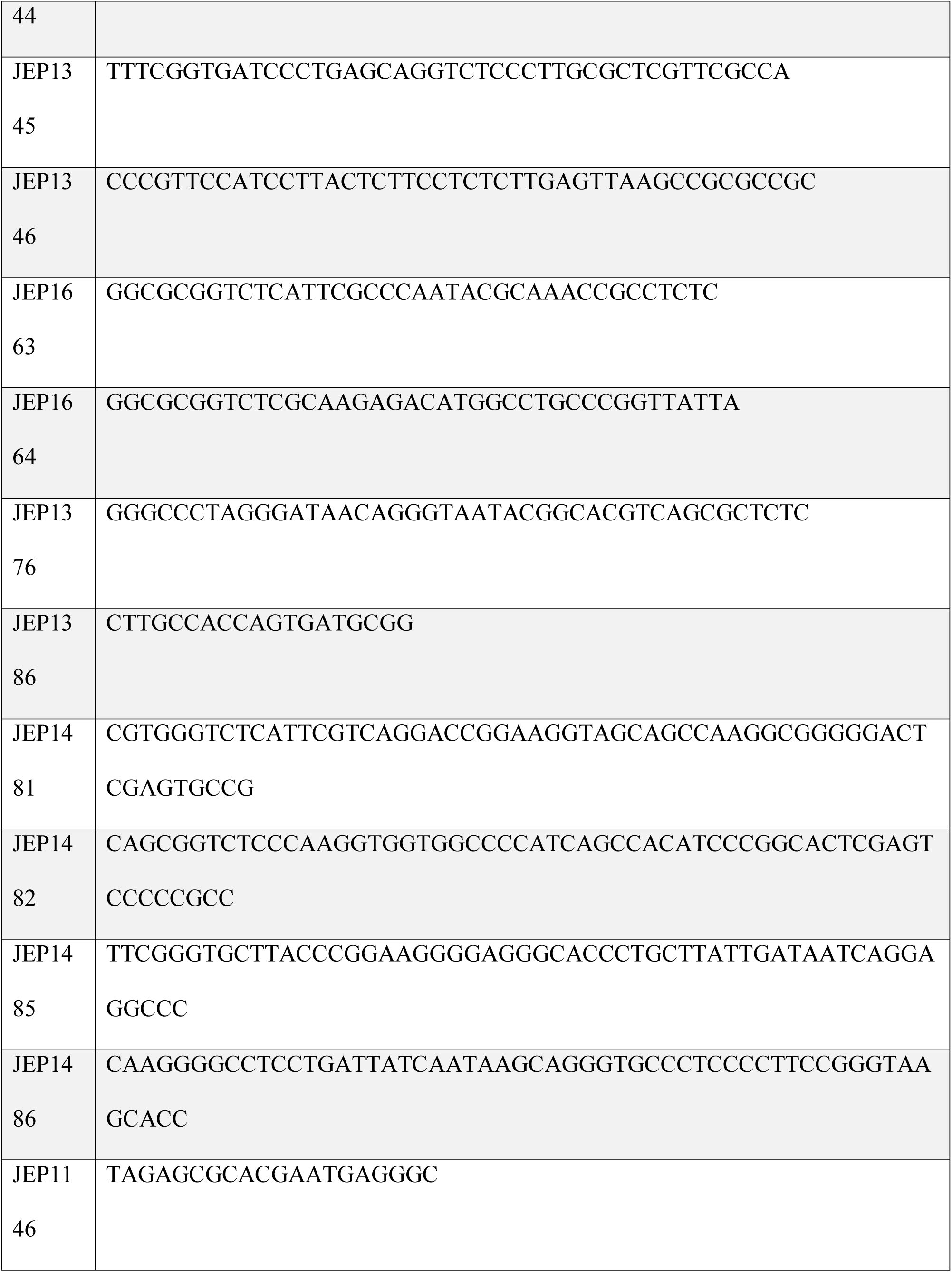
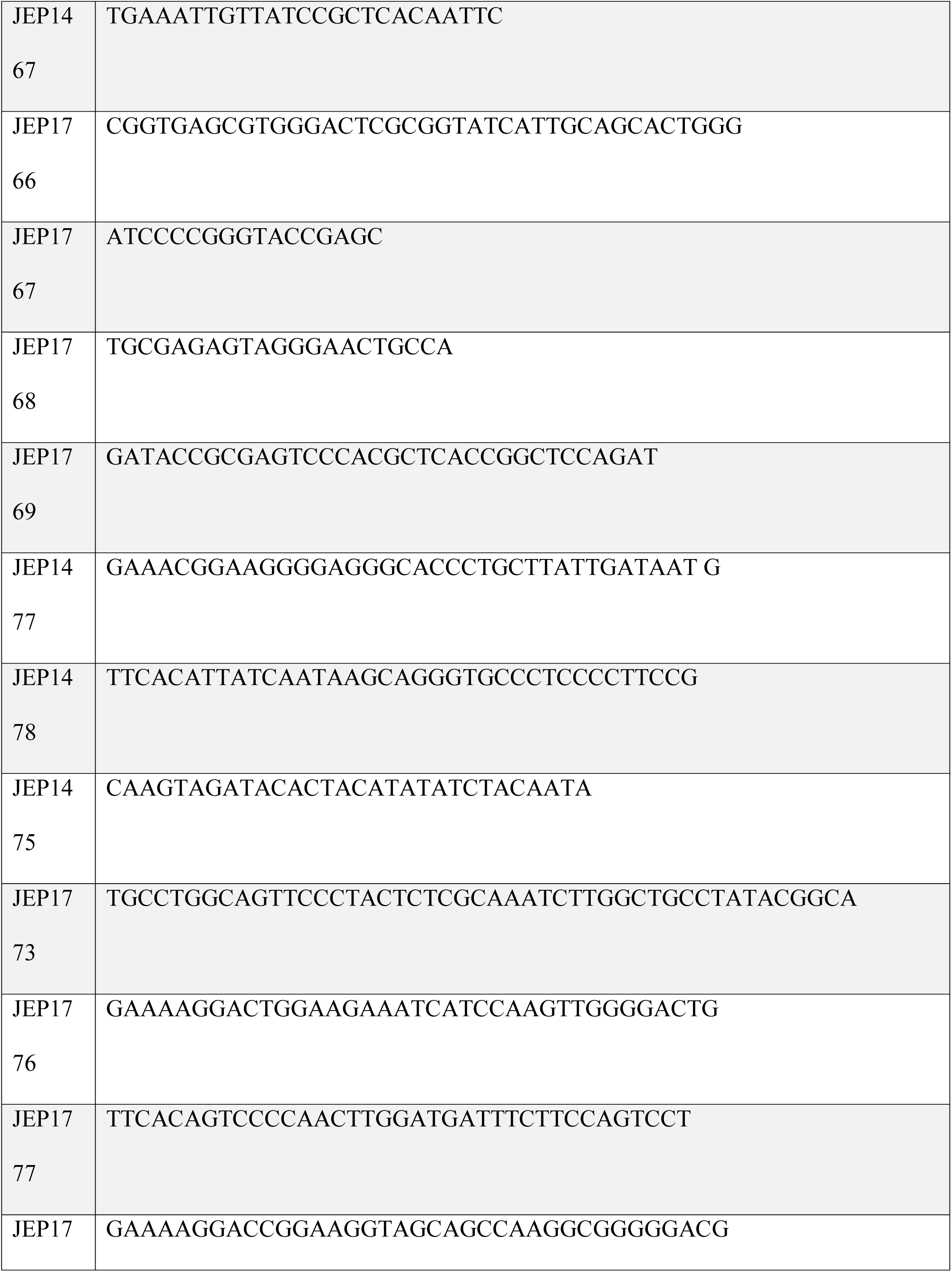
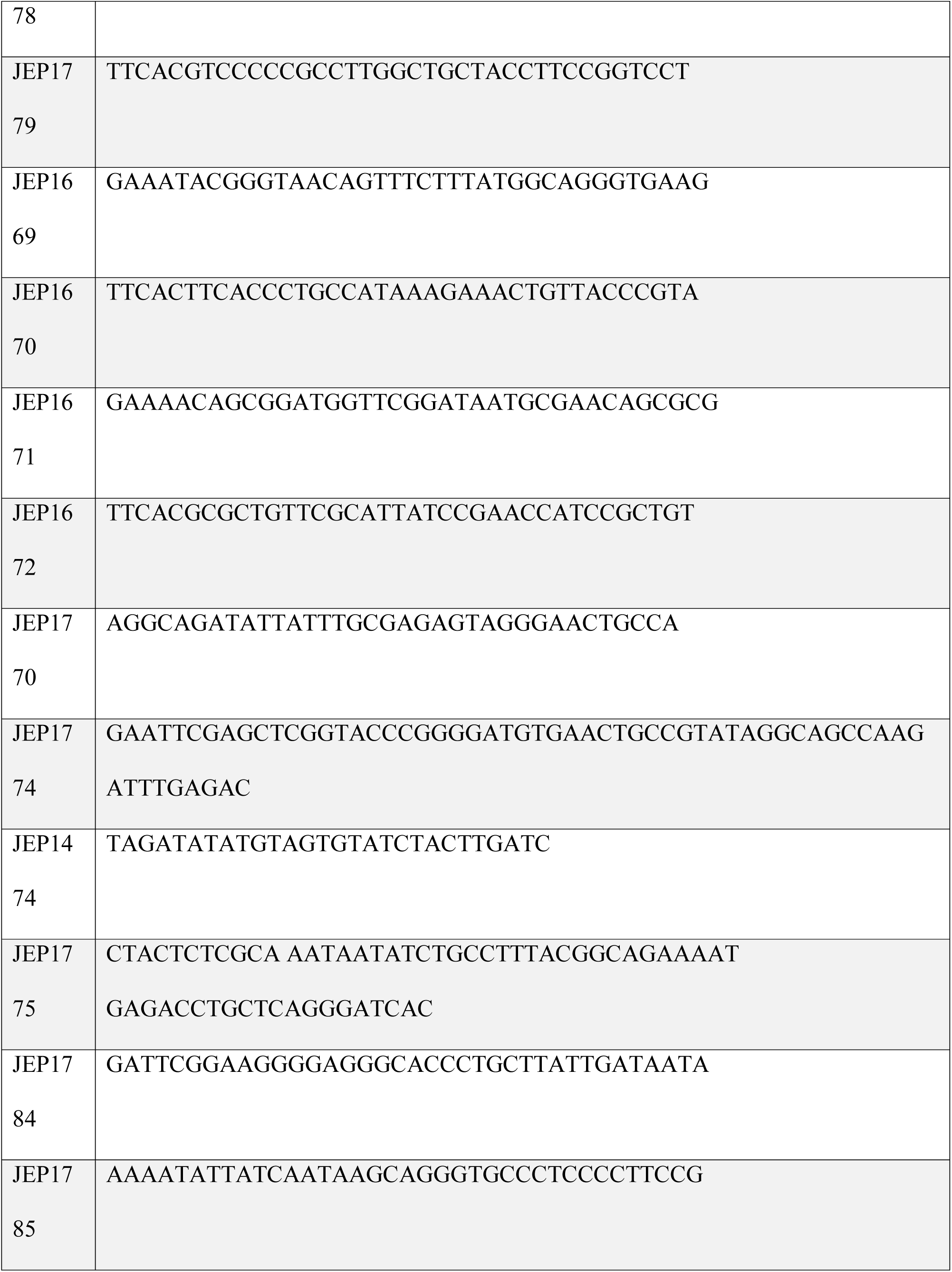
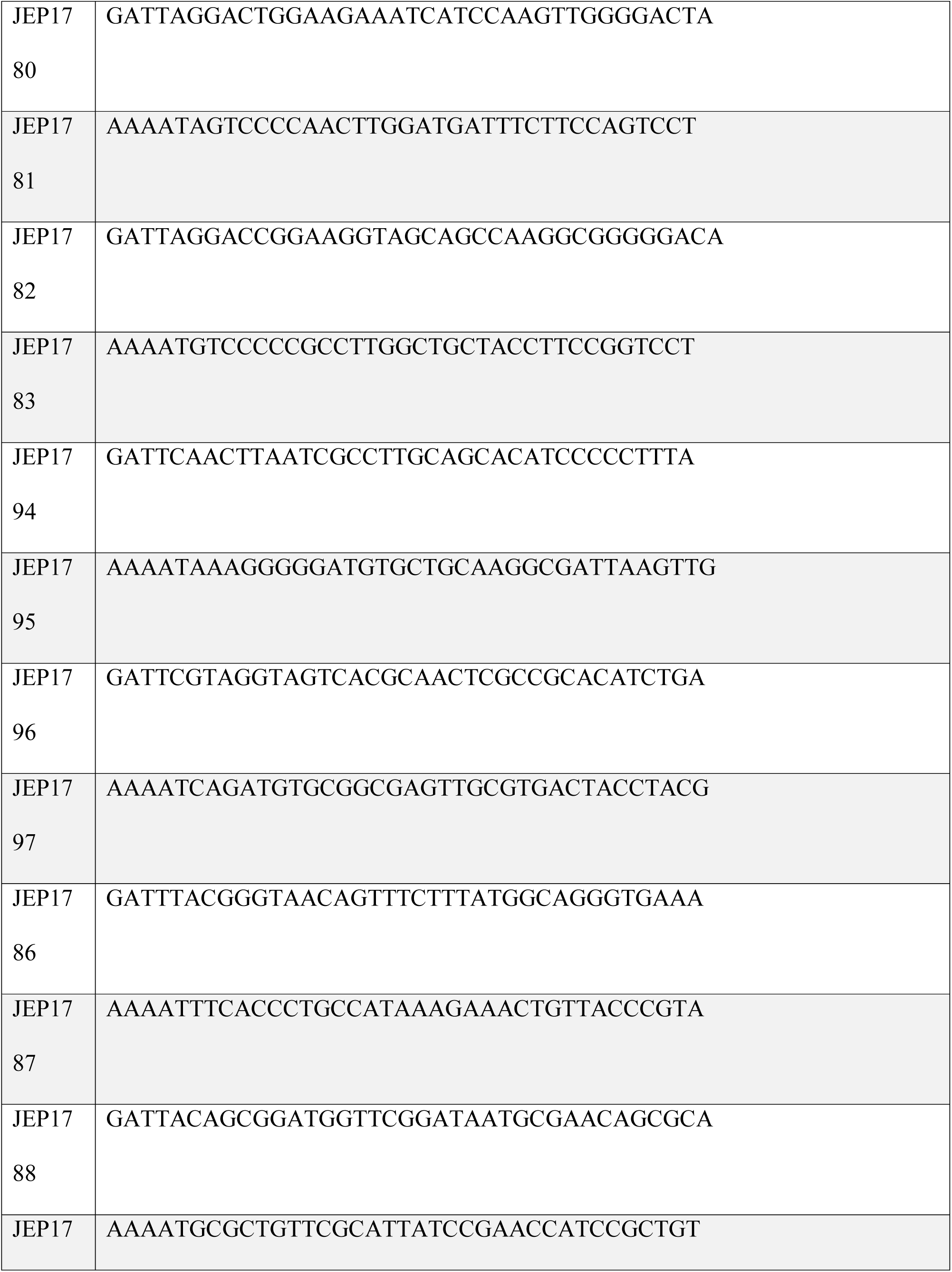
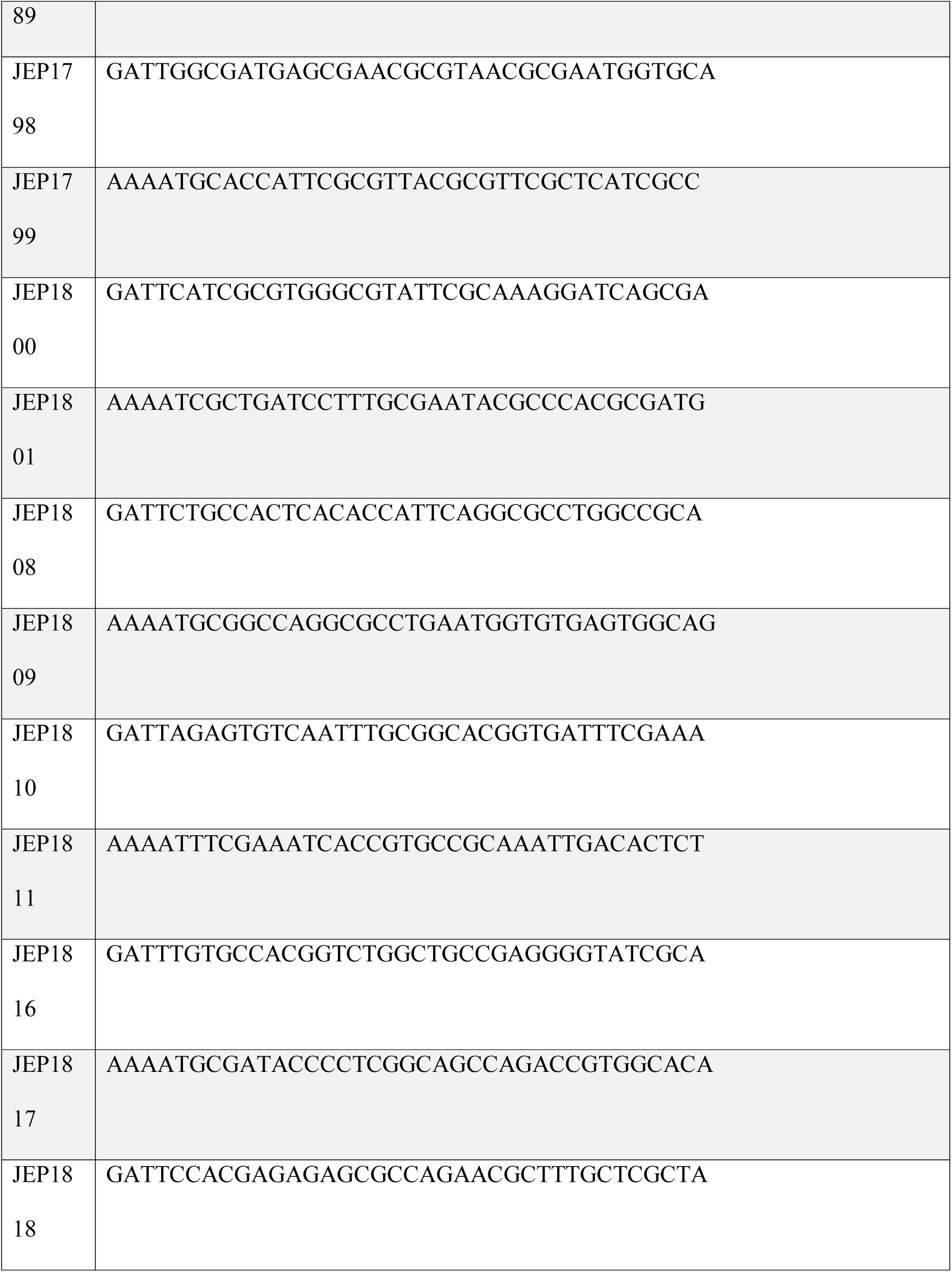
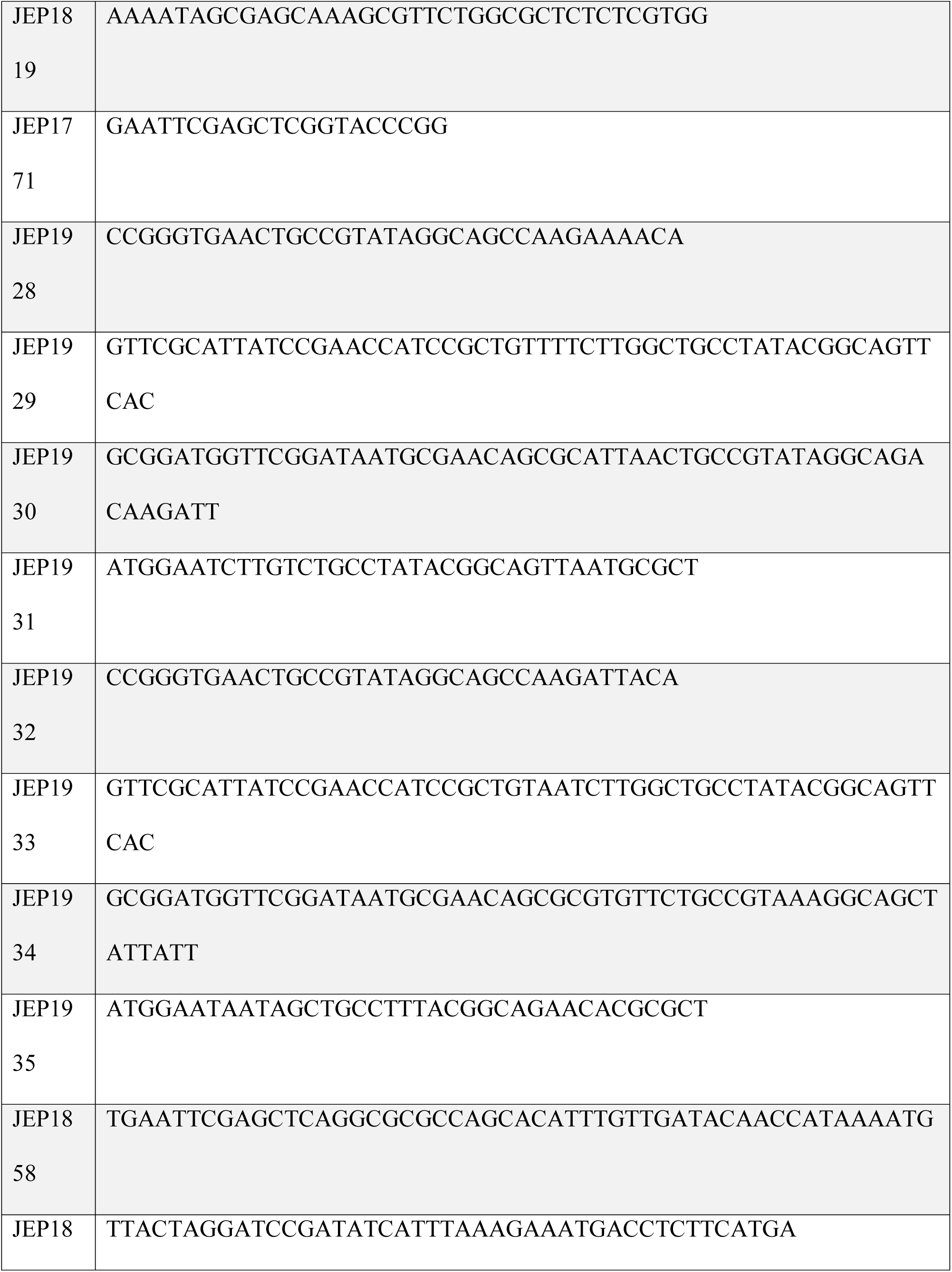
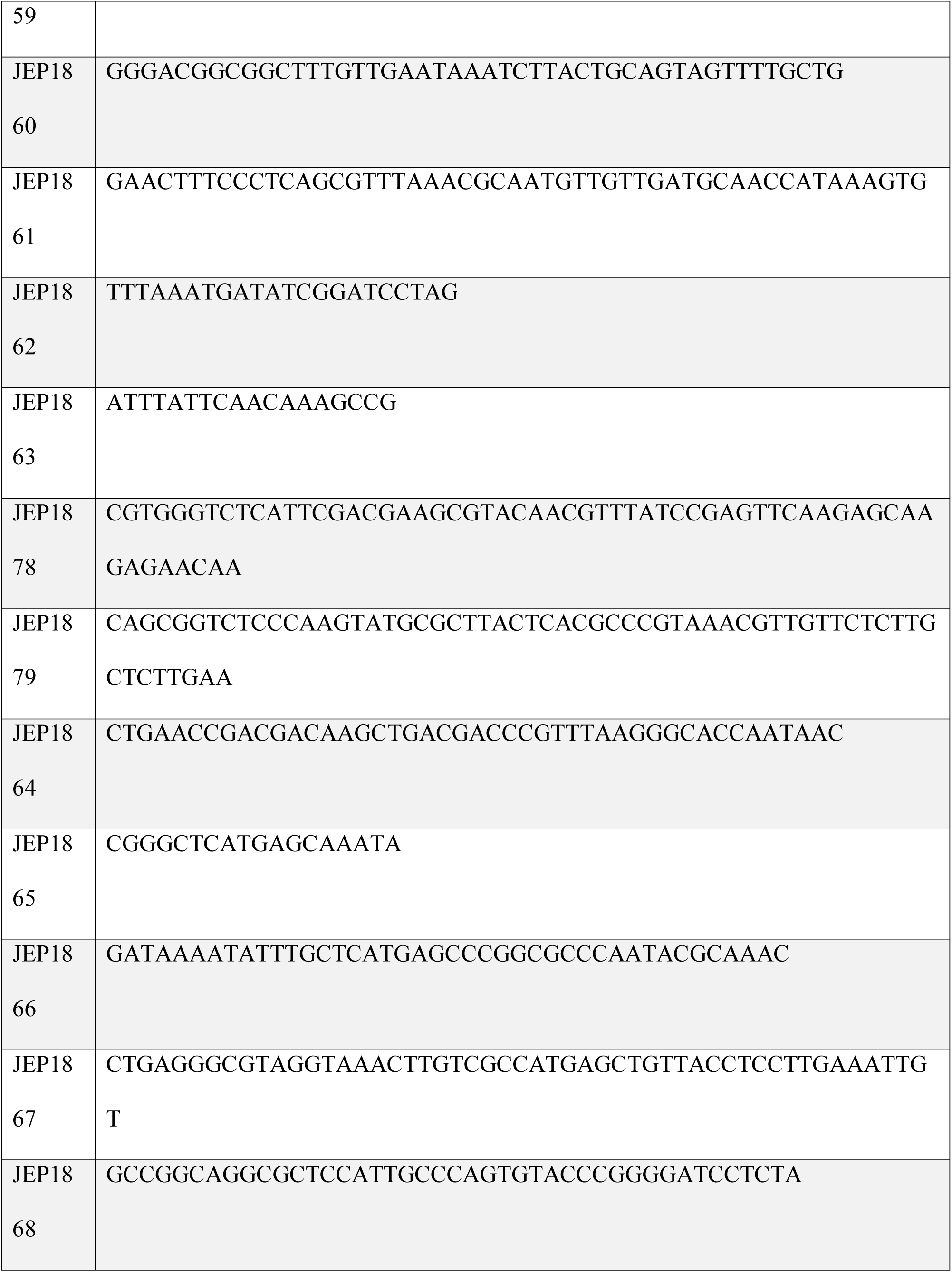
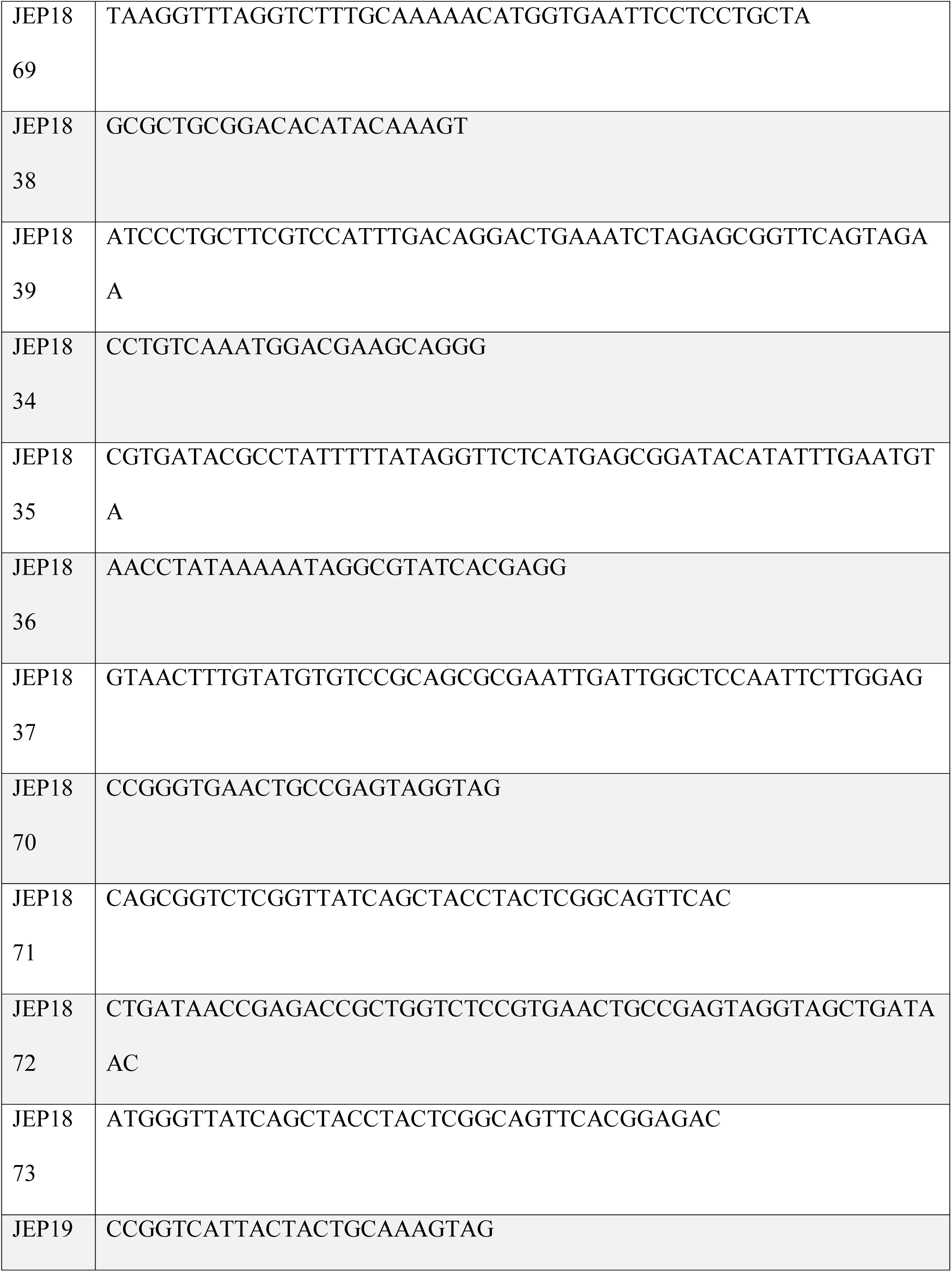
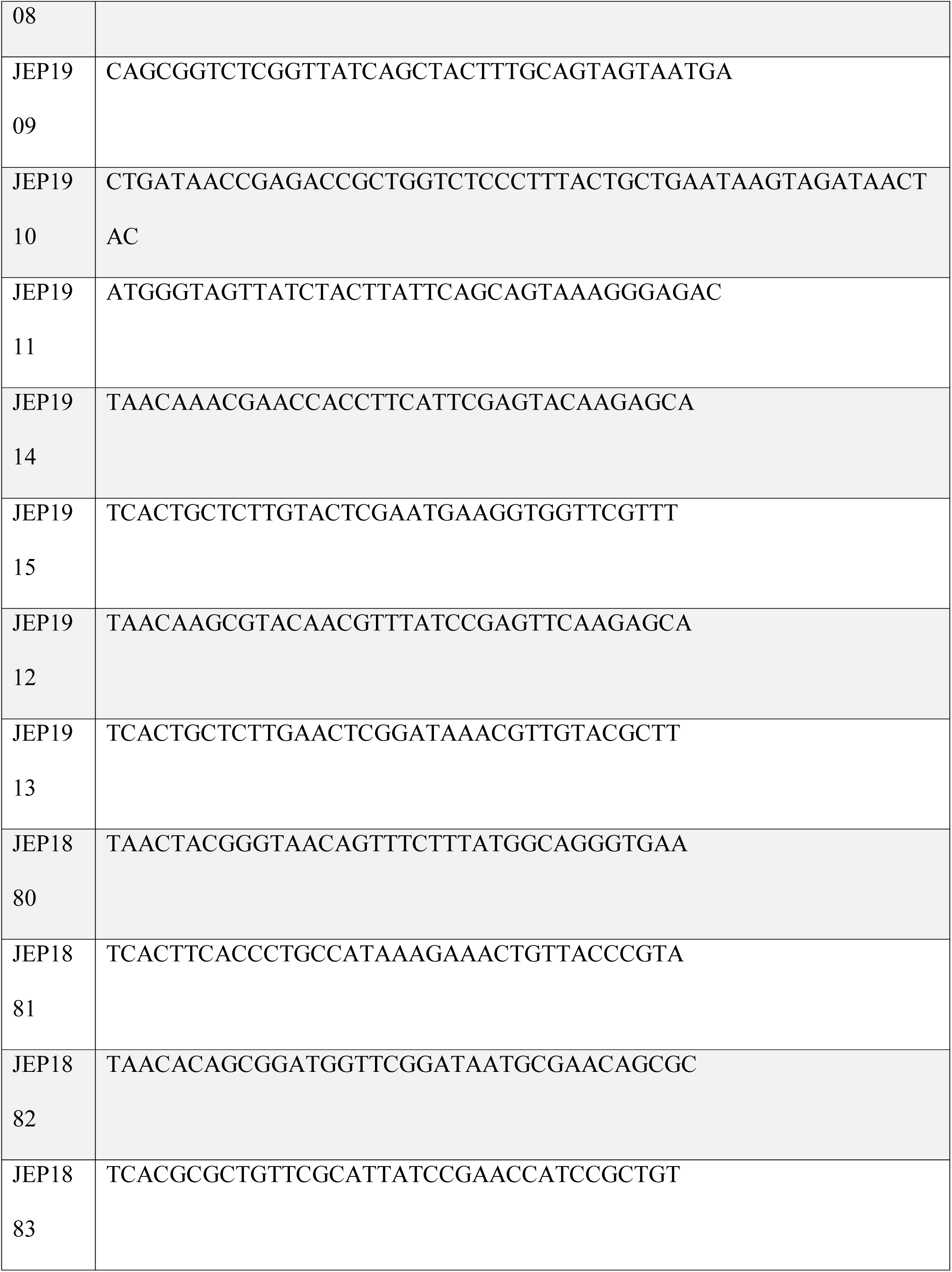
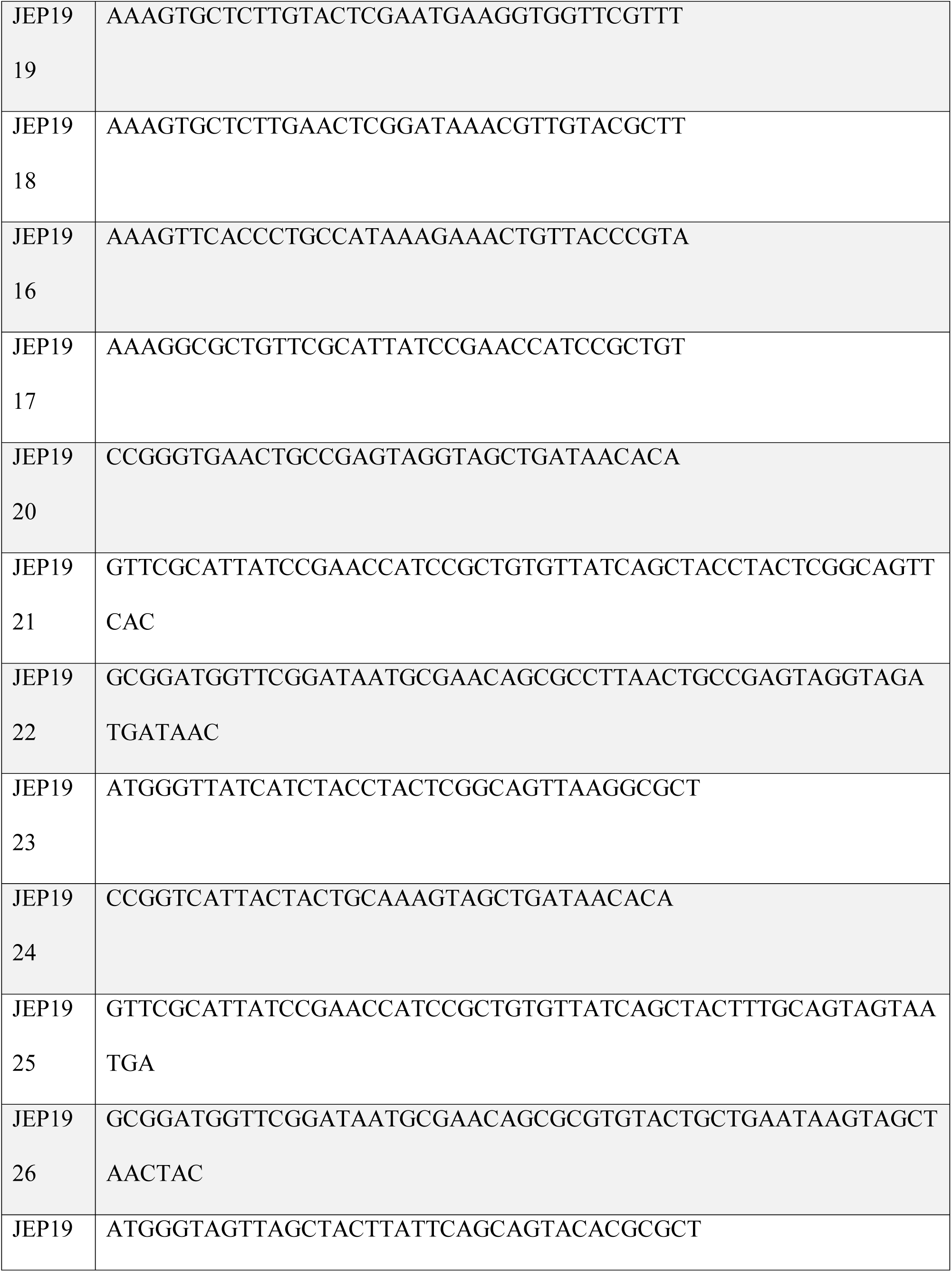
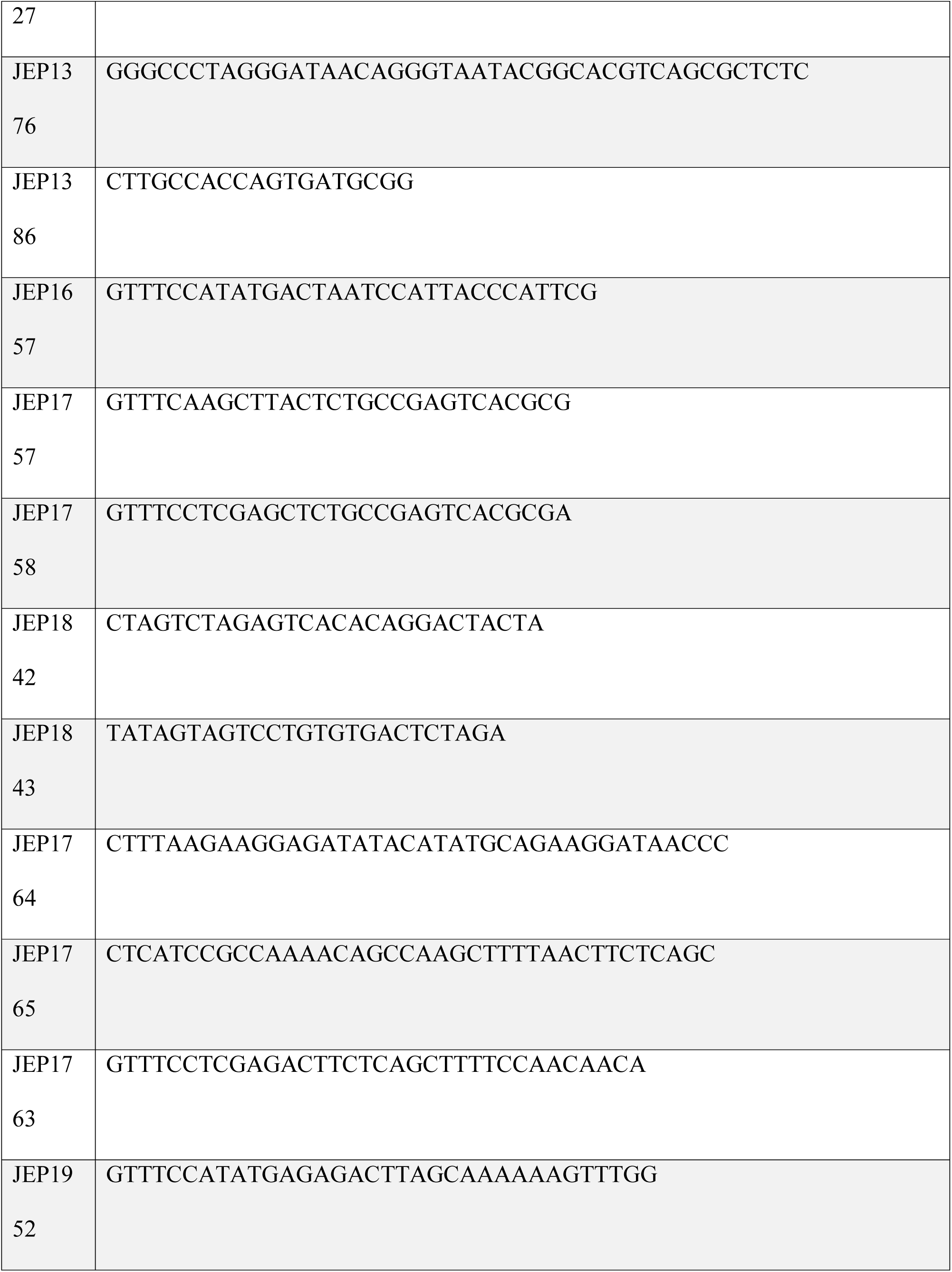
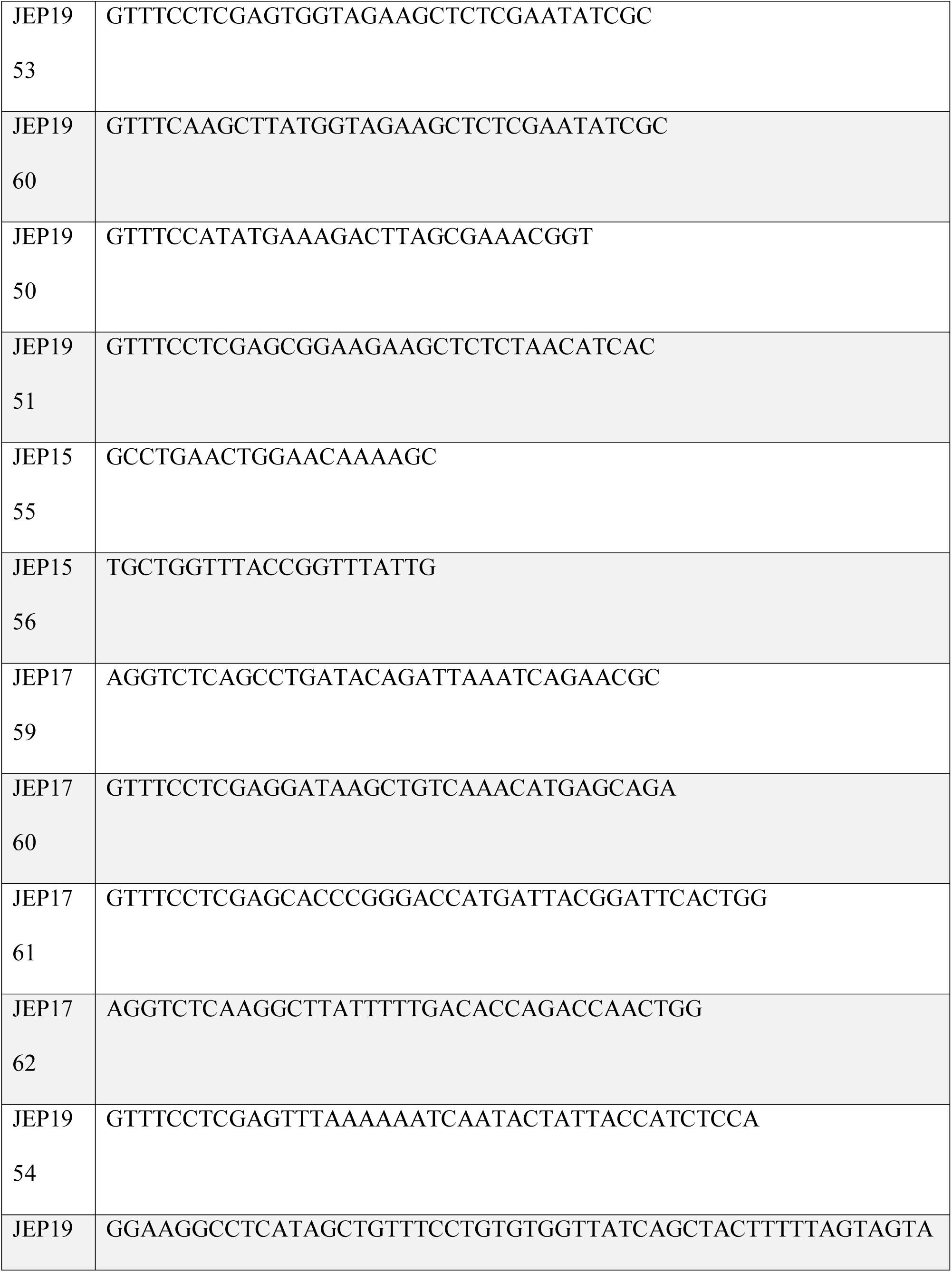
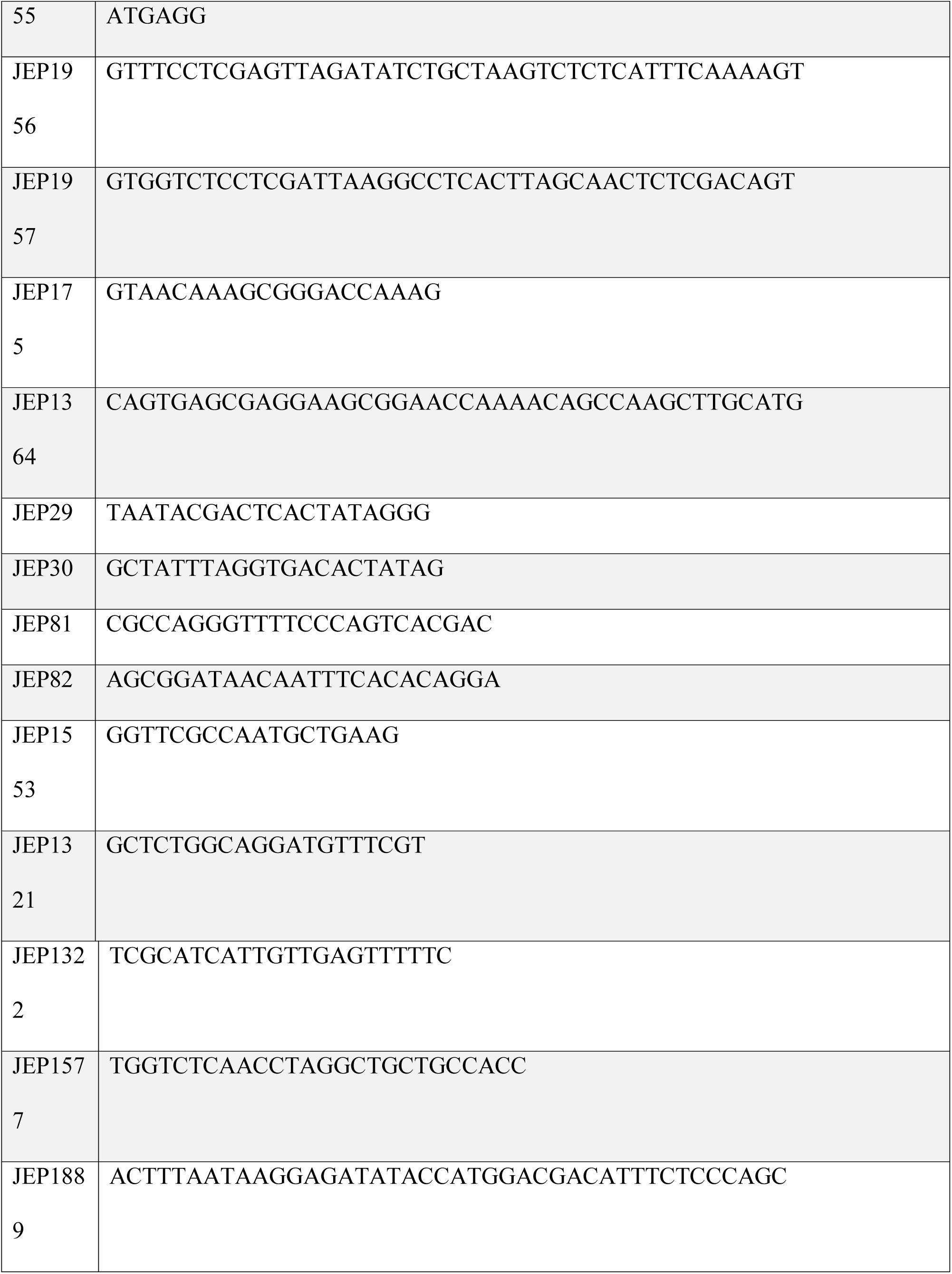
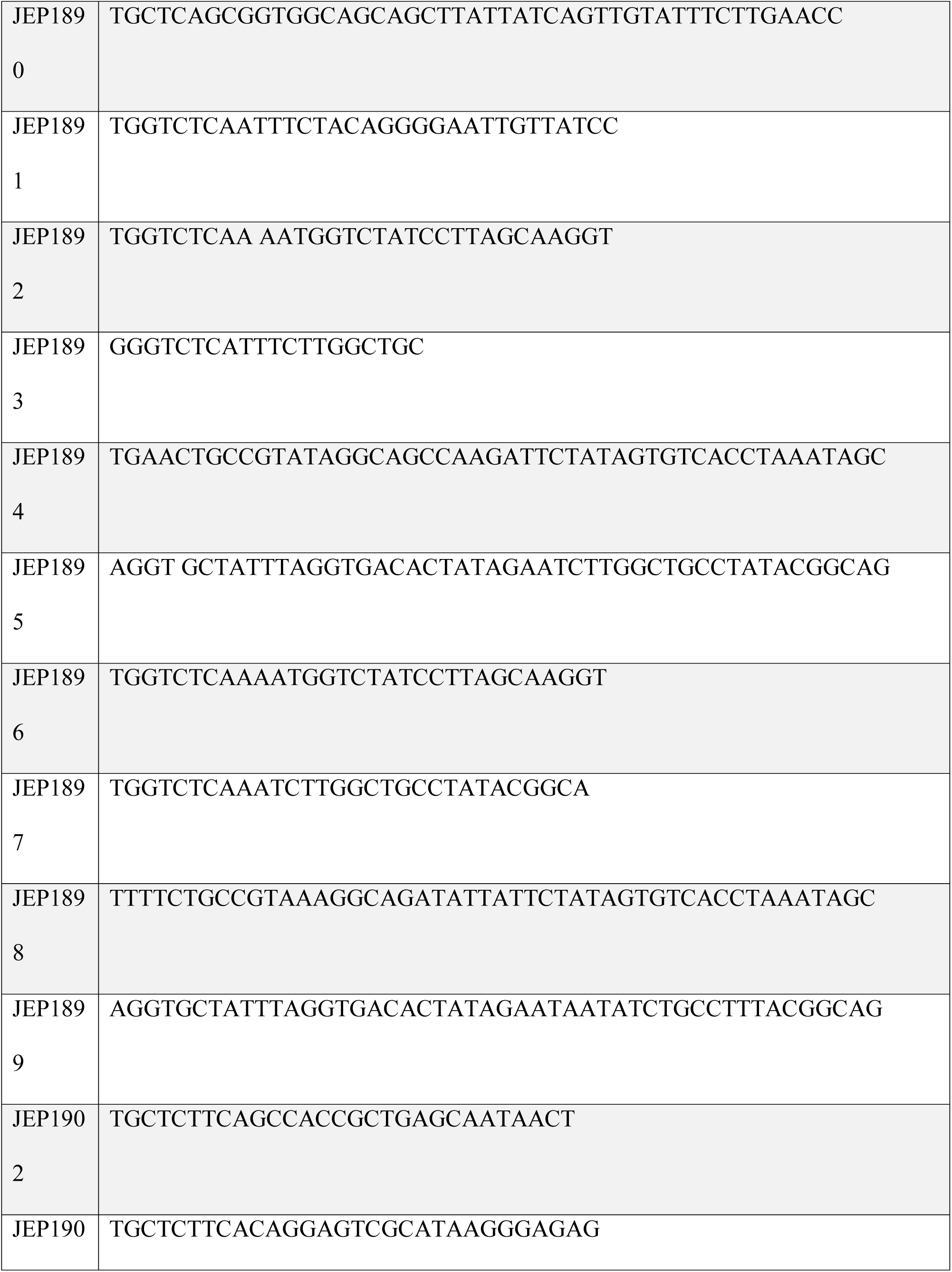
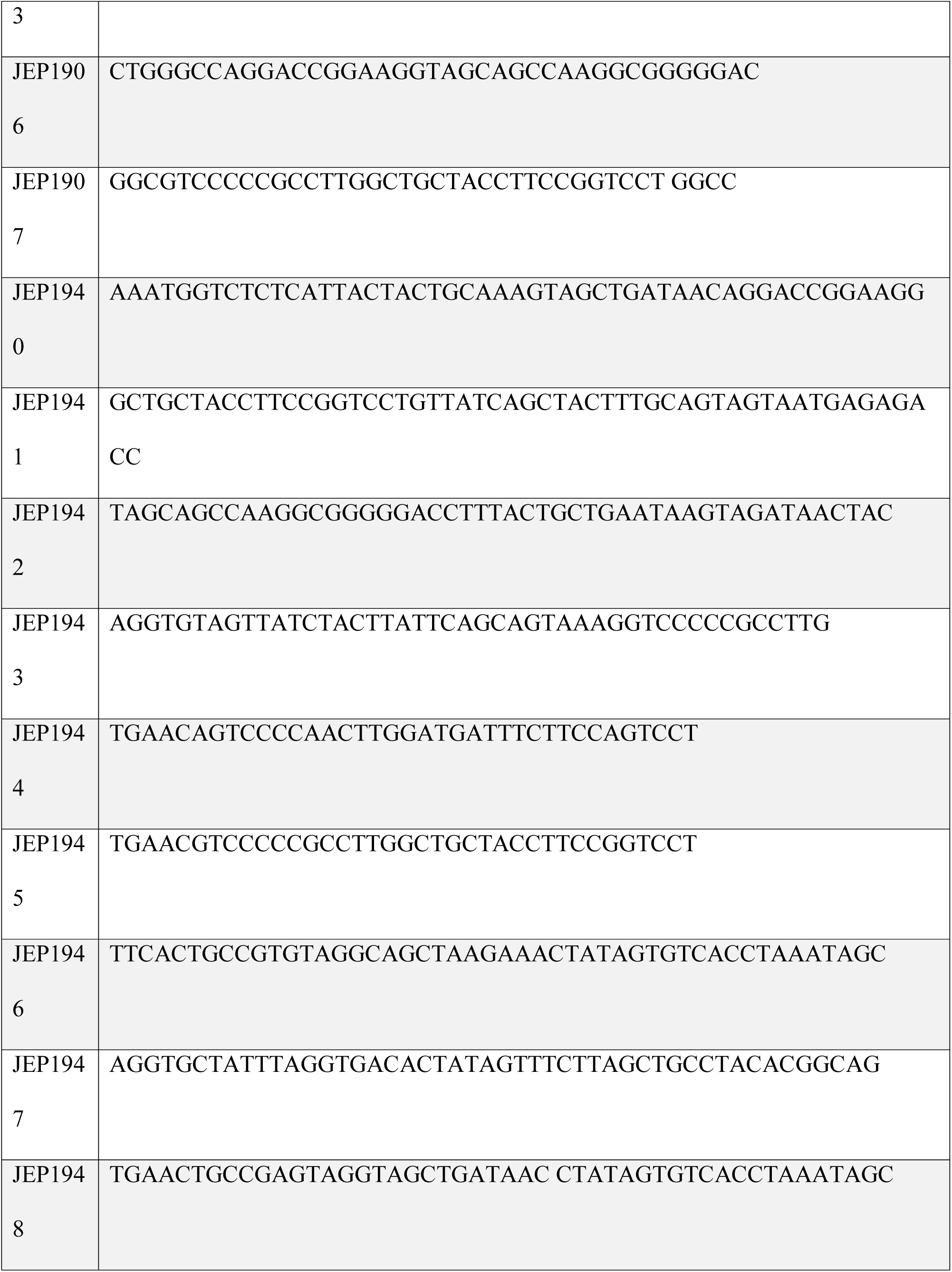
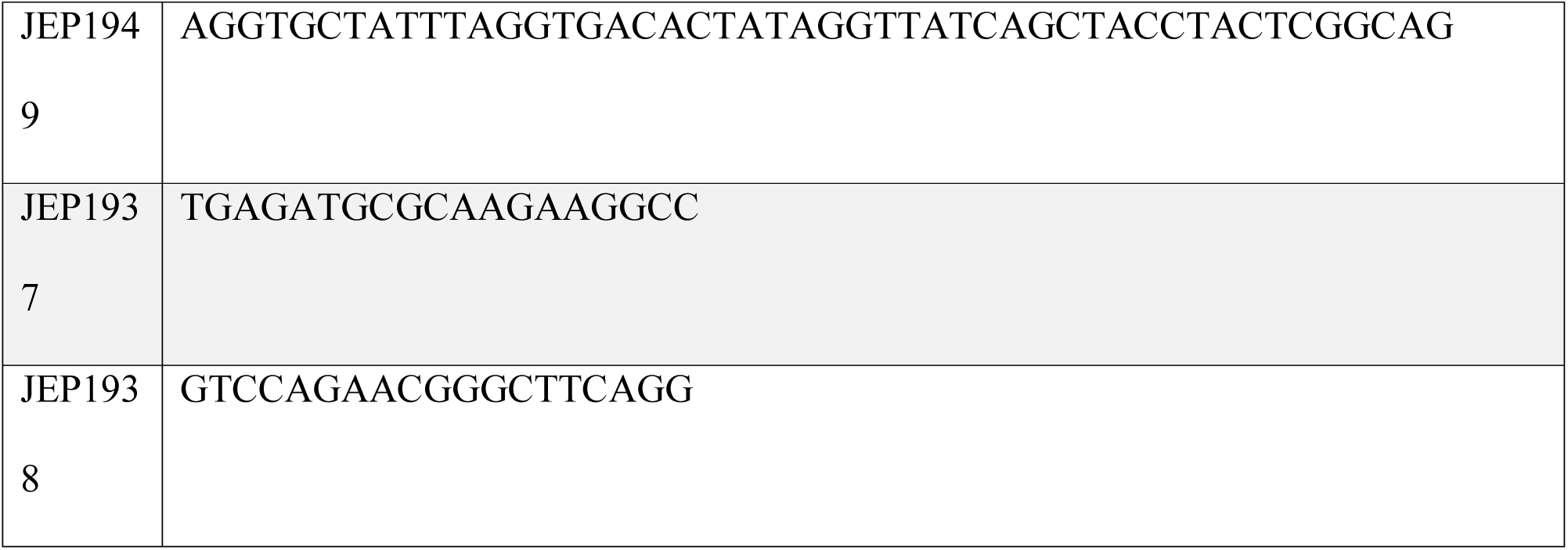

**Table 4.**
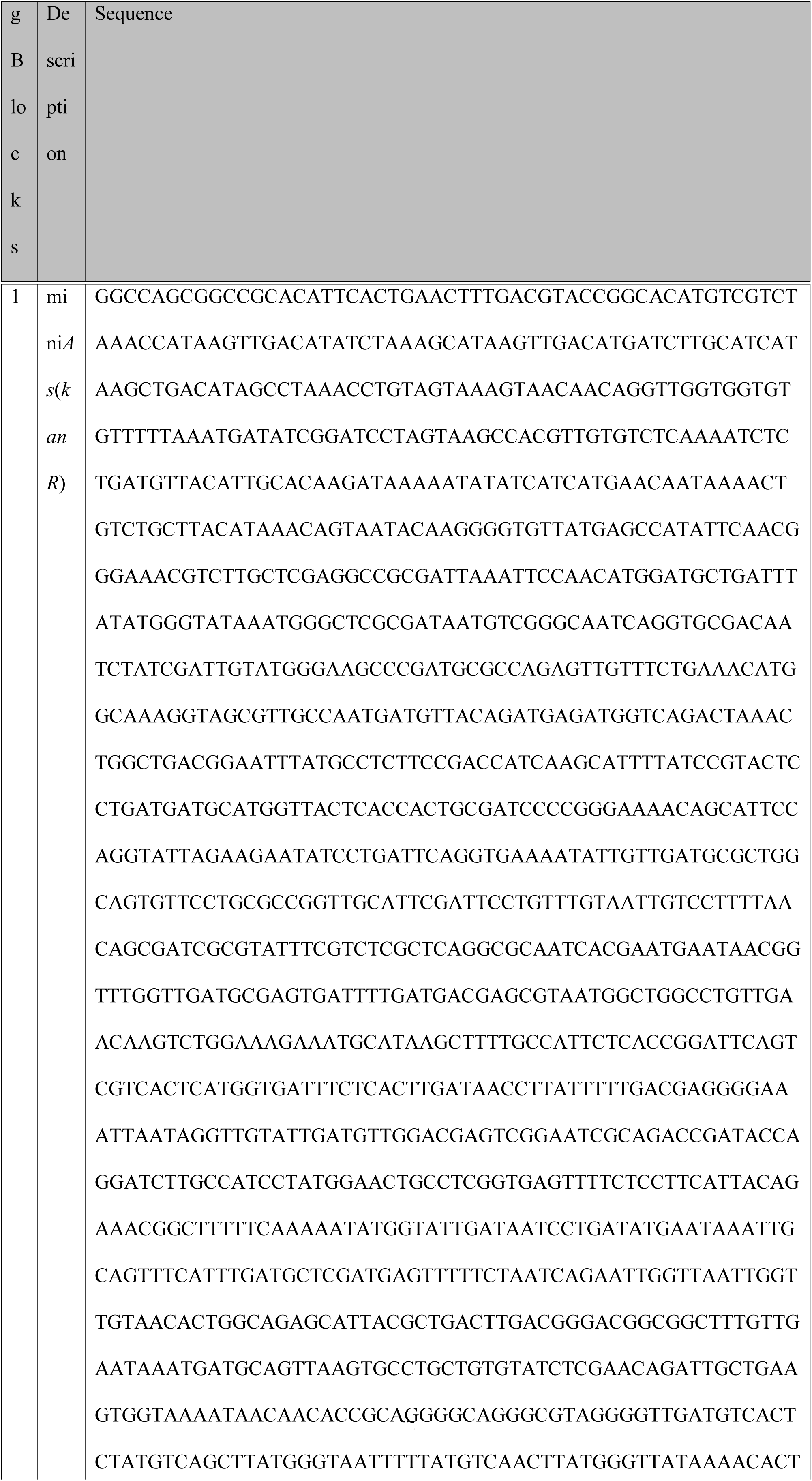
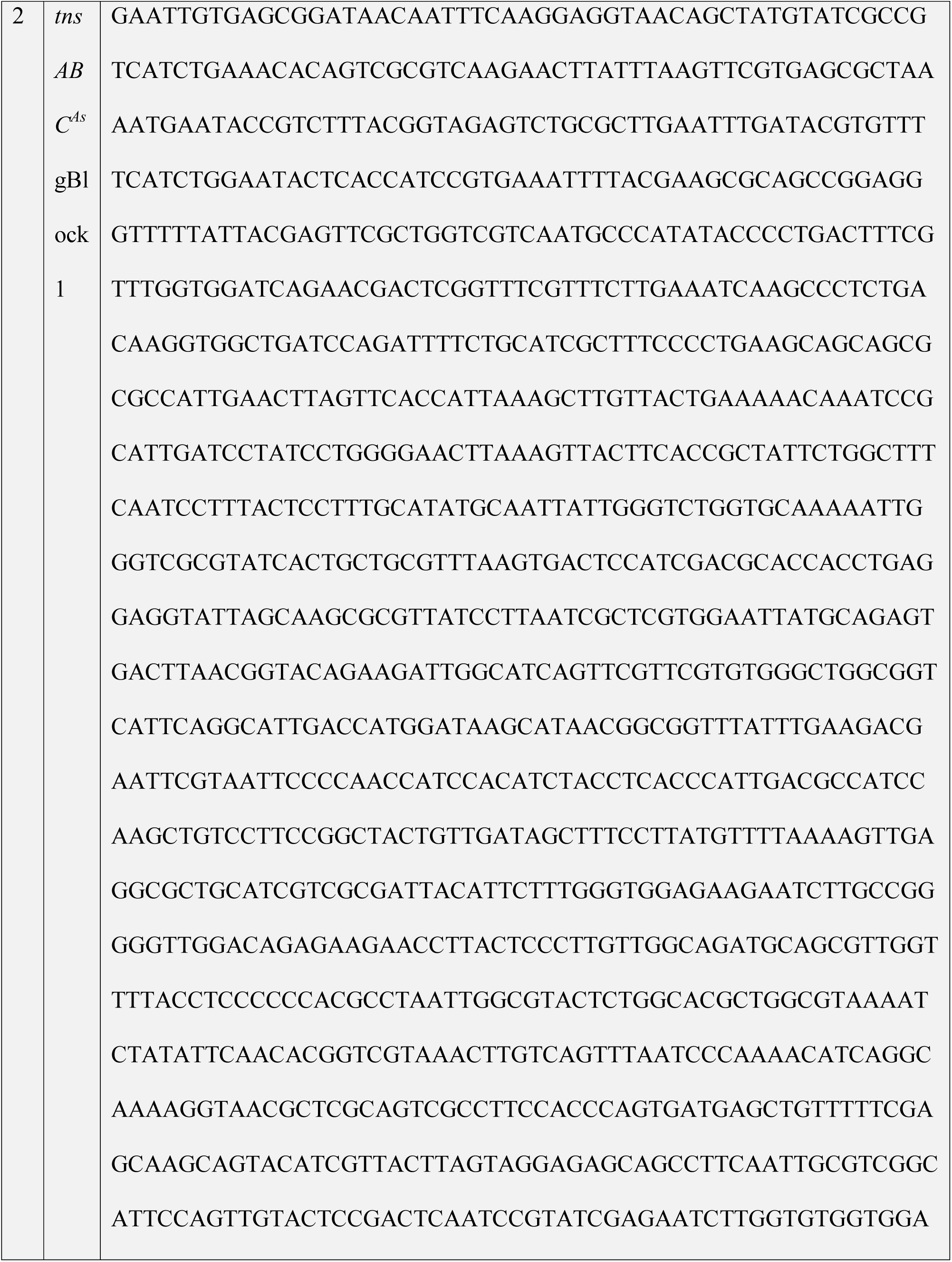
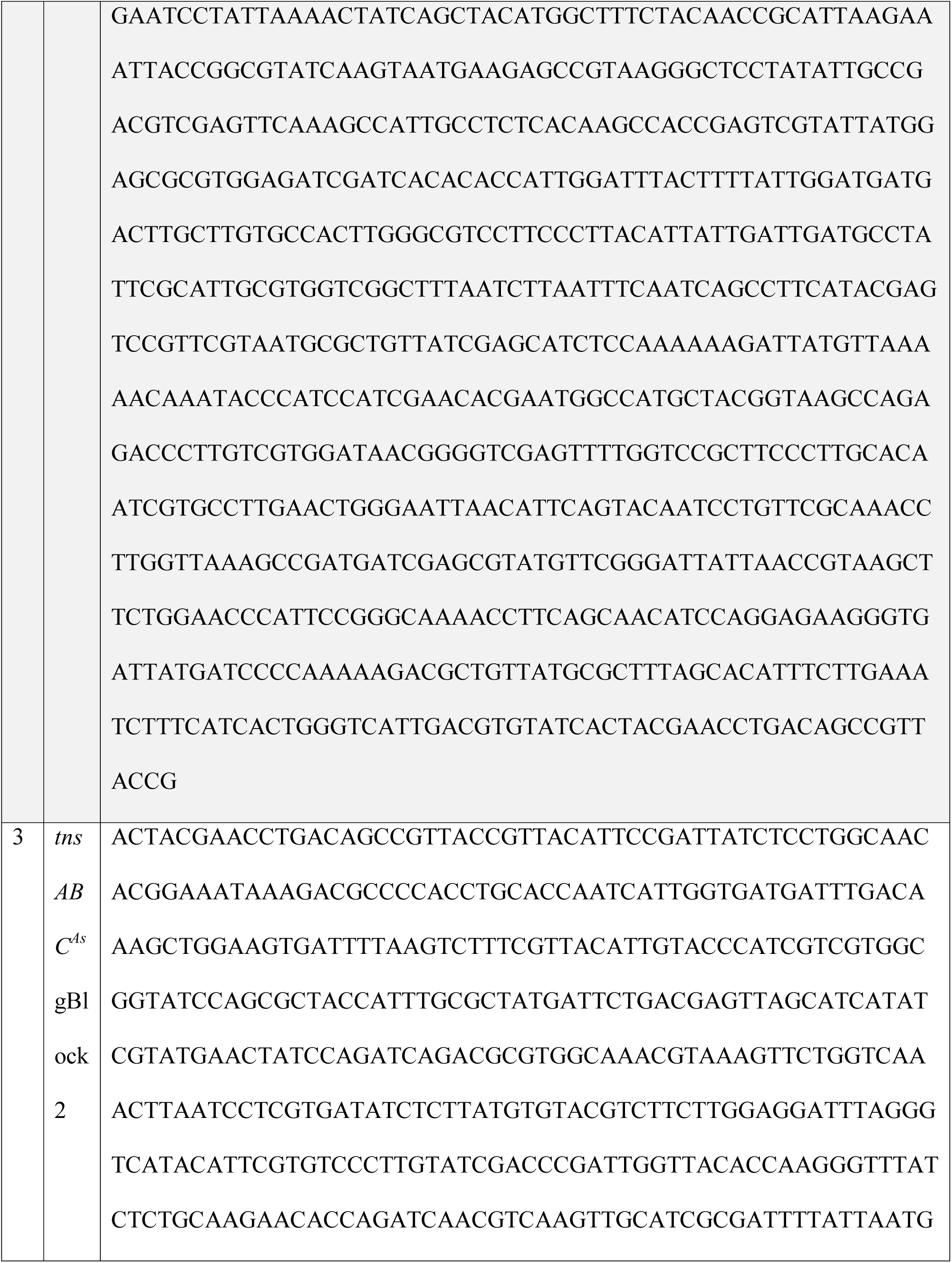
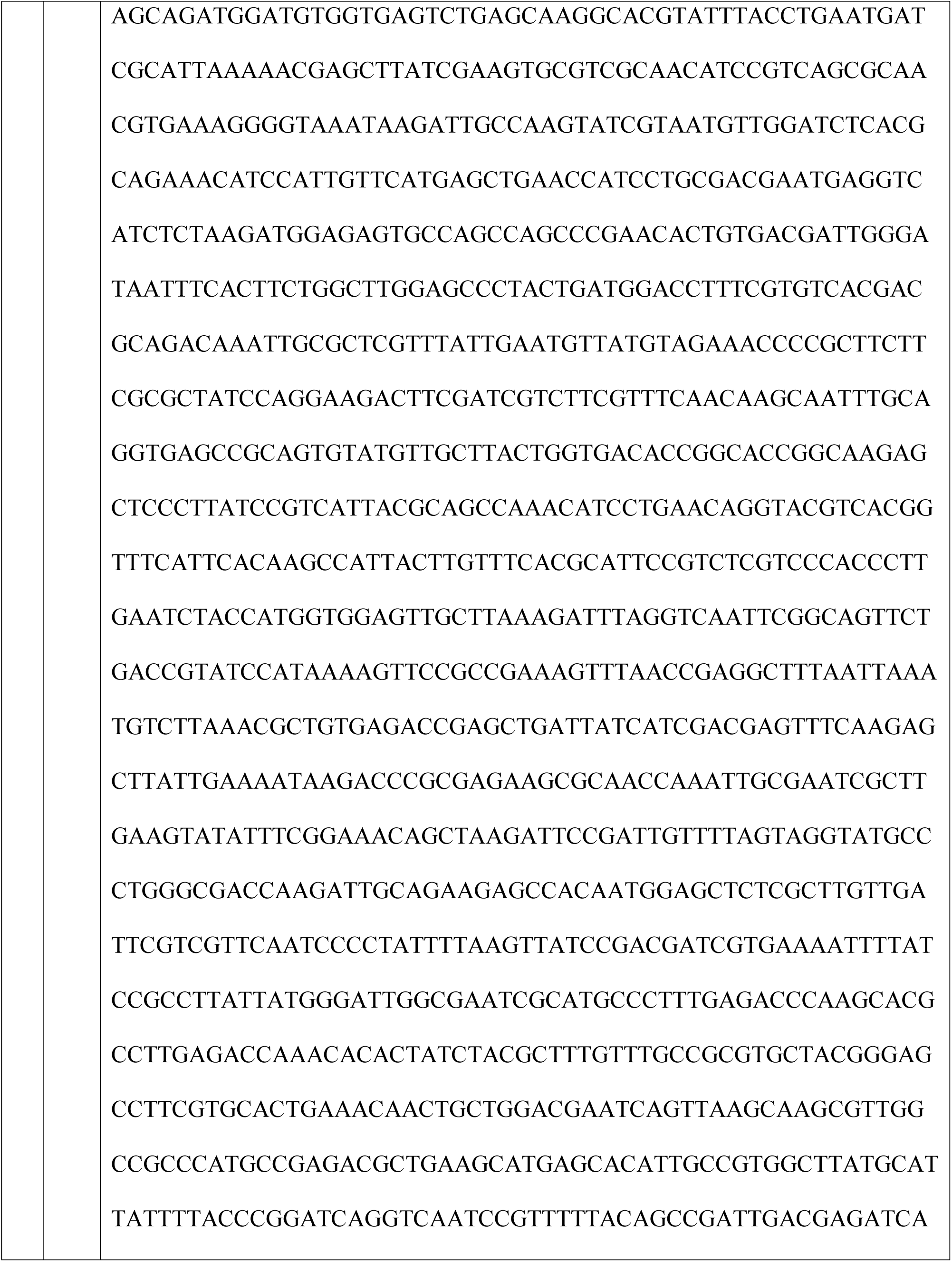
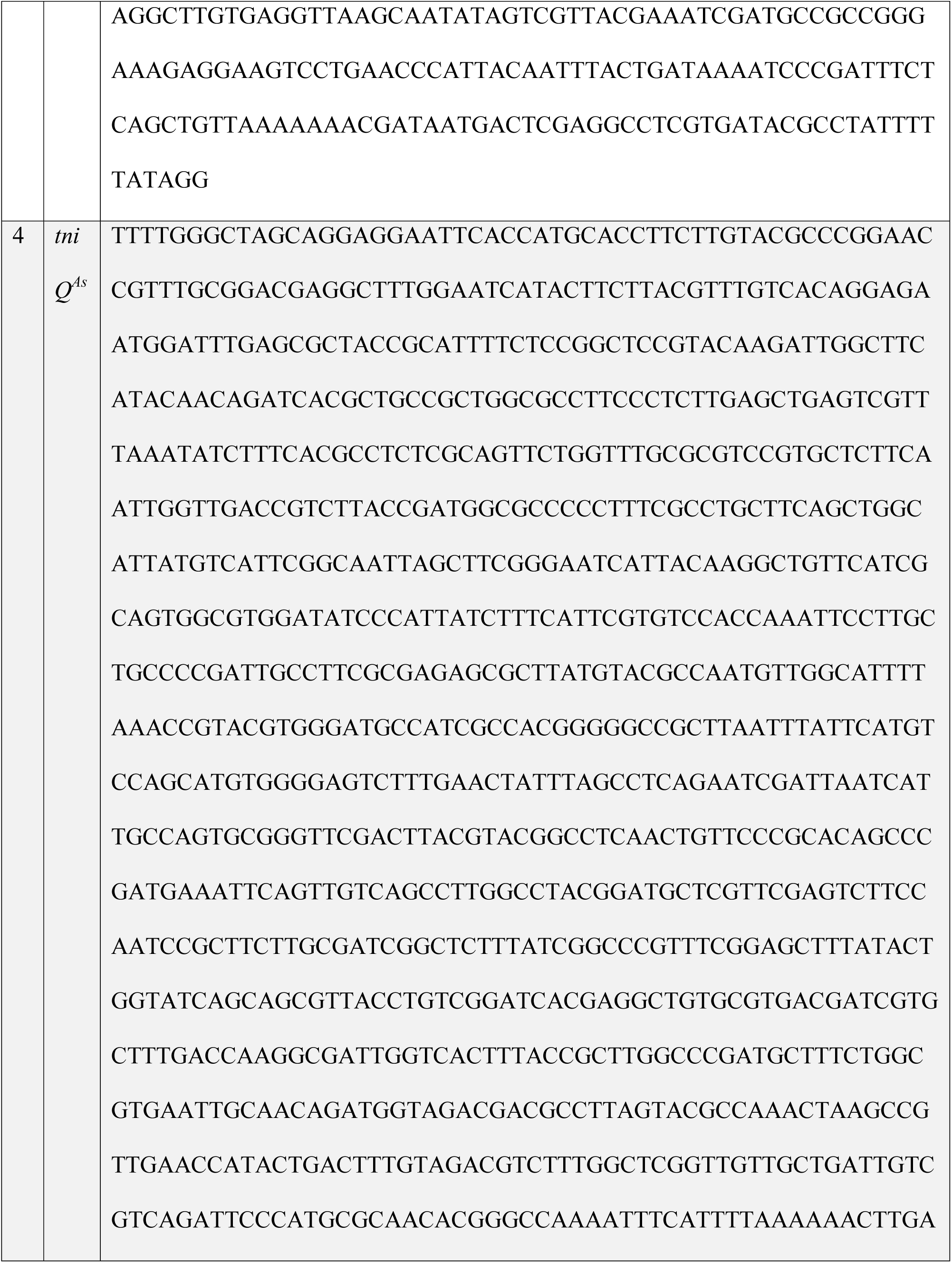
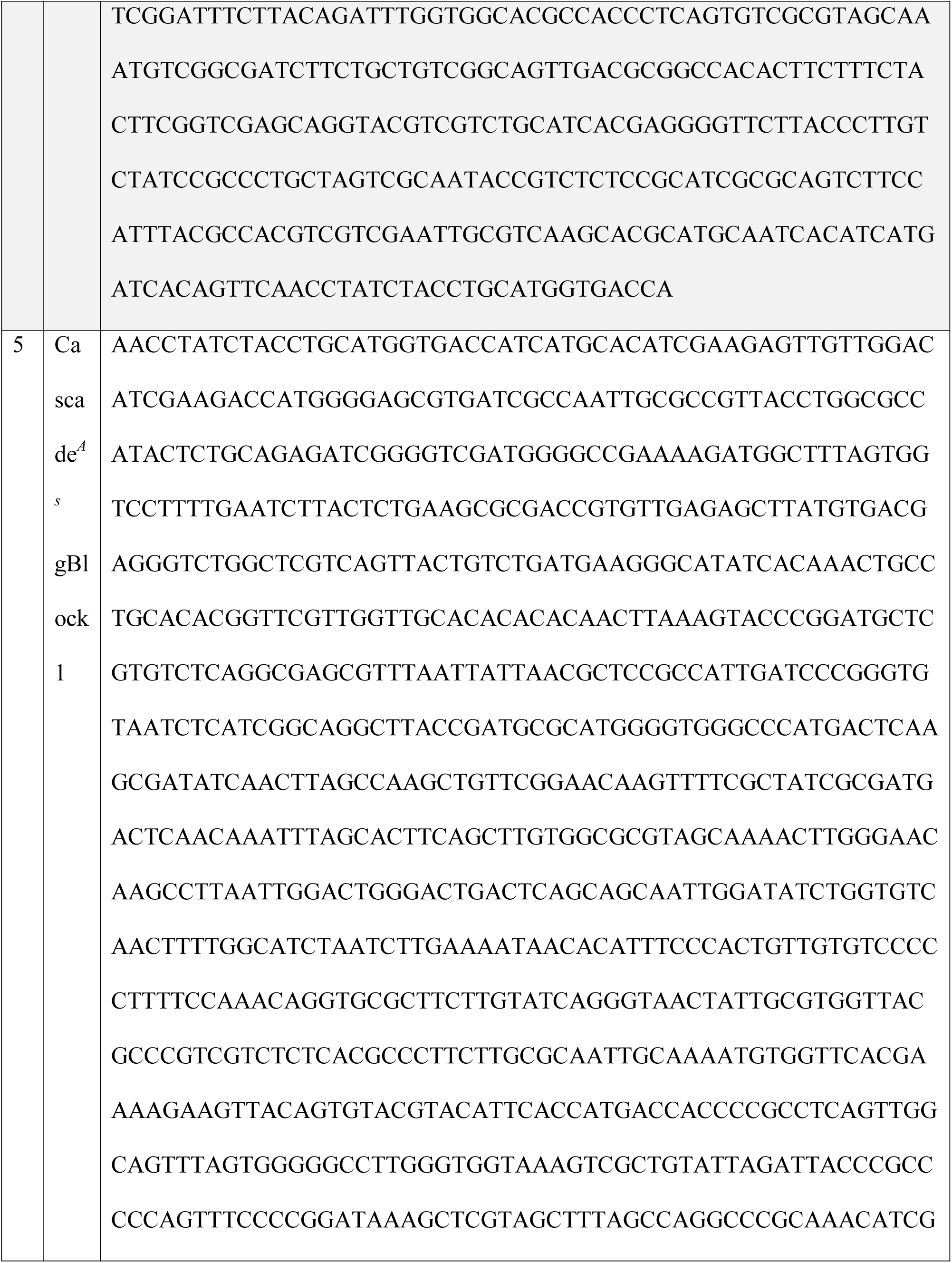
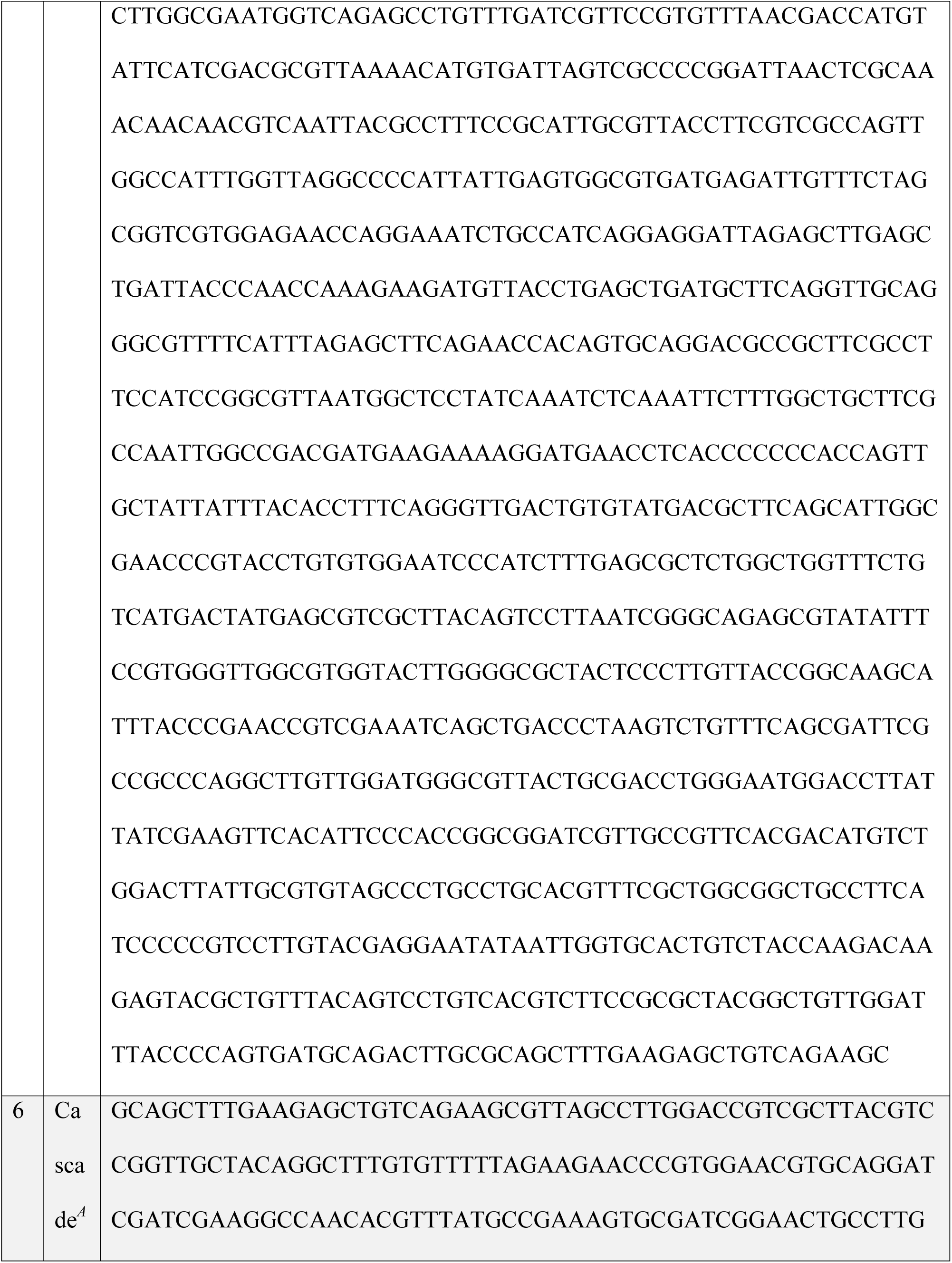
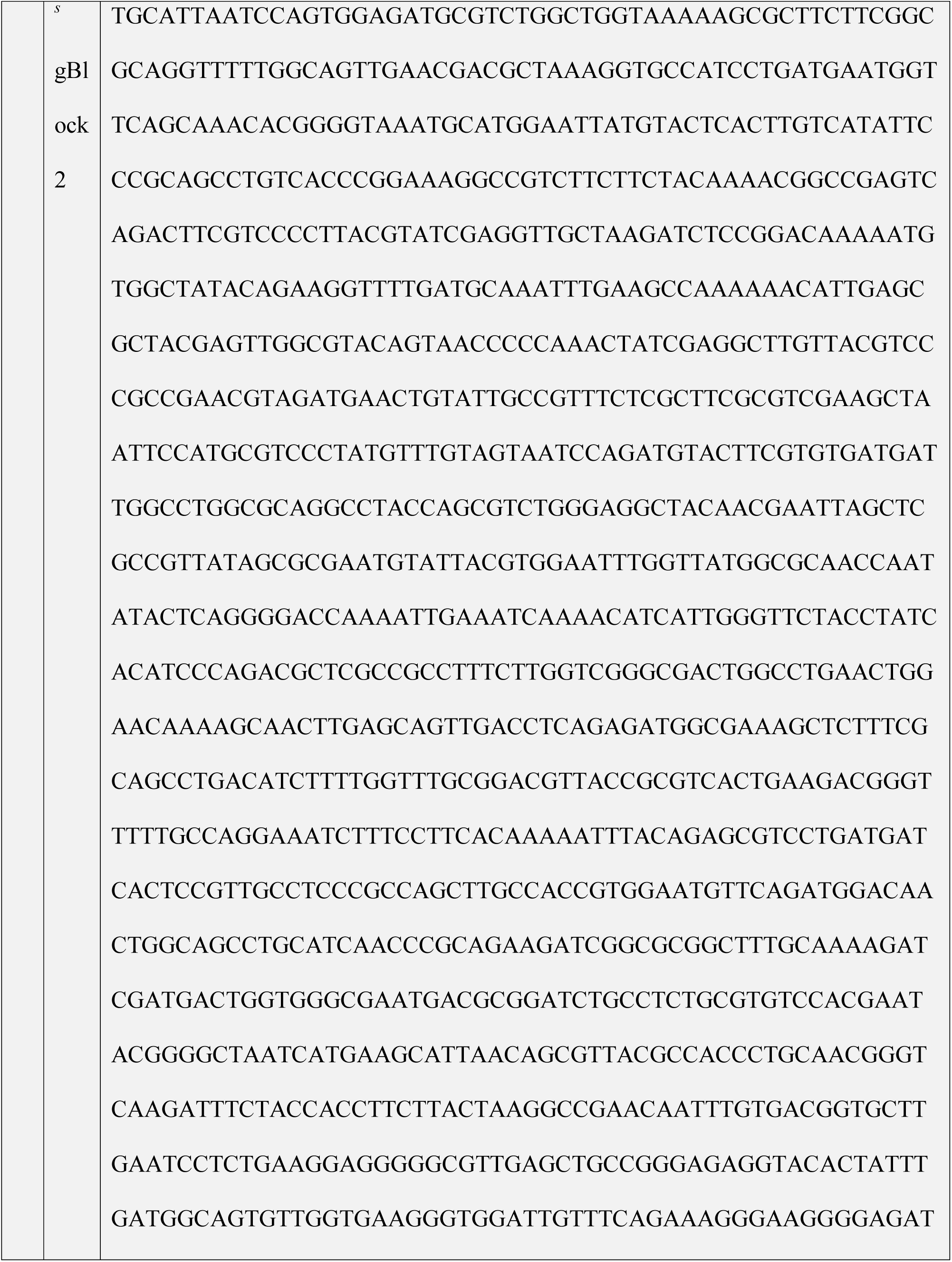
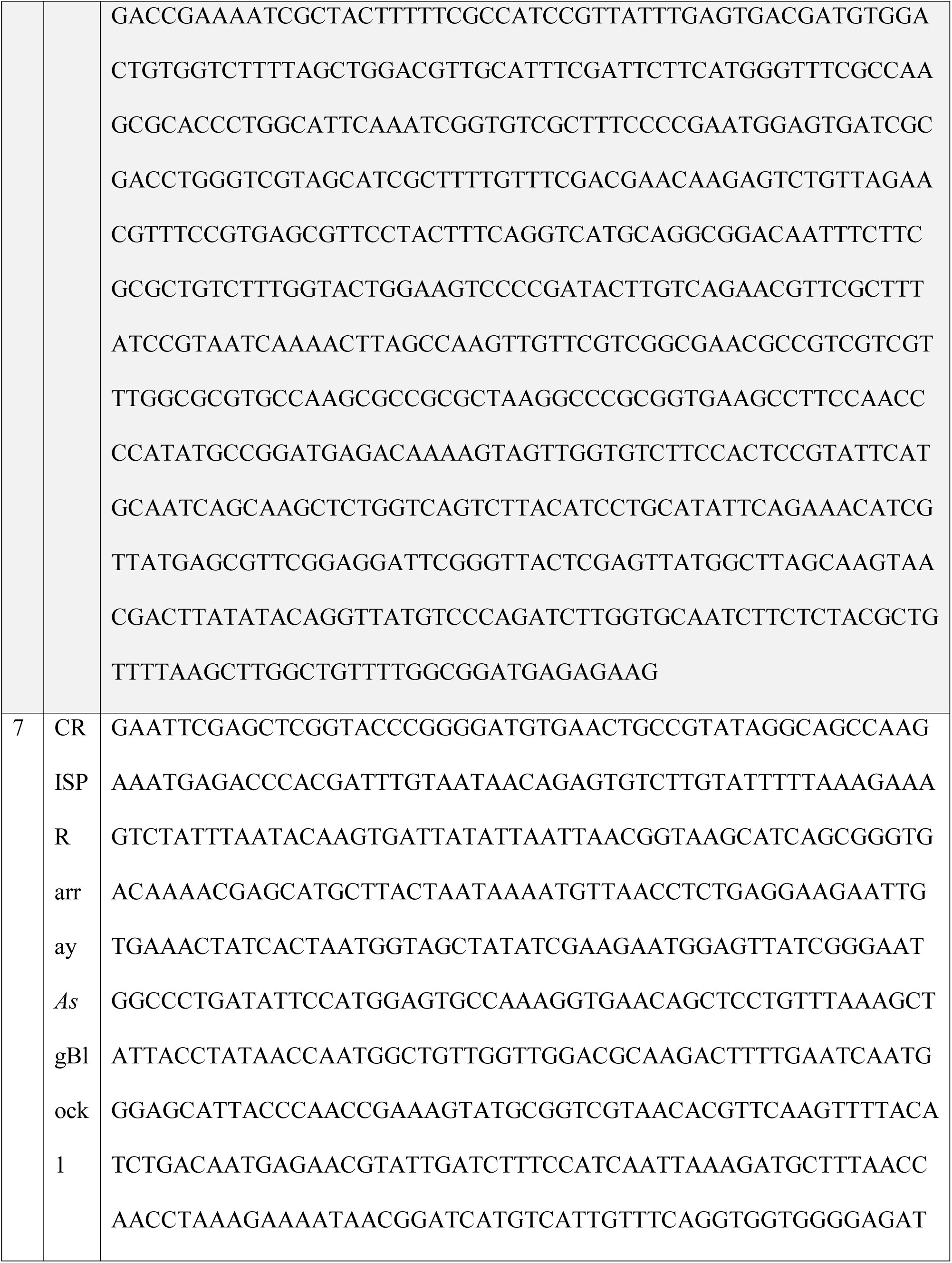
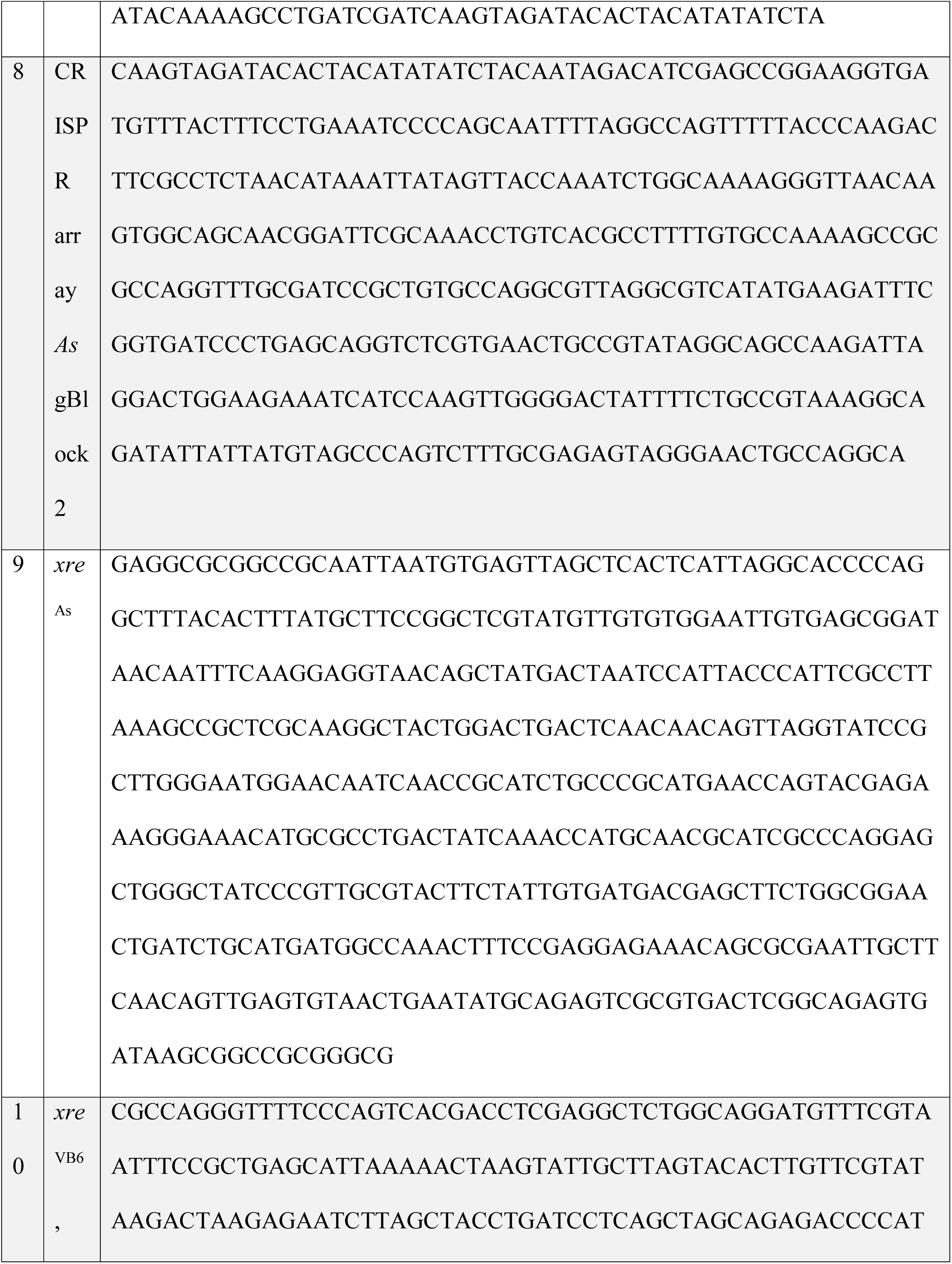
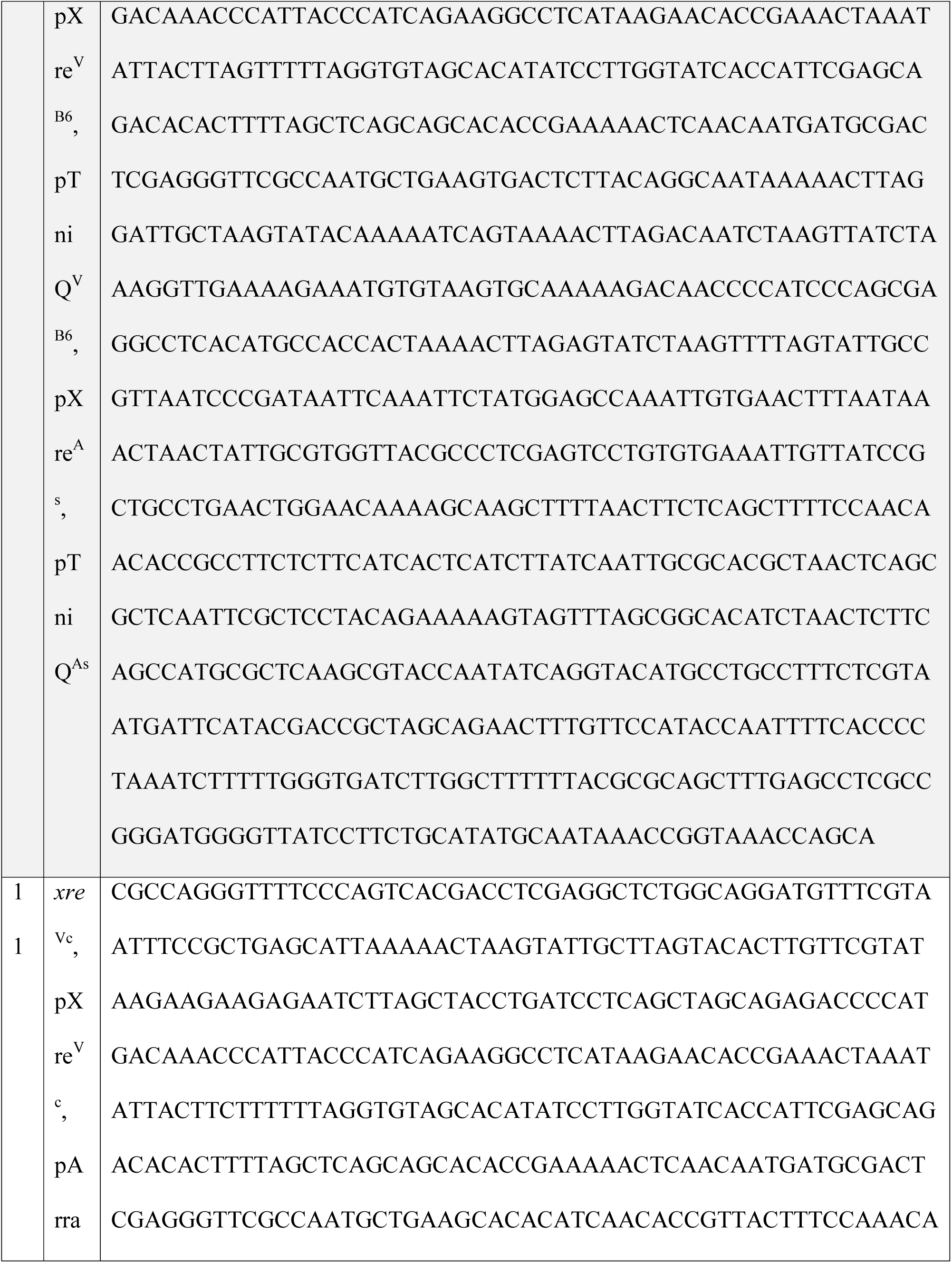
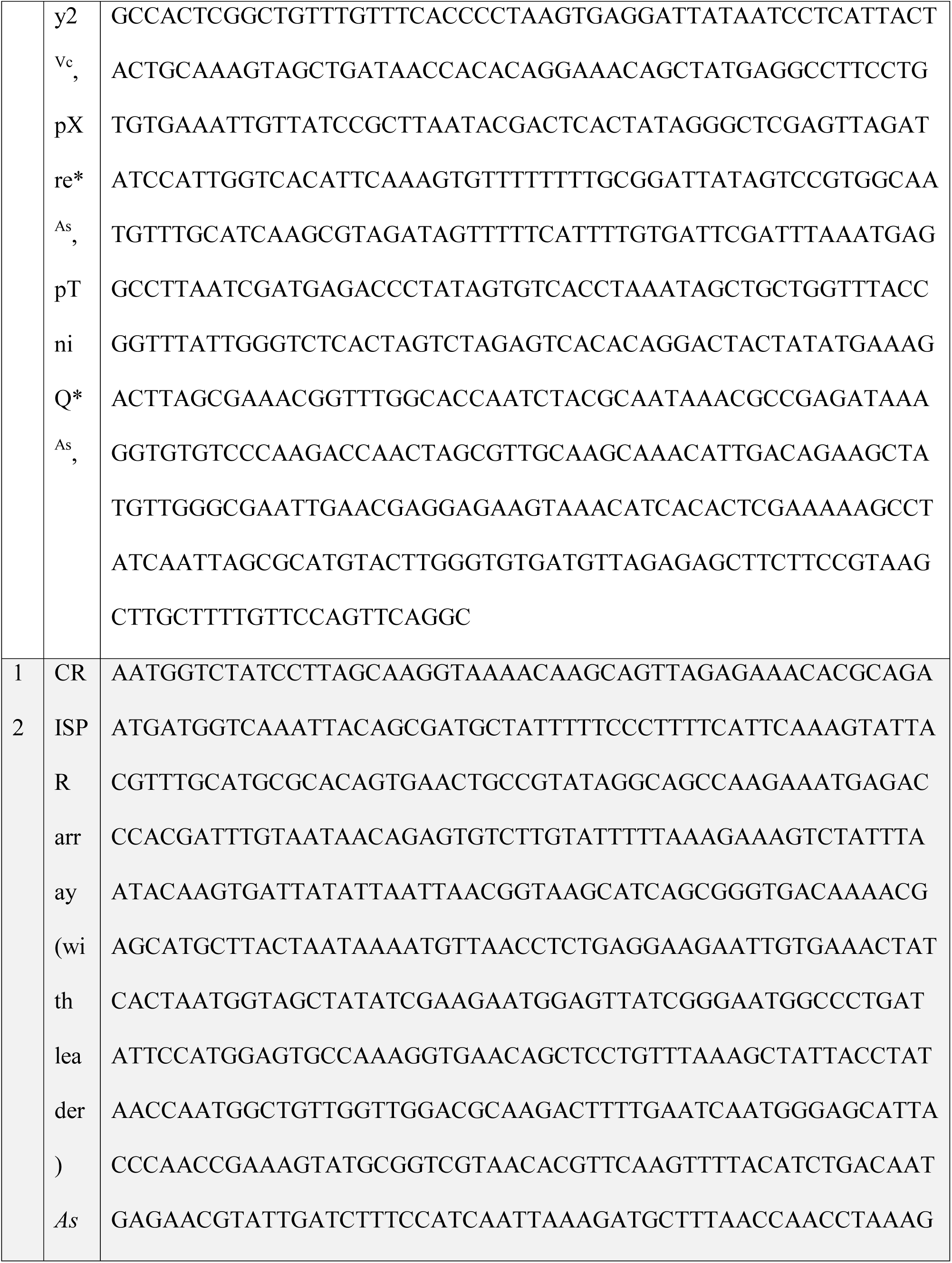
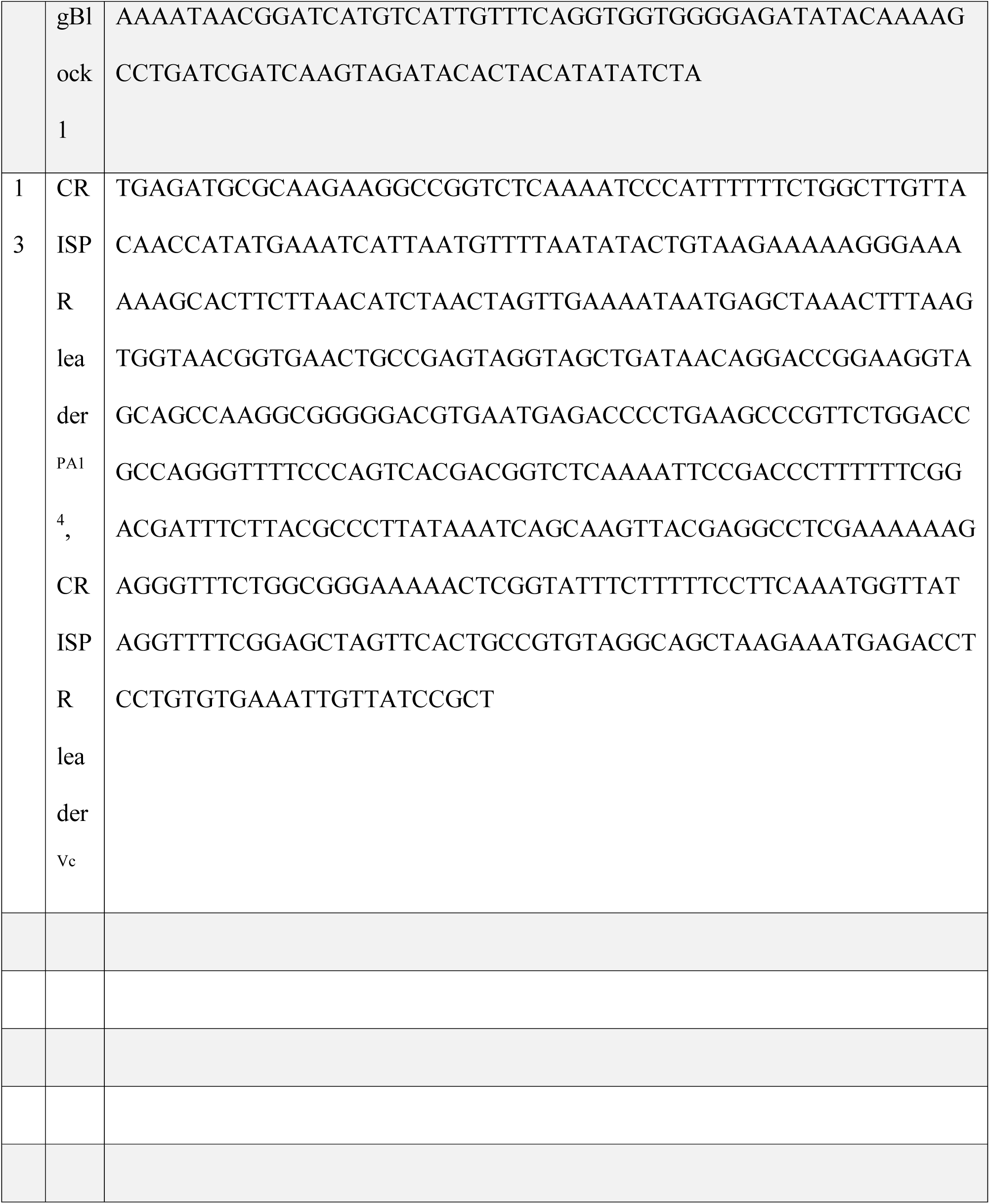

pMTP112 was constructed by ligating gBlock1 into the NotI site of pMS26 following digestion with NotI. The clone used has *A. salmonicida* left end proximal to the Tn7 right end. pMTP113 was constructed by assembling two PCR products amplified from pSL0527 (pDonor) (JEP1858+JEP1859 and JEP1860+JEP1861), one PCR product amplified from gBlock1 (JEP1862+JEP1863) and pMS26 digested with NotI using NEBuilder Hifi (NEB). pMTP114 was constructed by assembling two PCR products amplified from F plasmid (JEP1398+1340 and JEP1341+1399, GenBank: AP001918.1), one PCR product amplified from pMTP150 (JEP1343+JEP1344), one PCR product amplified from pBAD322S (JEP1345+JEP1346, GenBank: DQ131584.1) and pTSC29 digested with EcoRV using NEBuilder Hifi. pMTP115 was constructed by inserting a PCR product amplified from EMG2 (JEP1663+JEP1664, GenBank: U00096.3) into pMTP114 following digestion with BsaI using golden gate cloning (Engler et al., 2008). pMTP116 was constructed by inserting annealed oligos (JEP1485+JEP1486) into pMTP114 following digestion with BsaI using golden gate cloning. pMTP117 was constructed by inserting annealed and extended oligos (JEP1481+JEP1482) into pMP114 following digestion with BsaI using golden gate cloning. pMTP118 was constructed by inserting annealed and extended oligos (JEP1878+JEP1879) into pMTP114 following digestion with BsaI using golden gate cloning. pMTP130 was constructed by assembling gBlock2, gBlock3 and a PCR product amplified from pTA106 (JEP1146+JEP1467) digested by DraII with 3,800 bp fragment gel purified using NEBuilder Hifi. pMTP140 was constructed by assembling gBlock4, gBlock5, gBlock6 and pBAD322G digested with NcoI and HindIII using NEBuilder Hifi. pMTP150 was constructed by assembling two PCR products amplified from pBAD33 (JEP1766+JEP1767 and JEP1768+JEP1769) with gBlock7 and gBlock8 using NEBuilder Hifi.pMTP151 was constructed by inserting annealed oligos (JEP1477+JEP1478) into pMTP150 following digestion with BsaI using golden gate cloning. pMTP160 was constructed by assembling two PCR products amplified from pBAD33 (JEP1766+JEP1767 and JEP1768+JEP1769) with gBlock7 and one PCR product amplified from gBlock8 (JEP1475+JEP1773) using NEBuilder Hifi. pMTP161-165 were constructed by ligating annealed oligos (JEP1477+JEP1478, pMTP161; JEP1776+JEP1777, pMTP162; JEP1778+JEP1779, pMTP163; JEP1669+JEP1670, pMTP164; JEP1671+JEP1672, pMTP165). pMTP170 was constructed by assembling two PCR products amplified from pBAD33 (JEP1766+JEP1767 and JEP1770+JEP1769) with one PCR product amplified from gBlock7 (JEP1774+JEP1474) and one PCR product amplified from gBlock8 (JEP1475+JEP1775) using NEBuilder Hifi. pMTP171-183 were constructed by inserting annealed oligos (JEP1784+JEP1785, pMTP171; JEP1780+1781, pMTP172; JEP1782+JEP1783, pMTP173; JEP1794+JEP1795, pMTP174; JEP1796+JEP1797, pMTP175; JEP1786+JEP1787, pMTP176; JEP1788+JEP1789, pMTP177; JEP1798+JEP1799, pMTP178; JEP1800+JEP1801, pMTP179; JEP1808+JEP1809, pMTP180; JEP1810+JEP1811, pMTP181; JEP1816+JEP1817, pMTP182; JEP1818+JEP1819, pMTP183) into pMTP170 following digestion with BsaI using golden gate cloning. pMTP190 was constructed by assembling two PCR product amplified from pBAD33 (JEP1766+JEP1767 and JEP1771+JEP1769) using NEBuilder Hifi.pMTP191 and pMTP192 were constructed by annealing four oligos (JEP1928, JEP1929, JEP1930, JEP1931 : pMTP191; JEP1932, JEP1933, JEP1934, JEP1935 : pMTP192) and ligating with pMTP190 digested with XmaI and BsaI.

pMTP230 was constructed by assembling one PCR product amplified from pBAD33 (JEP1864+JEP1865), one PCR product amplified from pMTP130 (JEP1866+JEP1867) and pSL0284 digested with NcoI and PflFI with 3,707bp fragment gel purified using NEBuilder Hifi. pMTP240 was constructed by assembling a PCR product amplified from pBAD322 (JEP1868+JEP1869) with pSL0284 digested with NdeI and BglI with 5,152bp fragment gel purified using NEBuilder Hifi. pMTP250 was constructed by assembling a PCR product amplified from pCDFDuet-1 (JEP1838+JEP1839), a PCR product amplified from pBAD322 (JEP1834+JEP1835) and a PCR product amplified from pBBR1MCS-3 (JEP1836+JEP1837) using NEBuilder Hifi. pMTP260 and pMTP270 were constructed by annealing four oligos (JEP1870, JEP1871, JEP1872, JEP1873 : pMTP260; JEP1908, JEP1909, JEP1910, JEP1911 : pMTP270) and ligating with pMTP250 digested with XmaI and BsaI. pMTP261-264 were constructed by inserting annealed oligos (JEP1914+JEP1915, pMTP161; JEP1912+JEP1913, pMTP162; JEP1880+JEP1881, pMTP163; JEP1882+JEP1883, pMTP164) into pMTP260 following digestion with BsaI using golden gate cloning. pMTP271-274 were constructed by inserting annealed oligos (JEP1914+JEP1919, pMTP271; JEP1912+JEP1917, pMTP272; JEP1880+JEP1916, pMTP273; JEP1882+JEP1917, pMTP27) into pMTP270 following digestion with BsaI using golden gate cloning. pMTP275 and pMTP276 were constructed by annealing four oligos (JEP1920, JEP1921, JEP1922, JEP1923 : pMTP275; JEP1924, JEP1925, JEP1926, JEP1927 : pMTP276) and ligating with pMTP250 digested with XmaI and BsaI.

All F derivatives were made by using recombineering(Datsenko and Wanner, 2000) to replace a large region of plasmid F from strain EMG2 (GenBank: AP001918.1) with PCR fragments amplified from pMTP114 derivatives (JEP1376+1386. pMTP115, FΔ (*finO*-*fxsA*)::*lacZ specR*; pMTP116, FΔ (*finO*-*fxsA*)::*cysH^As^ specR*; pMTP117, FΔ (*finO*-*fxsA*)::*ffs^As^ specR*; pMTP118, FΔ (*finO*-*fxsA*)::*guaC^Vc^ specR*).

pOPO01 was constructed by ligating a PCR product amplified from gBlock9 (JEP1657+JEP1757) digested with NdeI and HindIII into pBAD33 digested with the same enzymes. The resulting construct was digested with NdeI and XbaI and ligated with phosphorylated annealed oligos (JEP1842+JEP1843). pOPO02 was constructed by assembling a PCR product amplified from gBlock10 (JEP1764+JEP1765) with pBAD33 digested with NdeI and HindIII using NEBuilder Hifi. The resulting construct was digested with NdeI and XbaI and ligated with phosphorylated annealed oligos (JEP1842+JEP1843). pOPO03 was constructed by ligating a PCR product amplified from gDNA of *V. parahaemolyticus* RIMD221063 (kindly provided by Tobias Doerr) (JEP1952+JEP1960) digested with NdeI and HindIII and phosphorylated annealed oligos (JEP1842+JEP1843) into pBAD33 digested with NdeI and HindIII. pOPO04 was constructed by ligating a PCR product amplified from gBlock11 (JEP1555+JEP1556) digested with SpeI and HindIII into pBAD33 digested with XbaI and HindIII. pOPO05 was constructed by ligating a PCR product amplified from pBAD24 (JEP1759+JEP1760) digested with BsaI and XhoI with a PCR product amplified from EMG2 (JEP1761+JEP1762) digested with the same enzymes. pOPO06-11, pOPO14, and pOPO15 were constructed by ligating fragments from gBlock10 or gBlock11 digested with XhoI and StuI (gBlock10 : pOPO06-09; gBlock11 : pOPO10, pOPO11, pOPO14, pOPO15) into pOPO05 digested with XhoI and SmaI. pOPO12 and pOPO13 were constructed by ligating PCR products amplified from gDNA of *V. parahaemolyticus* RIMD221063 (JEP1956+JEP1957, pOPO12; JEP1954+JEP1955, pOPO13) digested with XhoI and StuI into pOPO05 digested with XhoI and SmaI. pOPO16-19 were constructed by ligating PCR products amplified (from gBlock9, JEP1675+JEP1758, pMTP016; from gBlock10, JEP1556+JEP1764, pMTP017; from gDNA of *V. parahaemolyticus* RIMD221063, JEP1952+JEP1953, pMTP018; from gBlock11, JEP1950+1951, pMTP019) digested with NdeI and XhoI into pET22b(+) digested with the same enzymes. pOPO21 was constructed by ligating annealed oligos (JEP1906+JEP1907) into a PCR product amplified from pCOLADuet-1 (JEP1902+JEP1903) digested with SapI. pOPO23 was constructed by ligating a PCR product amplified from gBlock12 (JEP1892+JEP1893) digested with BsaI, annealed oligos (JEP1778+JEP1779), annealed oligos (JEP1894+JEP1895), and a PCR product amplified from pCDFDuet-1 (JEP1577+JEP1891) digested with BsaI. pOPO25 was constructed by ligating a PCR product amplified from gBlock12 (JEP1896+JEP1897) digested with BsaI, annealed oligos (JEP1782+JEP1783), annealed oligos (JEP1898+JEP1899), and a PCR product amplified from pCDFDuet-1 (JEP1577+JEP1891) digested with BsaI. pOPO26 was constructed by ligating annealed oligos (JEP1940+JEP1941), annealed oligos (JEP1942+JEP1943), and a PCR product amplified from pCDFDuet-1 (JEP1577+JEP1891) digested with BsaI. pOPO27 was constructed by ligating a PCR product from gBlock13 (JEP1937+JEP1938) digested with BsaI, annealed oligos (JEP1948+JEP1949), and a PCR product amplified from pCDFDuet-1 (JEP1577+JEP1891). pOPO29 was constructed by ligating a PCR product amplified from gBlock13 (JEP81+JEP82) digested with BsaI, annealed oligos (JEP1778+JEP1779), annealed oligos (JEP1946+JEP1947), and a PCR product amplified from pCDFDuet-1 (JEP1577+JEP1891) digested with BsaI. pOPO30 was constructed by assembling a PCR product amplified from pCas1_pCas2/3 (JEP1889+JEP1890) and pACYCDuet-1 digested with NcoI and AvrII using NEBuilder Hifi.

### Identifying type I-F CRISPR-guided Tn7-like transposons

Annotated protein fasta files, genomic sequences and feature tables of gamma proteobacteria were downloaded from National Center for Biotechnology Information (NCBI) FTP site on September 22, 2019. In total, there were 53079 genomes for analysis.

Profile HMMs associated with TnsA (PF08722,PF08721), TnsB (PF00665), TnsC (PF11426,PF05621), TniQ(PF06527), Cas5f(PF09614), Cas6f(PF09618), Cas7f(PF09615) and XRE family proteins(PF01381), which can be downloaded from The European Bioinformatics Institute (EMBL-EBI) Pfam database, were used for detecting homologs with hmmsearch (HMMER3, http://hmmer.org/ (Johnson et al., 2010)).

Candidate proteins were grouped into *tnsABC* operons and *tniQ-cas* operon based on their orientation and proximity. Then each *tnsABC* operon was grouped with its downstream *tniQ-cas* operon into one transposon functional unit. The Xre/HTH (helix turn helix) proteins situated between the two operons and are homologous to restriction controller proteins (blastp, identity >40%) were defined as candidate regulators.

### CRISPR array detection

Manually curated CRISPR repeats of CRISPR-guided transposons were used to create a DNA sequence profile, which was used as a query for nhmmscan searches (Wheeler and Eddy, 2013) to find CRISPR repeats in the downstream 20-kb region of *cas6*. Putative repeats were grouped into arrays by their distances to each other. The distance between repeats was required to be >55 bp and <65 bp, the bit-score threshold is −1. The distance between last repeat and previous repeat was allowed to be between 43 bp and 55 bp, but in such cases its bit-score had to be >=0.3. The sum of bit-scores of repeats in an array cannot be lower than 6.0. The longest non-overlapping arrays are collected as putative CRISPR arrays. All repeats besides the final repeat from the first array downstream of *cas6* were used to create an updated repeat profile, and the CRISPR detection procedure was repeated with the new profile twice.

### Protospacer detection

To detect protospacers that match the transposon-associated CRISPR spacers, each spacer was converted into position-specific scoring matrix (PSSM) and used to search upstream 1-kb DNA of *tnsA* for matches with biopython (threshold=11.0)(Cock et al., 2009). Because every 6th base of spacers is flipped out in type I CRISPR Cascade complex, all 6th positions of the matrix are set to have equal weight on all four bases.

Except for *ffs* (SRP-RNA) gene, the major attachment site genes that containing the candidate protospacers are classified with the annotations provided in NCBI. The attachment site SRP-RNA gene (*ffs*) is often poorly annotated, so it was reannotated using cmsearch (infernal, (Nawrocki and Eddy, 2013)) and SRP-RNA profile (RF00169) available on RFAM (https://rfam.xfam.org/).

### Constructing similarity trees

The TnsA, TniQ and Xre proteins were clustered using the Cd-hit (Li and Godzik, 2006), with identity threshold set to 90%. Multiple alignments of the representatives were done with MUSCLE (Edgar, 2004). Similarity trees were made with FastTree (Price et al., 2009) using WAG evolutionary model and the discrete gamma model with 20 rate categories as previously described (Peters et al., 2017a). The visualization of the trees, major attachment sites, CRISPR arrays and matched spacers was done with ETEToolkit (Huerta-Cepas et al., 2016).

### Searching shared promoter motifs of *xre* and CRISPR-Cas genes

The transposons were classified into two groups based on associated *xre* lengths (68 a.a. for I-F3a or ∼100 a.a. for I-F3b) and similarities to C.AhdI and C.Csp231I. For each group, the 100bp upstream of *xre*, second CRISPR array, and *tniQ-cas* operon were collected and deduplicated with dedupe.sh (BBTools, (Bushnell)) with threshold of 70% identity or 30 edit distance. The sequences were then sent to MEME (Bailey et al., 2009) for motif detection and comparison.

### Comparing consensus CRISPR repeat sequences of chromosome targeting spacers to those of other spacers

To make consensus sequences of CRISPR repeats, the transposon representatives with non-redundant TniQ were selected with Cd-hit and separated into two groups based on their attachment sites being *ffs*/*rsmJ* or *guaC*/*yciA*. The upstream and downstream CRISPR repeats of the chromosome targeting spacers were collected and turned into sequence logo using WebLogo 3 (Crooks et al., 2004) then compared to the sequence log made of other repeats.

### Transposition assays

All transposition assays were performed in MTP1191, or one of MTP997 or MTP1196 with an F plasmid derivative as indicated in Table 2.

For the elements derived from *A. salmonicida* S44, strains used to monitor transposition were made competent by standard chemical methods (Peters, 2007) and transformed with pMTP130, pMTP140, and a derivative of pMTP150, pMTP160, pMTP170, or pMTP190 as indicated in Table 2 onto LB + 100 μg/mL carbenicillin, 10 μg/mL gentamicin, 30 μg/mL chloramphenicol, .2% w/v glucose. After 16 hours incubation at 37°C, several hundred transformants were washed up in M9 maltose and diluted to a calculated OD = 0.2 in M9 maltose + 100 μg/mL carbenicillin, 10 μg/mL gentamicin, 30μg/mL chloramphenicol, .2% w/v arabinose, .1 mM IPTG to induce transposition.

For experiments monitoring transposition frequency through loss of sugar metabolism on MacConkey’s media, induction pools were incubated for 24 hours with shaking at 30°C before being serially diluted and plated on MacConkeys 1% w/v lactose, sorbitol, or galactose. Plates were incubated at 37°C for 16 hours before colonies were counted.

For experiments monitoring transposition frequency by the mate out assay (Supplemental Figure 2a), after 24 hours incubation with shaking at 30°C, a portion of induced cultures were washed once and resuspended in LB + .2% w/v glucose. After 2 hours incubation at 37°C induced pools were mixed with prepared mid-log CW51 recipient strain at a ratio of 1:5 donor:recipient and incubated with gentle agitation for 90 minutes at 37°C to allow mating. After incubation cultures were vortexed, placed on ice, then serially diluted and plated on LB + 20 μg/mL nalidixic acid, 100 μg/mL rifampicin, 100 μg/mL spectinomycin, 50 μg/mL X-gal, with or without 50 μ kanamycin to sample the entire transconjugant population or select for transposition respectively. Plates were incubated at 37°C for 36 hours before colonies were counted.

*V. cholerae* Tn6677 transposition assays were performed as above with function plasmids pMTP230, pMTP240, and a derivative of pMTP250, pMTP260, or pMTP270 as indicated in Table 2 with the exception of 8 μg/mL tetracycline replacing gentamicin when present to accommodate the different vector set.

In all experiments, non-target controls where the spacer did not match the target F plasmid were used, with transposition frequency similar to non-target rate in Figure 3B for *A. salmonicida* S44 transposition, or Figure 4D for Tn6677 transposition.

### *P. aeruginosa* CRISPR interference assays

All interference assays were performed in BL21-AI. BL21-AI was transformed with pOPO30, pCsy complex, and a derivative of pCOLADuet-1 as indicated in Table 2 under repressing conditions. Overnight cultures were diluted 1:50 in LB + 100 μ g/mL carbenicillin, 50 μg/mL kanamycin, and 30 μg/mL chloramphenicol and grown for one hour before addition of IPTG and arabinose to a final concentration of .1 mM and .2% w/v respectively. Cultures were grown to OD = 0.6 before electrocompetent cells were prepared by standard methods (Peters, 2007) and transformed with various dilutions of pOPO21 or pCOLADuet-1. Cells were recovered in SOB for one hour before being plated on LB + 100 μg/mL carbenicillin, 50 μg/mL kanamycin, 30 μg/mL chloramphenicol, and 100 μg/mL spectinomycin. Plates were incubated at 37°C for 16 hours before colonies were counted.

### Xre protein purification

pOPO16, pOPO17, pOPO18 or pOPO19 were transformed into BL21 (DE3), which was cultured in Terrific Broth at 37°C and induced with 0.1mM IPTG during log-phase. Cells were cultured an additional 12-16 hours at 18°C before being collected with centrifugation and lysed by sonication in nickel buffer (20 mM HEPES–NaOH (pH 7.5), 500 mM NaCl, 30 mM imidazole, 5% (v/v) glycerol, 5 mM β-mercaptoethanol) supplemented with 0.15 mg/mL lysozyme. Lysate was cleared by centrifugation and loaded on Nickel-NTA column, washed with nickel buffer, and eluted over a 30 mM to 500 mM imidazole gradient. Selected purified fractions were pooled, dialyzed and buffer exchanged into storage buffer (20 mM HEPES–NaOH (pH 7.5), 100 mM KCl, 5% (v/v) glycerol, 1mM DTT). The purified proteins were snap-frozen with liquid nitrogen and stored at −80°C.

### Electrophoretic mobility shift assay (EMSA)

The promoter fragments of putative Xre regulated genes and their mutated variants were PCR amplified and purified. 100 nM DNA was incubated with different amounts of purified Xre proteins in equilibrium buffer (50 mM Tris–HCl (pH 8.0), 1 mM DTT, 10 mM MgCl2) at 25℃ for 20min then mixed with glycerol (final concentration 6%). EMSAs were performed in 6% non-denaturing TBE PAGE (Polyacrylamide gel) with 0.5x TBE as running buffer, running at 80V for one hour at room temperature. The gels were EtBr stained and visualized with UV imager.

DNA substrates were produced as follows: ArapBAD was amplified from pBAD24 (JEP175+JEP1364), pXre(Vp) and pArray2(Vp) were amplified from *V. parahaemolyticus* RIMD221063 (JEP1956+JEP1957, pXre(Vp); JEP1954+JEP1955, pArray2(Vp)), pXre(Vc) and pArray2(Vc) were amplified from gBlock11 (JEP29+JEP30, pXre(Vc); JEP1553+JEP82, pArray2(Vc)), pXre(As) was amplified from pOPO08 (JEP1321+JEP81), pTniQ(As) was amplified from pOPO09 (JEP1322+JEP81), pXre*(As) was amplified from pOPO10 (JEP1321+JEP81)., pTniQ*(As) was amplified from pOPO11 (JEP1322+JEP81)., pXre(VB6) was amplified from pOPO06 (JEP1553+JEP81), and pTniQ(VB6) was amplified from pOPO07 (JEP1554+JEP81).

### In vivo promoter assay

To confirm and characterize the putative Xre-mediated regulation, the promoters of putative Xre-regulated genes were transcriptionally or translationally fused to a plasmid-encoded *lacZ* reporter gene. pOPO01, pOPO02, pOPO03, or pOPO04 and a derivative of pOPO05 as indicated in Table 2 were transformed into *E. coli* BW27783, Overnight cultures of transformed cells were diluted 100-times into LB broth supplied with different concentrations of sugar, and cultured for additional 20 hours at 30℃. The LacZ activities were measured with standard miller unit assay (Malke, 1993).

