## Supplemental Information for "Guide RNA categorization enables target site choice in Tn7-CRISPR-Cas transposons"

Michael T. Petassi, Shan-Chi Hsieh, and Joseph E. Peters
Department of Microbiology, Cornell University, Ithaca, NY 14853, USA

**Supplemental Table S1 - Type I-F3 Tn7-CRISPR-Cas elements in gammaproteobacteria**

List of 801 I-F3 Tn7-CRISPR-Cas elements indicated by the attachment site where located indicating the host genus, species, and strain with accession information (contig). Sequences were manually curated to identify the repeats and spacers. Repeats were scored by similarity to consensus sequence of non-terminal repeats of the first array. The spacer and spacer match (protospacer) is indicated with the size of the spacer and number of predicted base-pair contacts and where the matching spacer resided in the array. It is indicated if another I-F CRISPR-Cas system could be identified in the strain. See text for details.

**Supplemental Figure S1 – *Aeromonas* element features and transposition**

(a) Schematic representation of two nearly-identical I-F3b Tn7-CRISPR-Cas elements found in different bacterial species suggesting recent activity. Core features are indicated as in Figure 2a. Elements are located either in the chromosomal *ffs* site in *A. hydrophila* AFG_SD03 or inserted into a phosphoadenosine phosphosulfate reductase gene (*cysH*) found on a large conjugal plasmid (pS44-1) in *A. salmonicida* S44. The *A. hydrophila* element is split across several contigs interrupted by apparent IS element insertions.

(b) Spacers in the leader-proximal position of *A. salmonicida* S44 and *A. hydrophila* AFG_SD03 CRISPR arrays match protospacers in a plasmid encoded *cysH*. Relative position of the protospacers are indicated. Distance from the edge of the protospacer matching *A.salmonicida* S44 spacer to the central position in the target site duplication (TSD), the 5 bp TSD (underlined), as well as the terminal sequence of the transposon ends are shown.

(c) Repeats and spacers from Tn7-like CRISPR arrays in *A. salmonicida* S44 and *A. hydrophila* AFG_SD03. Repeats are annotated as in Figure 2c. Differences from the first repeat are indicated in red. Matches between the guide RNA and protospacer are indicated by a short vertical line. The putative I-F PAM is underlined.

(d) Protospacers on the chromosome or F plasmid are targeted at high efficiency with atypical guide RNA complexes. The same three *lacZ* guide RNA complexes were tested with either the F::*lacZ* plasmid or chromosome (*lacZ* in its native position) and insertion events were indicated by generating white verses red colonies on MacConkey's lactose indicator media. Graph shows the mean +/- standard deviation of three biological replicates and number of white colonies and total colonies observed.

(e) Different genes on the chromosome can be targeted for atypical guide RNA-directed transposition in the *E. coli* chromosome. Two genes where tested with two spacers each (top and bottom strand) for galactose (*galK*) and sorbitol (*srlD*). Transposition frequency was assayed by monitoring gene inactivation leading to loss of sugar catabolism as visualized by white verses red colonies on the appropriate MacConkey's indicator media. Graph shows the mean +/- standard deviation of three biological replicates.

**Supplemental Figure S2 – Assaying full transposition frequency and position**

(a) Mate out assay schematic. Target DNAs with the appropriate protospacer are recombined onto an F plasmid and transposition genes and arrays are supplied on expression vectors to mobilize a mini-Tn donor element located in the chromosome (Experimental procedures). After induction, transposition frequency is determined by mating out the population of F plasmids into a donor strain and quantifying antibiotic marker presence in transconjugants as shown.

(b) Transposition position and orientation in transconjugants are determined by PCR. An internal primer and two primers flanking the target site capture orientation of insertion.

(c) Transposition position and target site duplication are confirmed by Sanger sequencing for *A. salmonicida* S44 transposition. Arrows indicate position of the central base of the target site duplication for isolated transposition events targeting pS44-1, with distance from protospacer to the central position of the target site duplication listed for eight transposition events, confirming previously described target site wobble. Graph shows one representative of the actual target site duplication (TSD).

**Supplemental Figure S3 – Interference assay repeat sequences**

Repeat sequences used in interference assays. Differences from *P. aeruginosa* repeats are indicated in red. The orange box indicates the previously established conserved region that comprises the putative stem-loop in I-F repeats.  N32 indicates the position encoded in the spacer.

**Supplemental Figure S4 – I-F3 Xre cluster into two clades with restriction-modification C proteins**

(a) Similarity tree of Xre with associated C proteins C.*Ahd*I and C.*Csp*231I (marked in teal and fuchsia), indicating clustering in the two branches. Features are indicated as in Figure 1.

(b) Predicted regulator sequences for Xre and associated C proteins. Conserved inverted motif sequences are indicated by bold red text and black arrows. The start codon of the downstream gene is underlined, except for pArray2 sequences, where the first three bases of the *att*-targeting spacer are underlined.

**Supplemental Figure S5 – Elements with shortened spacers and their insertion positions**

(a) *ffs*-integrated elements,

(b) *araC*-like integrated elements. Features are indicated as in Figure 2b. Arrays are shown for each element, with repeat sequences marked in blue, spacers marked in red, and conserved Cas6 binding motifs are underlined.

**Supplemental Figure S6 – Schematic representation of elements inserted downstream of *parE***

Similarity tree of TniQ proteins indicates that *Parashewanella curva* C51 has representatives that group with elements that target the *parE att* site and elements that use the I-F3 CRISPR-Cas system. In cases where two TniQs are found in the element, the one used for the similarity tree is indicated with orange highlighting.

#### Supplemental Figure S1 – *Aeromonas* element features and transposition

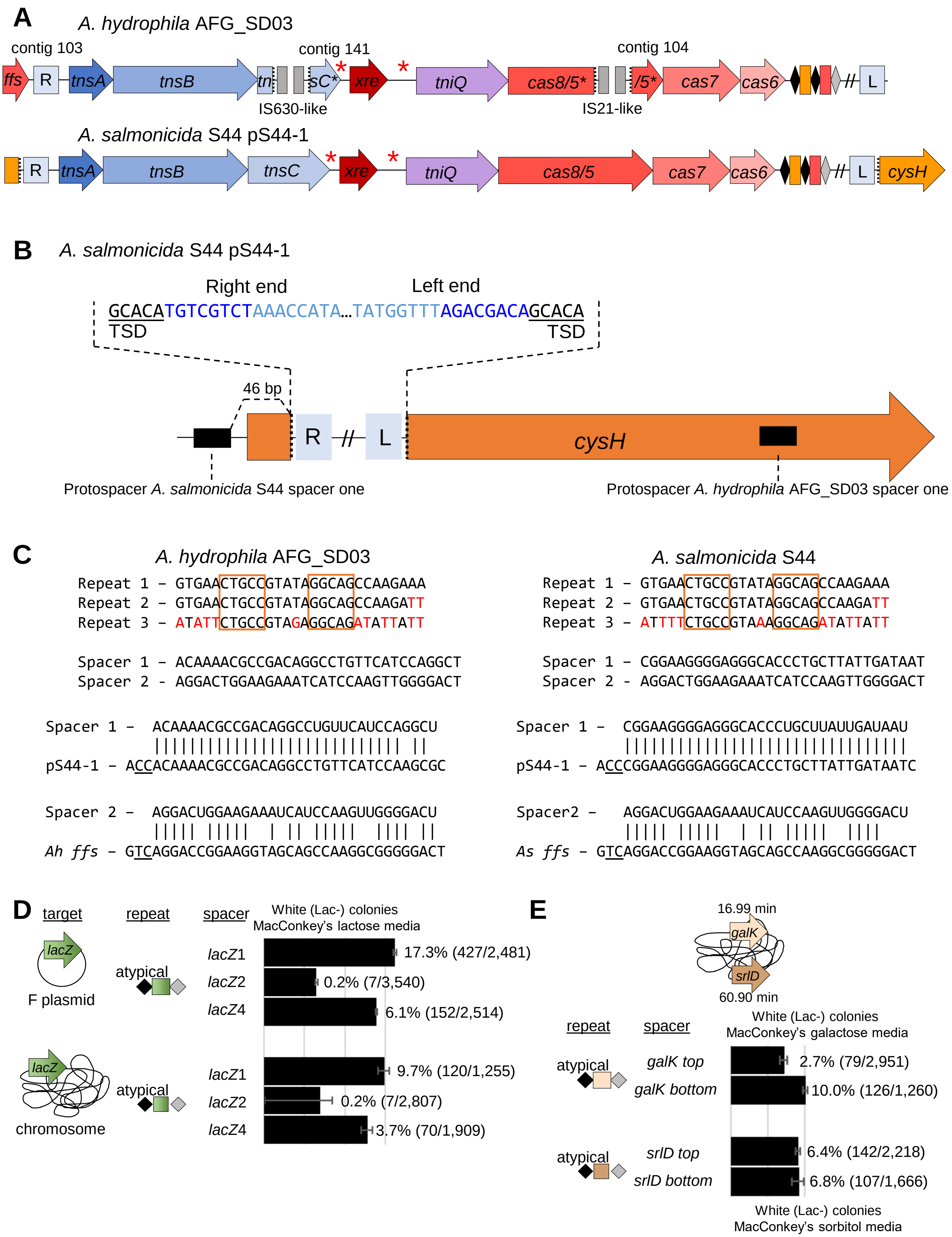

#### Supplemental Figure S2 – Assaying full transposition frequency and position

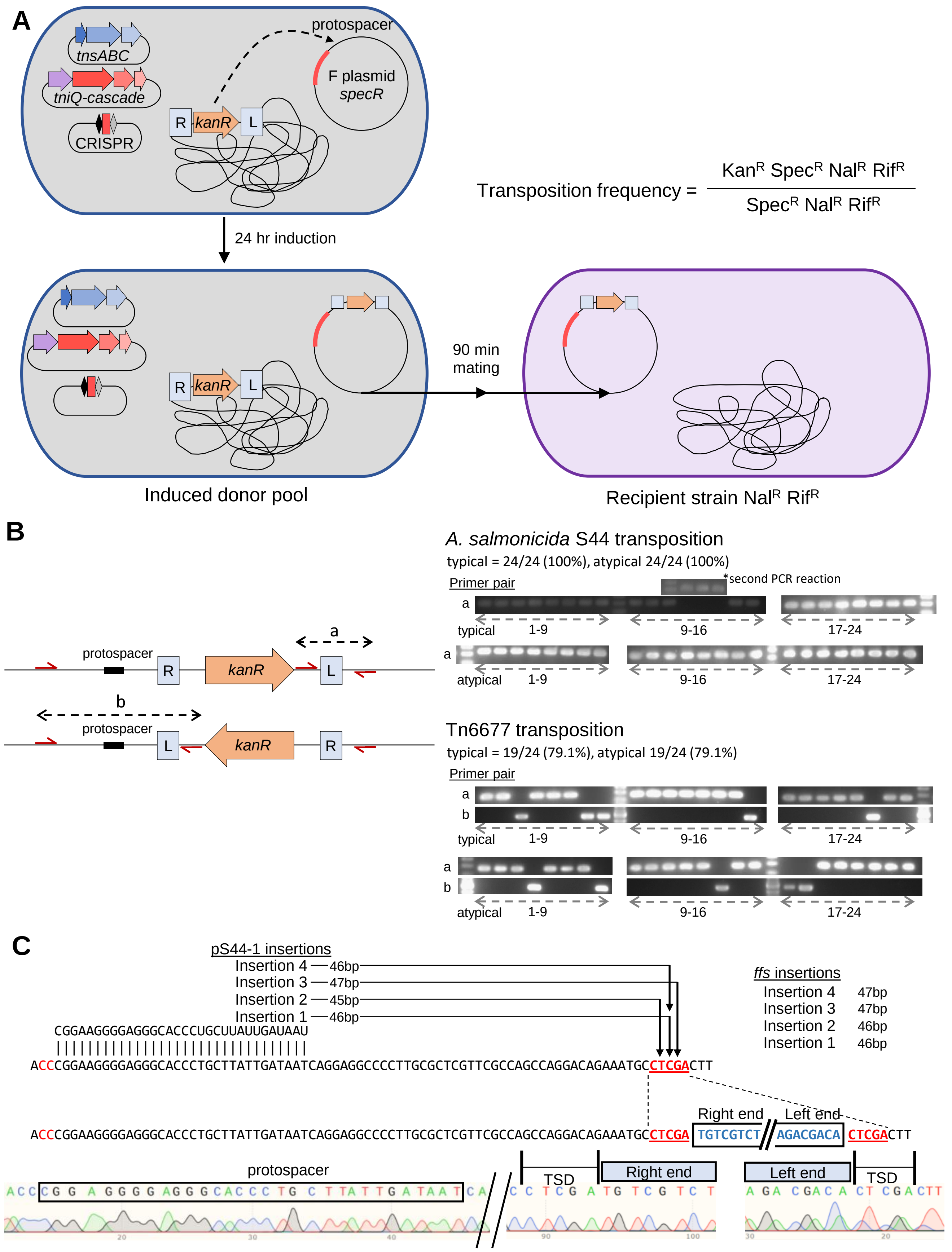

#### Supplemental Figure S3 – Interference assay repeat sequences

*P. aeruginosa* PA14

```
Repeats  - GTTCACTGCCGTGTAGGCAGCTAAGAAA[N32]GTTCACTGCCGTGTAGGCAGCTAAGAAA
```

*A. salmonicida* S44

```
Typical  – GTGAACTGCCGTATAGGCAGCCAAGAAA[N32]GTGAACTGCCGTATAGGCAGCCAAGATT

Atypical – GTGAACTGCCGTATAGGCAGCCAAGATT[N32]ATTTTCTGCCGTAAAGGCAGATATTATT
```

*V. cholerae* Tn6677

```
Typical  – GTGAACTGCCGAGTAGGTAGCTGATAAC[N32]GTGAACTGCCGAGTAGGTAGCTGATAAC

Atypical – TCATTACTACTGCAAAGTAGCTGATAAC[N32]CTTTACTGCTGAATAAGTAGATAACTAC
```

#### Supplemental Figure S4 – I-F3 Xre cluster into two clades with restriction-modification C proteins

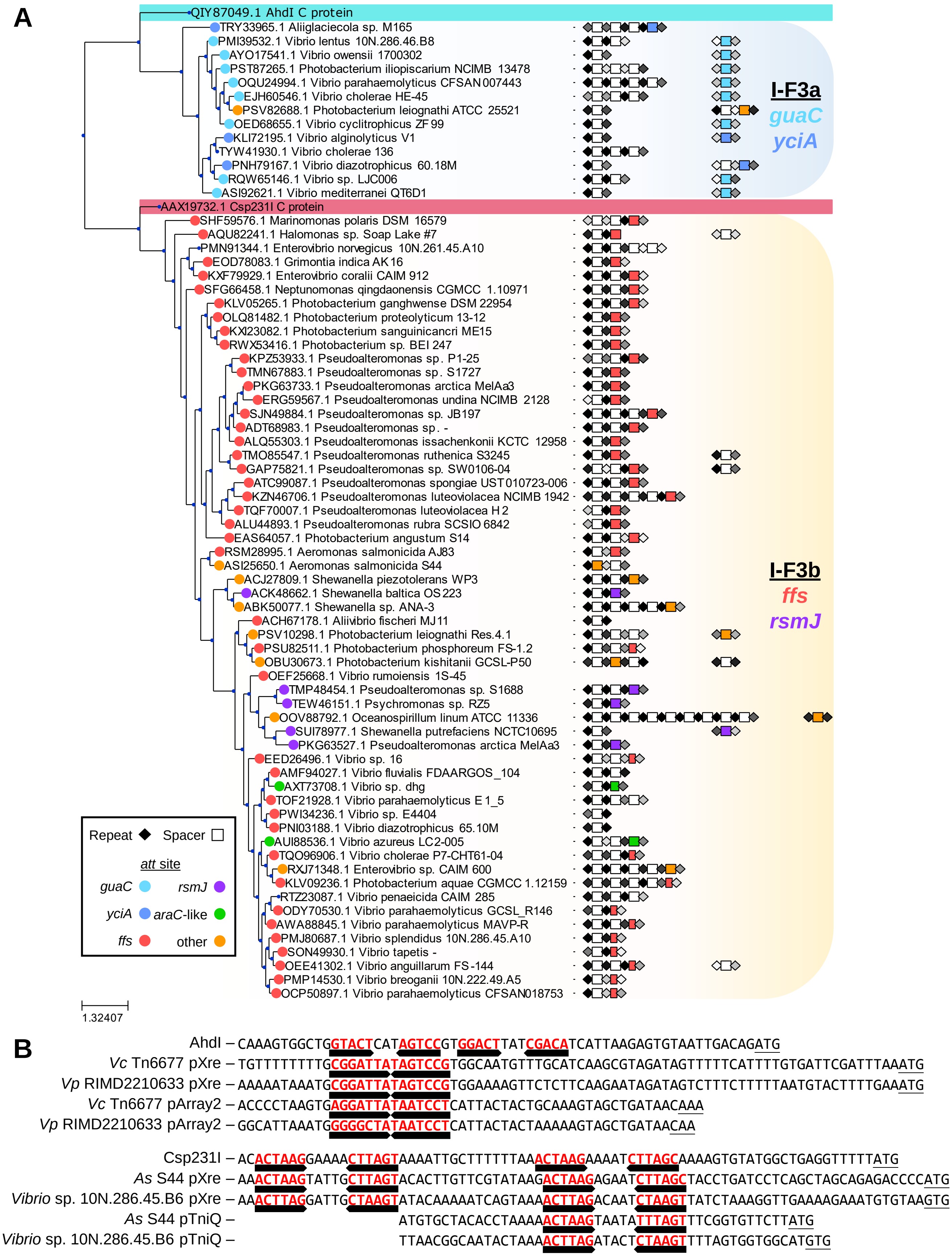

### Supplemental Figure S5 – Elements with shortened spacers and their insertion positions

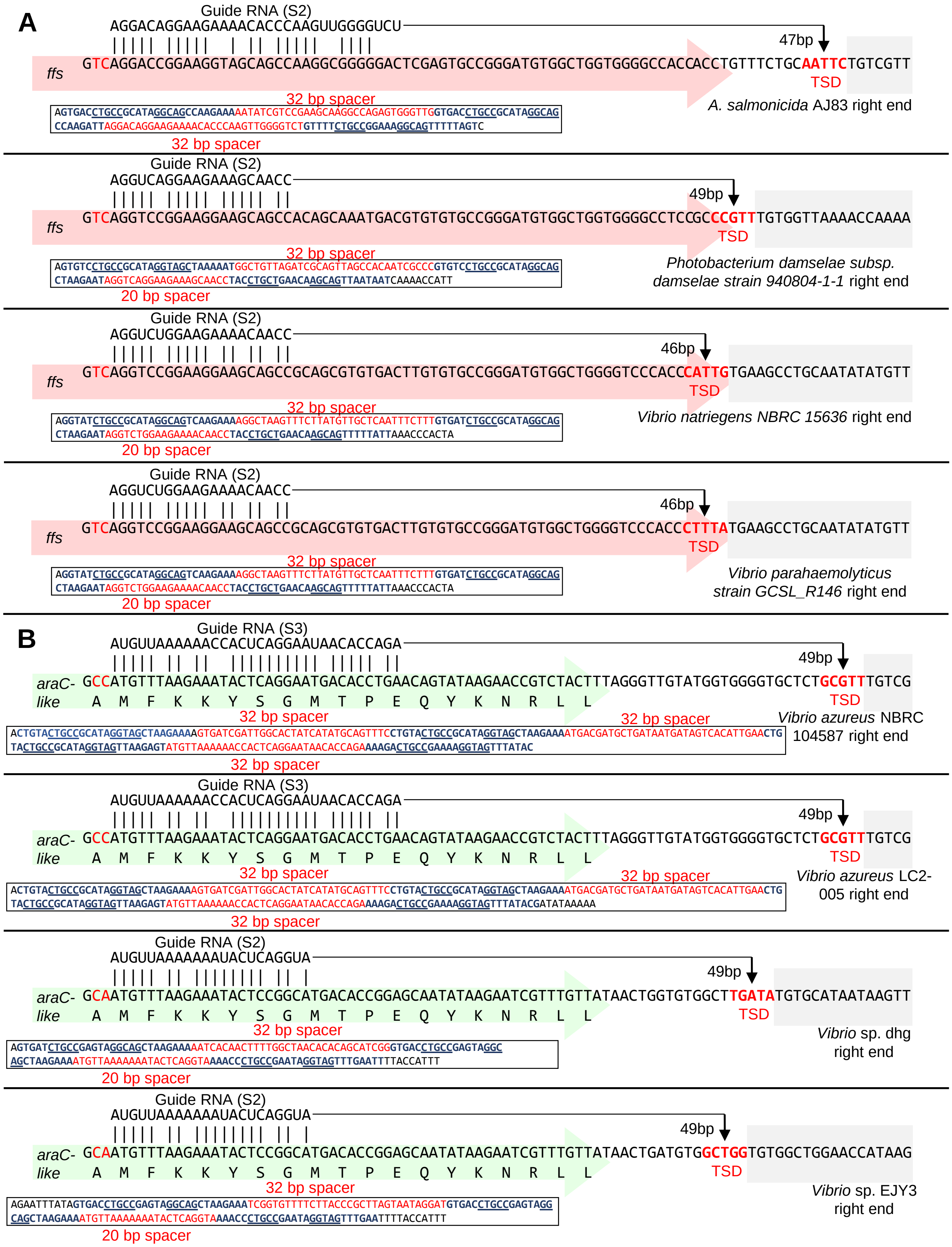

**Supplemental Figure S6 – Schematic representation of elements inserted downstream of *parE***

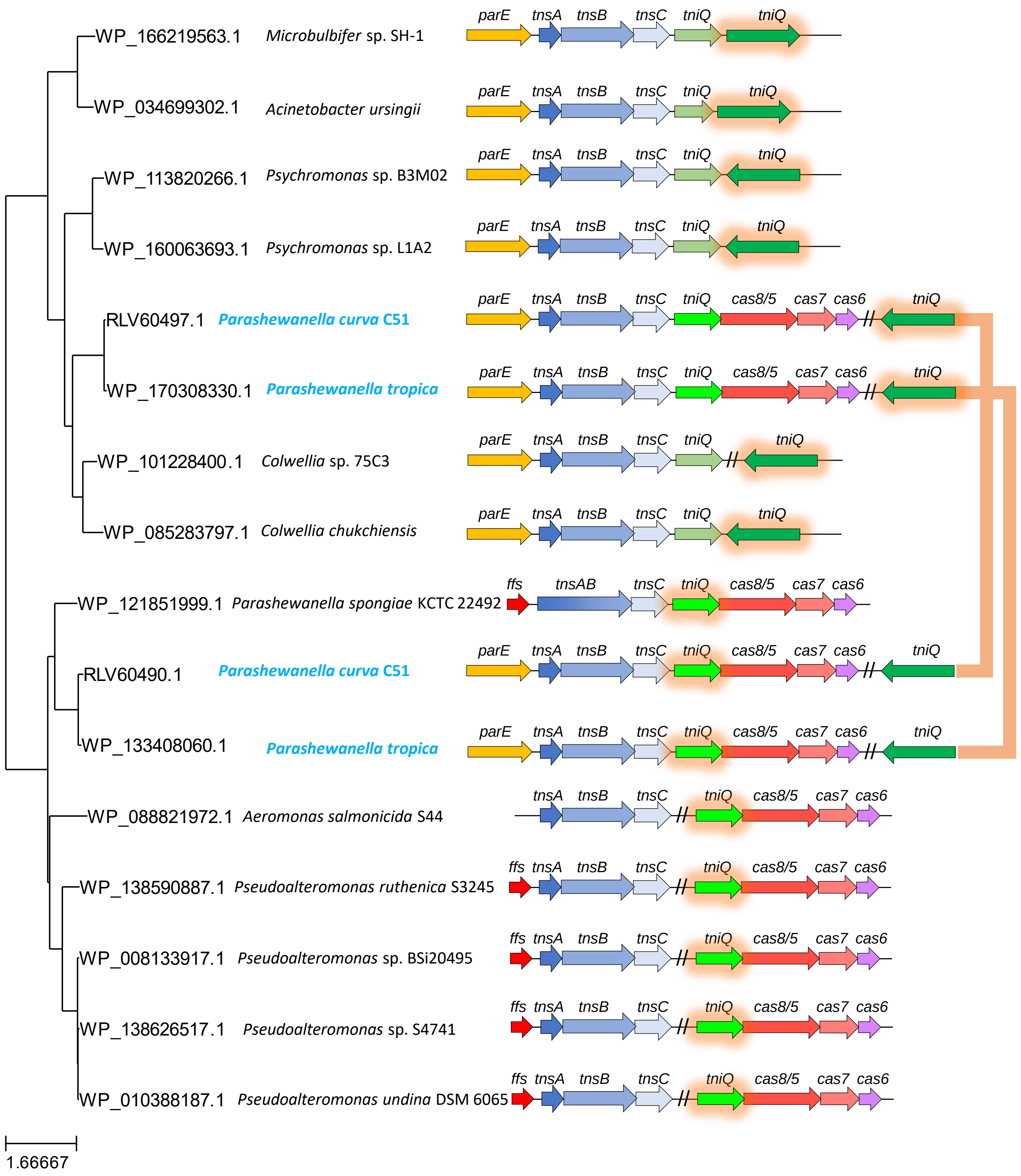
